# Time-resolved fluorescent proteins towards fluorescence microscopy in the temporal and spectral domains

**DOI:** 10.1101/2024.12.16.628778

**Authors:** Zizhu Tan, Chia-Heng Hsiung, Jiahui Feng, Yangye Zhang, Junlin Chen, Ke Sun, Peilong Lu, Jianyang Zang, Wenxing Yang, Ya Gao, Jiabin Yin, Tong Zhu, Yuxuan Ye, Yihan Wan, Xin Zhang

**Author notes:** These authors contributed equally (Zizhu Tan, Chia-Heng Hsiung, Jiahui Feng). Corresponding authors: Xin Zhang.

## Abstract

Fluorescence microscopy has been widely applied in life sciences. While steady-state fluorescence intensity is mostly used, the time-resolved detection mode relying on fluorescence lifetime remains underexplored. Herein, we present a family of time-resolved fluorescent proteins (TRFP). Using a strategy that regulates lifetime without affecting spectra of FPs, we have developed different TRFPs that cover the visible spectrum and a wide range of lifetime. TRFPs are employed in the temporal-spectral resolved microscopy, allowing the simultaneous imaging of 9 different proteins in live cells and correlation of multiple activities to cell cycles. Finally, TRFPs are integrated into multiplexing super-resolution microscopy to concurrently visualize 4 proteins in live cells, acquiring a remarkable 50 nm spatial resolution using lifetime as signal. TRFPs represent a transformative toolset towards fluorescence microscopy in both the temporal and spectral domains. Beyond the multiplexing imaging, TRFPs could introduce unprecedented opportunities of system complexity and quantitative accuracy into biological research.

## INTRODUCTION

Fluorescence microscopy has been widely applied in all areas of life sciences (Lu et al., 2023; Tian et al., 2022; Wolff et al., 2023; Yu et al., 2015). The most widespread fluorescent detection mode is steady-state fluorescence, as commonly referred as fluorescence intensity. The second detection mode is time-resolved fluorescence that is monitored as a function of time upon excitation, akin the lifetime of excited state or fluorescence lifetime (Datta et al., 2020). Until now, most of the fluorophores or fluorescent proteins are developed with a focus on optimizing photophysical properties centering the steady-state fluorescence intensity (Chudakov et al., 2003; Heim et al., 1994; Matz et al., 1999). By contrast, much less efforts have been invested to uncover the mechanisms or approaches that regulate fluorescence lifetime (Frei et al., 2022).

Taking fluorescent proteins (FP) as an example, the broad family of FP derivatives provide with a variety of spectroscopic properties for steady-state fluorescence microscopy, including excitation and emission spectrum, photostability, brightness, photo-switching property, and others (Chudakov et al., 2004; Filonov et al., 2011; Pédelacq et al., 2006; Rizzo et al., 2004; Yu et al., 2014). However, very few studies have been documented in the literature to systemically regulate fluorescence lifetime of FPs and develop time-resolved microscopic approaches towards a wide range of biological applications. On one hand, the few reported lifetime investigations had been purposed as a screening strategy for increase of quantum yields (Bindels et al., 2017; Goedhart et al., 2010), a steady-state property of fluorescence. On the other hand, one of the most common applications of time-resolved fluorescence microscopy is assessing the efficiency of fluorescence resonance energy transfer in live cells (Handlin and Dai, 2023; Lee et al., 2009). Thus, there is a prominent gap that the field needs to fill by systematically regulating the fluorescence lifetime of FPs and shifting the paradigm of time-resolved fluorescence microscopy.

The lack of regulatory exploration of fluorescence lifetime does not reflect its real value in imaging applications. On the contrary, time-resolved fluorescence offers unique advantages that the fluorescence intensity cannot parallel with. For instance, fluorescence lifetime is independent of local concentrations of fluorophores, making it a robust signal without influence from concentration fluctuation (Stringari et al., 2015; Sun et al., 2010; Yaseen et al., 2017). Importantly, time-resolved and steady-state fluorescence exist in the orthogonal temporal and spectral domains. Hence, the combination of these two fluorescent detection modes is expected to generate new applications of biological imaging, exemplified by the emerging technical developments towards multiplexed imaging that plays an important role in revealing organelle interactions and cell signal transduction. Canonical fluorescence microscopy relying on the steady-state detection mode fails to simultaneously visualize more than 6 different proteins, due to the overlap of spectra in the visible region (Valm et al., 2017). While a method named “signaling reporter islands” could distinguish signals with the same excitation and emission spectrum through clustering at different spots in cells, this approach is laborious and technically demanding with the complicated procedures of post-imaging sample processing (Linghu et al., 2020). Most recently, the temporally multiplexed imaging (TMI) technique is reported using the regulated photo-switching kinetics of FPs to achieve multiplexing imaging of 7 different signals in living cells (Qian et al., 2023). However, results from TMI require a 20-70 frame brief movie to unmix 3-6 FPs for each color channel, thus bearing limitations of temporal and spatial resolutions. To overcome these limitations, a complementary and powerful approach is fluorescence lifetime imaging microscopy (FLIM), which offers an attractive opportunity for multiplexed imaging (Niehörster et al., 2016; Scipioni et al., 2021; Starling et al., 2023). However, the only available imaging agent is the rhodamine fluorophore that exhibits different fluorescence lifetime via conjugating with HaloTag variants (Frei et al., 2022), limiting the broad application of the multiplexed imaging.

In this work, we present a new family of FPs, named time-resolved fluorescent proteins (TRFP), developed via protein engineering of residues that are in close vicinity to the electron-donating group of FP chromophores. Remarkably, mutations that result in TRFPs exert minimal effect on spectral properties, with primarily influence on their fluorescence lifetime. Computational and experimental studies collectively revealed that the mutations only affected the excited state structure, but not the ground state, of FP chromophores, providing the underlying mechanisms of TRFP. The strategy was applied to fluorescent proteins in 7 excitation and emission channels, generating an unprecedented toolset of 29 different TRFPs that cover the entire visible spectrum (383-627 nm) and a wide range of fluorescence lifetime (1-5 ns). We applied TRFPs to temporal-spectral resolved microscopy, realizing multiplexing imaging with the simultaneous visualization of 9 different targeting proteins in live cells. Moreover, we converted the traditional Fucci assay into the time resolved Fucci assay that uses only a single spectral channel to monitor cell cycle (Bajar et al., 2016; Sakaue-Sawano et al., 2008), thus leaving other spectral channels to couple activities of CDK2 and CDK4/6 with cell cycles (Spencer et al., 2013; Yang et al., 2020). Finally, we incorporated TRFPs into multiplexing super-resolution microscopy to simultaneously visualize 4 subcellular organelles and cytoskeleton proteins in live cells, with a remarkable spatial resolution as high as 50 nm for microtubule and 110 nm for nuclear pore complex. In summary, TRFP represents a transformative approach to design and engineer fluorescent proteins. Beyond applications demonstrated in this work, we envision that TRFPs could be introduced to FP sensors that detect biological analytes and activities (Greenwald et al., 2018; Wu et al., 2023; Zhao et al., 2011; Zhu et al., 2023), thus offering unprecedented opportunities of system complexity and quantitative accuracy into life sciences.

## RESULTS

### A general strategy to engineer TRFPs

The goal of this work is to combine the steady state and time-resolved fluorescence in microscopy. While traditional fluorescent proteins are differentiated by their steady state properties, such as excitation and/or emission spectra, TRFP should feature differences in the time of proteins spend at their excited states, akin fluorescence lifetime. To enable this technical development, we choose the following three fluorescent proteins, as cyan fluorescent protein (mTurquoise2) (Goedhart et al., 2012) green fluorescent protein (mNeonGreen) (Shaner et al., 2013) and red fluorescent protein (mScarlet) (Bindels et al., 2017). These proteins are commonly used in steady state fluorescence microscopy, exhibiting excitation and emission spectra in the cyan channel (ex = 435 nm and em = 474 nm for mTurquoise2; Figures 1A-1B), the green channel (ex = 506 nm and em = 517 nm for mNeonGreen; Figures 1A-1B), and the red channel (ex = 569 nm and em = 593 nm for mScarlet; Figures 1A-1B).

**Figure 1.**
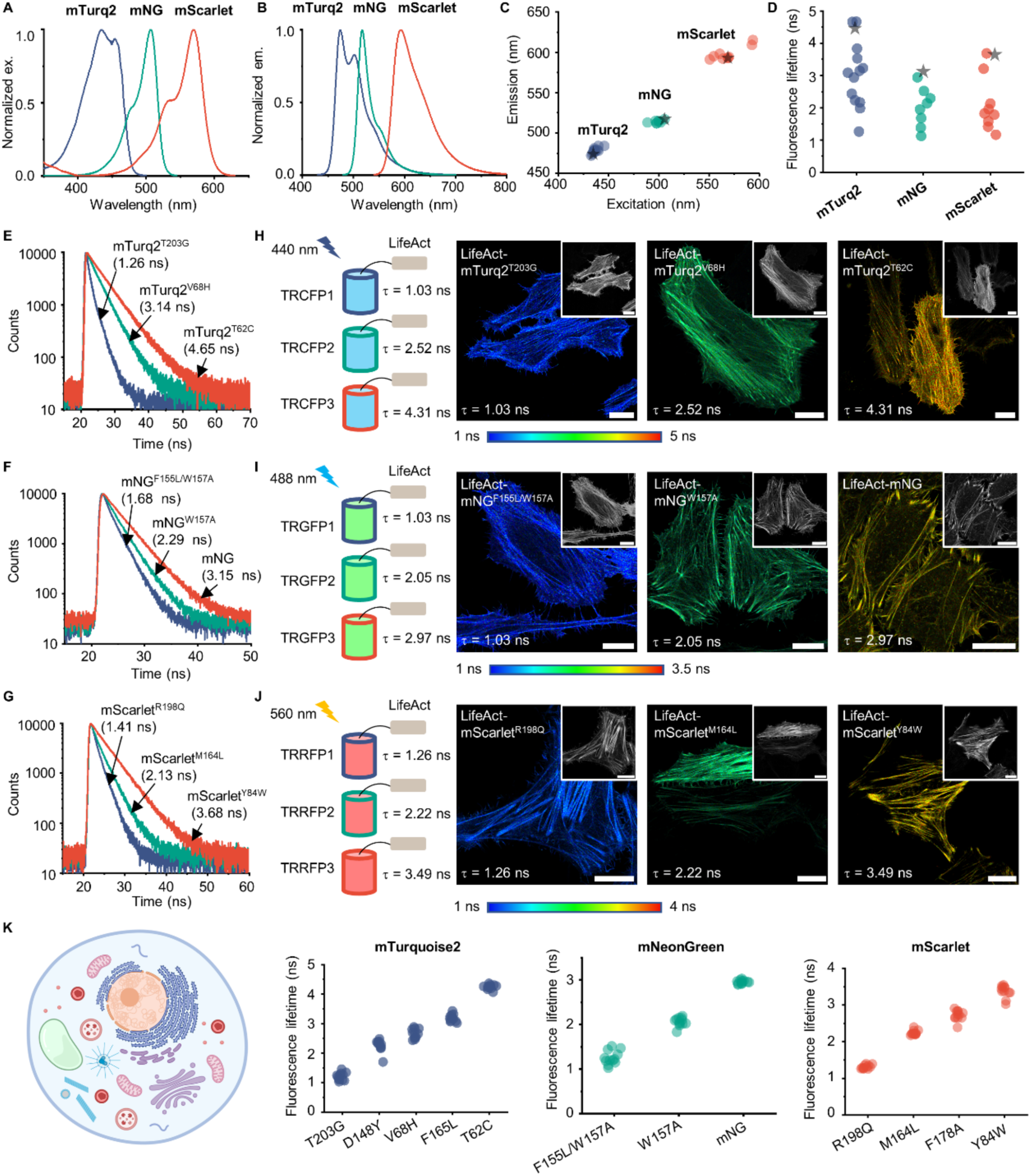
Characteristics of TRFPs derived from mTurquoise2, mNeonGreen and mScarlet. (A) The excitation spectra of mTurquoise2, mNeonGreen and mScarlet in the buffer (20 mM Tris, 500 mM NaCl, pH 7.5). (B) The emission spectra of mTurquoise2, mNeonGreen and mScarlet in the buffer (20 mM Tris, 500 mM NaCl, pH 7.5). (C) The excitation and emission wavelength of different variants derived from mTurquoise2 (blue), mNeonGreen (green) and mScarlet (red) in the buffer (20 mM Tris, 500 mM NaCl, pH 7.5). *indicates the wild type. (D) The fluorescence lifetime of different variants derived from mTurquoise2 (blue), mNeonGreen (green) and mScarlet (red). TRFPs were excited using 10 MHz pulse laser to allow a complete decay and 1×10^4^ photons were collected for each sample. The fluorescence lifetime is the average of three independent measurements. *indicates the wild type. (E) The fluorescence lifetime decay of mTurquoise2^T203G^, mTurquoise2^V68H^ and mTurquoise2^T62C^ in the buffer (20 mM Tris, 500 mM NaCl, pH 7.5). (F) The fluorescence lifetime decay of mNeonGreen^F155L/W157A^, mNeonGreen^W157A^ and mNeonGreen in the buffer (20 mM Tris, 500 mM NaCl, pH 7.5). (G) The fluorescence lifetime decay of mScarlet^R198Q^, mScarlet^M164L^ and mScarlet^Y84W^ in the buffer (20 mM Tris, 500 mM NaCl, pH 7.5). (H) FLIM images of mTurquoise2^T203G^, mTurquoise2^V68H^ and mTurquoise2^T62C^ fusing LifeAct. The inset display the intensity images. The mTurquoise2-based TRFPs are excited by pulsed laser at 440 nm. (I) FLIM images of mNeonGreen^F155L/W157A^, mNeonGreen^W157A^ and mNeonGreen fusing LifeAct. The inset display the intensity images. The mNeonGreen-based TRFPs are excited by pulse laser at 488 nm. (J) FLIM images of mScarlet^R198Q^, mScarlet^M164L^ and mScarlet^Y84W^ fusing LifeAct. The inset display the intensity images. The mScarlet-based TRFPs are excited by the pulse laser at 560 nm. (K) Fluorescence lifetime of mTurquoise2-based, mNeonGreen-based, and mScarlet-based TRFPs fusing to different subcellular-targeted proteins in HeLa cells. The fluorescence lifetime reported is the average of three independent experiments. mTurq2 and mNG are the abbreviation of mTurquoise2 and mNeonGreen, respectively. Scale bar: 20 μm. See also Figures S1-S9.

Our initial goal was to discover their variants that differ only in fluorescence lifetime, but not in excitation and emission spectra. Chromophore of FPs are canonical molecular rotors, whose excited state conformation and rotational motion have been shown to significantly affect emission efficiency of their fluorescence. Thus, we hypothesized that mutations in the close proximity to the FP chromophores would potentially change their excited state lifetime without altering their excitation and emission spectra. To this end, we conducted saturation mutation for amino acids that were within 6 Å distance from chromophores for mTurquoise2 (Figure S1A), mNeonGreen (Figure S1B), and mScarlet (Figure S1C). After a few rounds of mutational cycles, we found that most mutations minimally perturbed the excitation and emission wavelengths (Figure 1C and Tables S1-S3). Interestingly, a collection of TRFP variants was acquired to produce a wide array of fluorescence lifetime (Figure 1D), spanning evenly within the range of 1-5 ns for 13 mutants of mTurquoise2 (Table S1), 1-3 ns for 8 mutants of mNeonGreen (Table S2), and 1-4 ns for 9 mutants of mScarlet (Table S3). To verify the photophysical behavior of TRFP variants, selected proteins were purified and characterized via time-resolved fluorescence spectroscopy through the time-correlated single photon counting (TCSPC). The spectral measurement demonstrated primarily single exponential decay for most of the TRFP variants, as represented by T62C (4.7 ns), V68H (3.1 ns) and T203G (1.3 ns) for mTurquoise2 (Figure 1E), mNG (3.2 ns), W157A (2.3 ns) and F155L/W157A (1.7 ns) for mNeonGreen (Figure 1F), and Y84W (3.7 ns), M164L (2.1 ns) and R198Q (1.4 ns) for mScarlet (Figure 1G).

We next asked whether FLIM microscopy could resolve TRFPs with same spectra and different lifetime. To this end, we firstly chose three mTurquoise2-based TRFPs (T203G, V68H, T62C) with varying lifetimes and genetically fused them to the actin filaments (LifeAct). When TRFP-LifeAct was expressed in HeLa cells, FLIM microscopy was able to separate these proteins via their distinct lifetime (Figure 1H), although the same excitation pulse laser of 440 nm and emission channel of 450-600 nm were used for all samples. By contrast, confocal fluorescence microscopy produced images that were unable to resolve three proteins using the identical excitation and emission channels (inset of Figure 1H). In addition to TRFPs based on mTurquoise2, similar observations were made with TRFPs based on mNeonGreen (F155L/W157A, W157A, mNG; Figure 1I) and mScarlet (R198Q, M164L, Y84W; Figure 1J).

We further investigated whether the TRFP variants exhibit consistent fluorescence lifetime in various live cell imaging conditions. Firstly, we found that fluorescence lifetime was primarily consistent between purified samples and proteins expressed in HEK293T cells, with the averaged difference as 0.18 ns, 0.19 ns, and 0.09 ns for TRFPs based on mTurquoise2, mNeonGreen, and mScarlet, respectively (representative spectra and images in Figure S2, Tables S1-S3). Secondly, we investigated whether expressing TRFPs at varying subcellular localizations would affect their excited state lifetime. To this end, we chose a total of 12 variants (Figure 1K), including five TRFPs for mTurquoise2 (T203G, D148Y, V68H, F165L, T62C), three TRFPs for mNeonGreen (F155L/W157A, W157A, mNG), and four TRFPs for mScarlet (R198Q, M164L, F178Y, Y84W). To express these proteins in different subcellular compartments and membranes in live HeLa cells, we conducted FLIM experiments by expressing the tandem fusion of selected TRFPs to 12 different targeting proteins, including LifeAct (actin filaments), B4GalT1 (golgi apparatus), calnexin (endoplasmic reticulum), FBL (fibrillar center of nucleoli), H2B (nucleus), keratin 18 (intermediate filaments), LAMP1 (lysosome), LMNB1 (nuclear membrane), mitochondrial targeting sequence (mitochondrial), NPM1 (granular compartment of nucleoli), vimentin (intermediate filaments) and α-tubulin (microtubules). Our results show that the fusion of target proteins resulted in negligible changes to the lifetime of TRFPs, in comparison to the lifetime values measured by proteins that were either *in vitro* purified or expressed in HEK293T cells (Figures 1K and S3-S9). Taken together, these results suggest that we have generated a novel set of TRFPs whose excited states can be controlled to span a wide range of lifetime. Moreover, the fluorescence lifetime of TRFPs remains rather constant regardless of their fusion proteins and cellular locations, providing a robust imaging approach wherein targeting proteins can be separated by their distinct lifetime via FLIM microscopy.

### The structural and mechanistic basis of the lifetime regulation for TRFP

We next attempted to understand how mutations in the above FPs affected their lifetime, resulting in a series of TRFPs. Through examining their x-ray crystal structures, we found that the three FPs, albeit harboring chromophores with different electron donating groups (EDG: indole for mTurquoise2 versus phenol for mNeonGreen and mScarlet), followed similar trends in how adjacent residues regulate fluorescence lifetime. We found that the residues within 3.5 Å above the EDG resulted in the most pronounced reduction to the fluorescence lifetime, as represented by T203, R195 and R198 for mTurquoise2, mNeonGreen and mScarlet, respectively (orange boxes in Figures 2A, S10A, S10C; orange residues in Figures 2B, S10B, S10D). The residues that have a secondary effect of the fluorescence lifetime were found within 2-5 Å away from the edge of the EDG, represented by T62, F146, D148, F165, S205 for mTurquoise2, C139, F155, W157 for mNeonGreen, and M164, F178 for mScarlet (blue boxes in Figures 2A, S10A, S10C; blue residues in Figures 2B, S10B, S10D).

**Figure 2.**
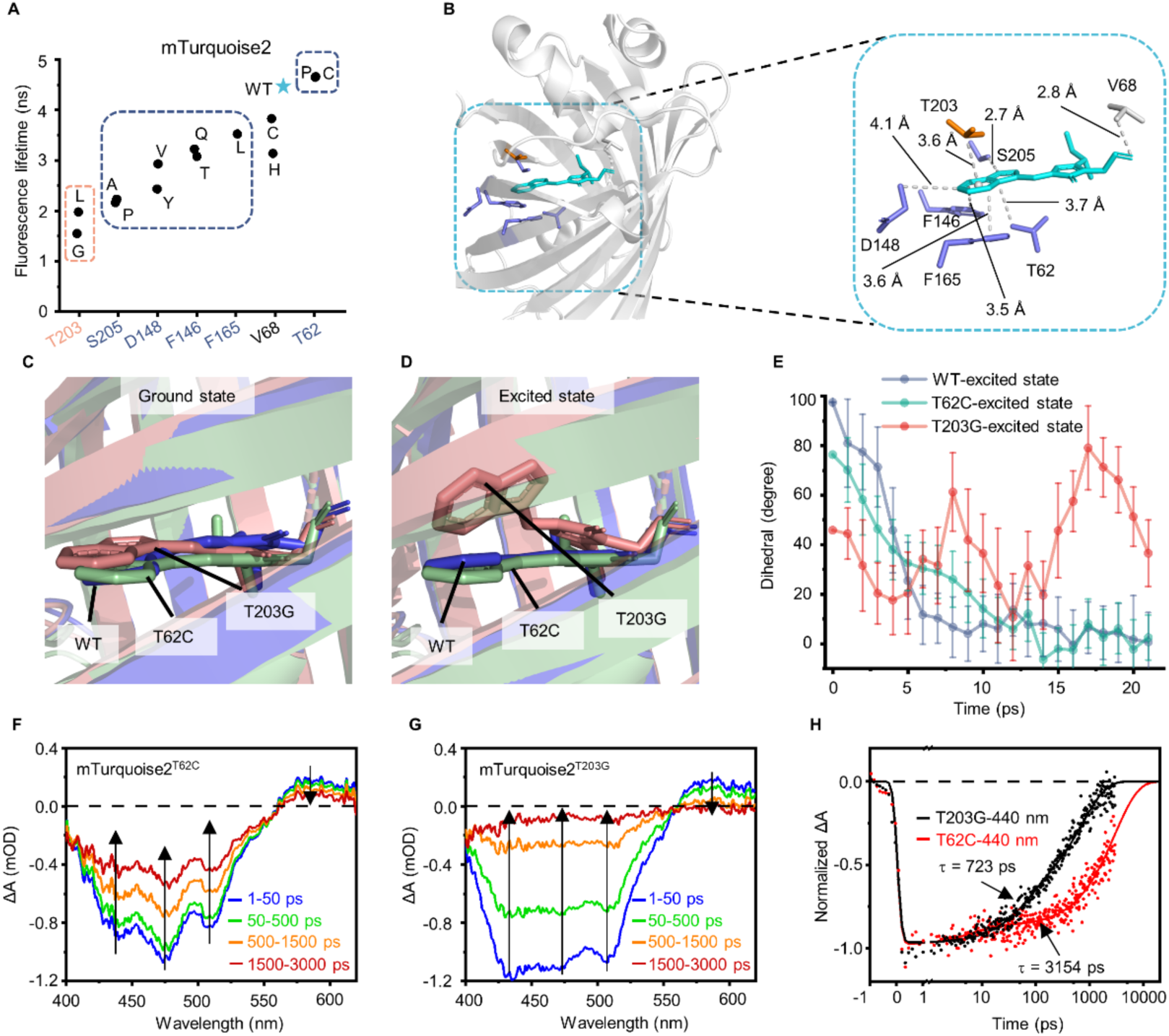
Structural and mechanistic basis of fluorescence lifetime control for mTurquoise2. (A) Effect of different residues replacement on fluorescence lifetime in mTurquoise2. The fluorescence lifetime reported is the average of three independent measurements. *indicates the fluorescence lifetime of mTurquoise2. Orange and blue box indicate the residues which on the top and on the side of indol. (B) Overall structure of mTurquoise2 (left panel) and the distance of the chromophore to the residues that influence the fluorescence lifetime of mTurquoise2 (right panel). The Orange and the blue color indicate the residues which on the top and on the side of indol. The structure of mTurquoise2^K206A^ was used to analysis the structure basis of fluorescence lifetime control for mTurquoise2, PDB ID: 3ztf. (C-E) The structure of wild type (WT) mTurquoise2 (blue) and the variants of T203G (red) and T62C (green) at ground state (C) and excited state (D). Structures were generated by molecular dynamic simulation. (E) The variation of chromophore dihedral angle at excited state with a 20-ps molecular dynamics simulation. (F-H) Femtosecond transient difference absorption spectra recorded at different time delays after a femtosecond laser excitation (370 nm) for mTurquoise2^T62C^ (F) and mTurquoise2^T203G^ (G) in the buffer (50 mM HEPES, 50 mM NaCl, pH 8.0). (H) Kinetic traces of different mTurquoise2 variants at 440 nm and the respective fit based on a global analysis with two exponential functions. In these experiments, the pump-laser energy density was low enough (0.23 mJ cm^-2^) to avoid multiphoton process. See also Figures S10-S15.

To understand the effect of these mutation on the structure of FP, we calculated the structure of these variants through Rosetta. For all three FPs, simulation results showed minimal changes to the backbone and sidechain structures when mutations were introduced (Figures S11, S12, and S13). Consistent to the Rosetta data, a high-resolution quantum mechanism/molecular mechanics (QM/MM) simulation also resulted in the almost identical ground state structure of mTurquoise2, mTurquoise2^T203G^ and mTurquoise2^T62C^ wherein the chromophores were primarily at a planar conformation (Figure 2C), albeit these variants exhibited different fluorescence lifetime (Figure 1E). Considering fluorescence lifetime is a property of the excited state, we next conducted time-dependent density-functional-theory (TD-DFT)/MM simulation to acquire the excited state structure. At excited state, the chromophore of mTurquoise2 and mTurquoise2^T62C^ still adopted a planar conformation, indicating the avoidance of twisted structures that are associated with reduced lifetime. By contrast, the EDG of mTurquoise2^T203G^ at the excited state adopted twisted conformation in relative to the imidazole group of the chromophore (Figure 2D), resulting in a dihedral angle around 40° that most likely contributes to the short lifetime of mTurquoise2^T203G^. To further confirm this notion, we conducted molecular dynamics simulation (MD) to examine the ability of FP chromophores resume planar structure after an initial twisting perturbation. A 20-ps MD simulation revealed that the chromophore for both mTurquoise2 and mTurquoise2^T62C^ was able to resume the planar structure within 10-15 ps (blue and green curves in Figure 2E). However, the chromophore for mTurquoise2^T203G^ was unable to arrive at the planar structure within the duration of simulation, consistent to its short lifetime compared to mTurquoise2 and mTurquoise2^T62C^ (red curve in Figure 2E).

To validate the computational results, femtosecond transient absorption (TA) spectroscopy was utilized to study the photophysical processes of the mTurquoise2^T62C^ (Figure 2F) and mTurquoise2^T203G^ (Figure 2G) samples after photoexcitation at 370 nm. The TA spectra of both samples at the indicated time delay (Figures 2F-2G) include a broad negative signal in the 400 to 565 nm range and a positive photoinduced absorption (PA) band in the 565 to 620 nm range. The former band can be attributed to the overlap of ground-state bleach and stimulated emission (SE), while the PA band corresponds to the excited-state absorption of the samples. The minor spectral shape difference of the negative signal between mTurquoise2^T62C^ and mTurquoise2^T203G^ samples is attributed to the variation of fluorescence quantum yield of the two samples, resulting in varied contributions from SE. Importantly, the TA spectra of mTurquoise2^T203G^ revealed a faster decay than that of mTurquoise2^T62C^ across the entire spectral region. Taking 440 nm for example, multiexponential fittings of the decay kinetics result in amplitude-weighed time constants of 723 ps and 3,154 ps for mTurquoise2^T203G^ and mTurquoise2^T62C^, respectively (Figure 2H). A similar observation was made by comparing the kinetic profiles probed at other wavelengths as well (475, 505, and 580 nm as shown in Figure S14). Taken together, these mechanistic studies suggest that a general strategy to generate TRFP could be mutating residues at the vicinity of EDG to affect the excited state structure of FP chromophores.

### Generating a toolset of TRFPs to cover the entire visible spectrum

To verify whether this strategy is applicable to other FPs, we chose EBFP2 (Ai et al., 2007), mVenus (Kremers et al., 2006), mOrange (Shaner et al., 2004) and mKate2 (Shcherbo et al., 2009), which harbor varying excitation and emission spectral channels to cover the visible region (Figures 3A and 3B). Based on the strategy, we selected the residues above or to the side of EDG for saturation mutagenesis (Figure S15). As we expected, we obtained TRFP variants for each FPs with minimal changes in spectrum (Figure 3C). These TRFPs exhibited a distribution of fluorescence lifetime (Figure 3D), ranging between 2-4 ns for EBFP2 (Figure S16), 2.4-3.2 ns for mVenus (Figure S17), 1.7-3.6 ns for mOrange (Figure S18), and 2-3 ns for mKate2 (Figure S19).

**Figure 3.**
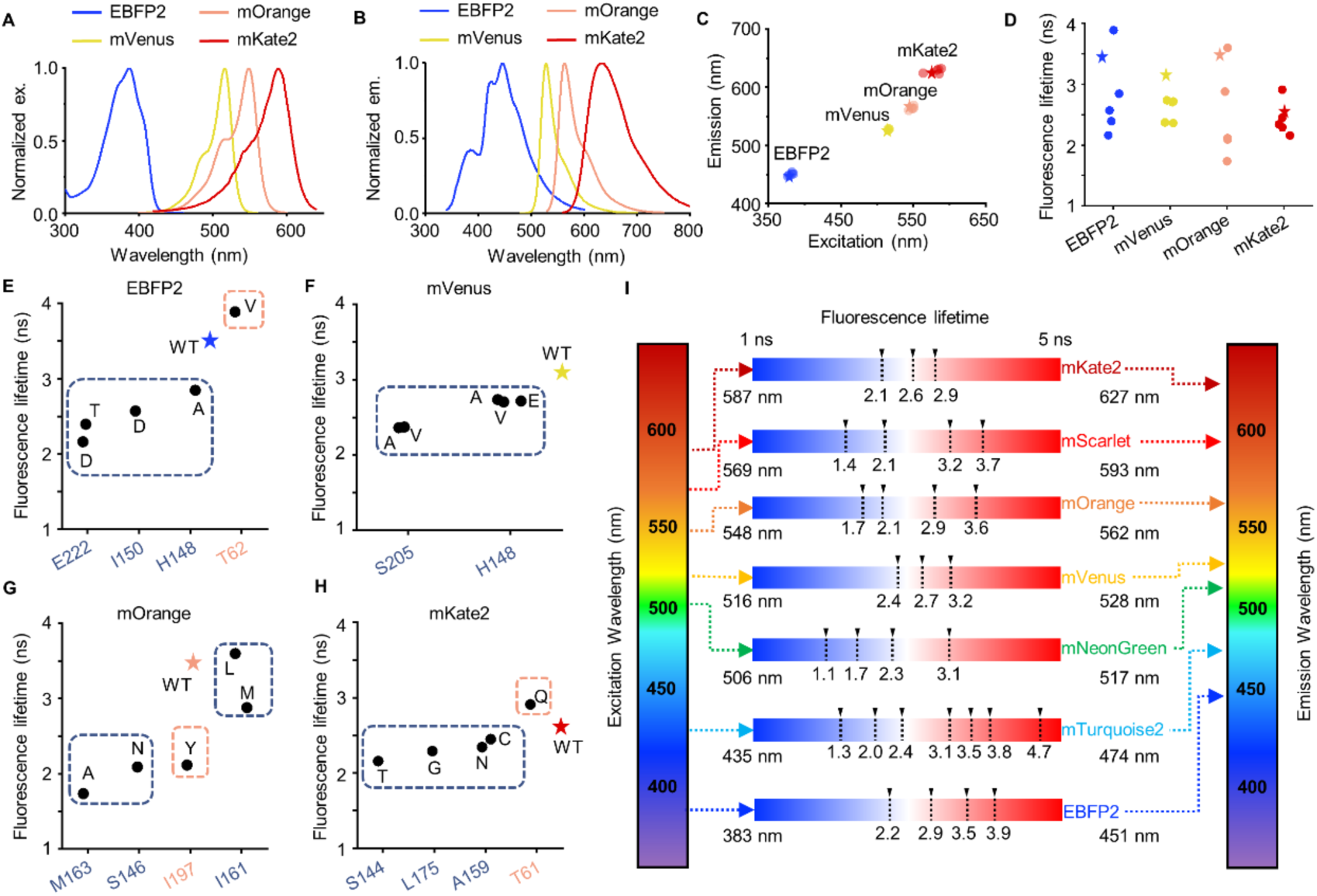
A toolset of TRFPs that cover the entire visible spectrum. (A) The excitation spectra of EBFP2, mVenus, mOrange and mKate2 in the buffer (20 mM Tris, 500 mM NaCl, pH 7.5). (B) The emission spectra of EBFP2, mVenus, mOrange and mKate2 in the buffer (20 mM Tris, 500 mM NaCl, pH 7.5). (C) The excitation and emission wavelength of different variants derived from EBFP2, mVenus, mOrange and mKate2 in the buffer (20 mM Tris, 500 mM NaCl, pH 7.5). *indicates the wild type. (D) The fluorescence lifetime of different variants derived from EBFP2, mVenus, mOrange and mKate2 in the buffer (20 mM Tris, 500 mM NaCl, pH 7.5). The fluorescence lifetime reported is the average of three independent measurements. *indicates the wild type. (E-H) Effect of the different residues on fluorescence lifetime for EBFP2 (E), mVenus (F), mOrange (G) and mKate2 (H) in the buffer (20 mM Tris, 500 mM NaCl, pH 7.5). *indicates the wild type. The Orange and the blue box indicate the residues which on the top and on the side of chromophore. (I) A palette of TRFPs with different fluorescence lifetime and spectra. See also Figures S16-S23.

Structures of all TRFPs were simulated using Rosetta (Figures S20-S23). Consistent to cases of the other three FPs, minimal changes to the ground state conformation of chromophores, indicating excited state structure would be primarily responsible to their different lifetime. Structural analyses indicate that residues at vicinity of the EDG exerted the most pronounced influence on fluorescence lifetime (effect of each residue summarized in Figures 3E-3H). For instance, mutating residues H148 and S205 on the side of EDG for mVenus resulted in a distribution of fluorescence lifetimes between 2-3 ns (Figures 3F, S15B, S17, and S21). In addition, I197 and M163 on the top and edge of EDG for mOrange had significant effect on fluorescence lifetime (Figures 3G, S15C, S18, and S22). The final example is provided by T62 of EBFP2 (Figures S15A and S20) and T61 of mKate2 (Figure S15D), both of which locate on top of EDG. Mutating of both residues increased fluorescence lifetime (Figures 3E, 3H, S16, and S19), indicating mutations of these sites could also prolong the lifetime. Hence, we have successfully generated a rainbow-set of TRFPs, which include a total of 29 different sequences based on EBFP2, mTurquoise2, mNeonGreen, mVenus, mOrange, mScarlet, and mKate2 (Figure 3I and Table S8). These TRFPs cover the entire visible spectrum, spanning from blue to far-red. Moreover, each individual spectral channel of TRFPs is equipped with a temporal resolution that allow us to separate them using FLIM microscopy, providing a powerful tool for novel imaging applications.

### TRFPs enable temporal-spectral resolved multiplexing imaging

TRFPs has the potential to enable new applications due to their temporal resolution via regulated fluorescence lifetime. We next attempted to apply TRFP in FLIM microscopy to realize the temporal and spectral resolved multiplexing imaging. We firstly expressed in HeLa cells a series of TRFP pairs targeting different subcellular locations with the same spectra and fluorescence lifetimes that differ by 0.8-3.2 ns, as represented by LifeAct-mTurquoise2^T203G^ and keratin-mTurquoise2^V68H^ with a 1.8 ns lifetime difference (Figure S24A). Due to their identical spectra, regular confocal microscopy without lifetime information would result in no separation of both proteins. However, FLIM imaging would result in hotspots reflecting lifetime of both TRFPs in the phasor plots acquired at 80 MHz for optimal imaging quality (plot at the right-end of Figure S24A). During the data processing, we positioned the cursor (circular tool) on the phasor plot based on the lifetime of the pure TRFP species. As a result, the two TRFPs could be temporally separated as demonstrated by the two pseudo-colors in the composite image (Figure S24A). In addition to a number of demonstrations using the mTurquoise2-based TRFPs (Figure S24), additional experiments were carried out for mNeonGreen-based TRFPs (Figure S25) and mScarlet-based TRFPs (Figure S26).

Similar strategy can be applied to image cells expressing three TRFPs, as exemplified by co-expression of calnexin-mNG, karetin-mNeonGreen^W157A^, and MTS-mNeonGreen^F155L/W157A^ in HeLa cells (Figure 4A). Although these proteins reside in distinct subcellular localization, it was challenging to discern their identity using confocal microscopy (Figure 4A). By contrast, FLIM would result in phasor plot with triangular hotspots, which could be separated by positioning lifetime of the pure species on the phasor plot (Figure 4B). The assigned pseudo-color to each lifetime could be used to separate three TRFPs in a composite image (Figure 4C). In addition to this case study, we also tested multiple sets of mTurquoise2-based TRFPs (Figure S27), mNeonGreen-based TRFPs (Figure S28), and mScarlet-based TRFPs (Figure S29). In all cases, FLIM microscopy is able to separate TRFPs with different lifetimes, despite of their identical spectra. Furthermore, TRFPs were combined with z-stack scanning to produce three-dimensional images, as demonstrated by co-expression of two TRFPs (MTS-mTurquoise2^T203G^ and keratin-mTurquoise2^V68H^; Figure S30A) and three TRFPs (MTS-mTurquoise2^T203G^, LAMP1- mTurquoise2^V68H^, keratin-mTurquoise2^T62C^; Figure S30B).

**Figure 4.**
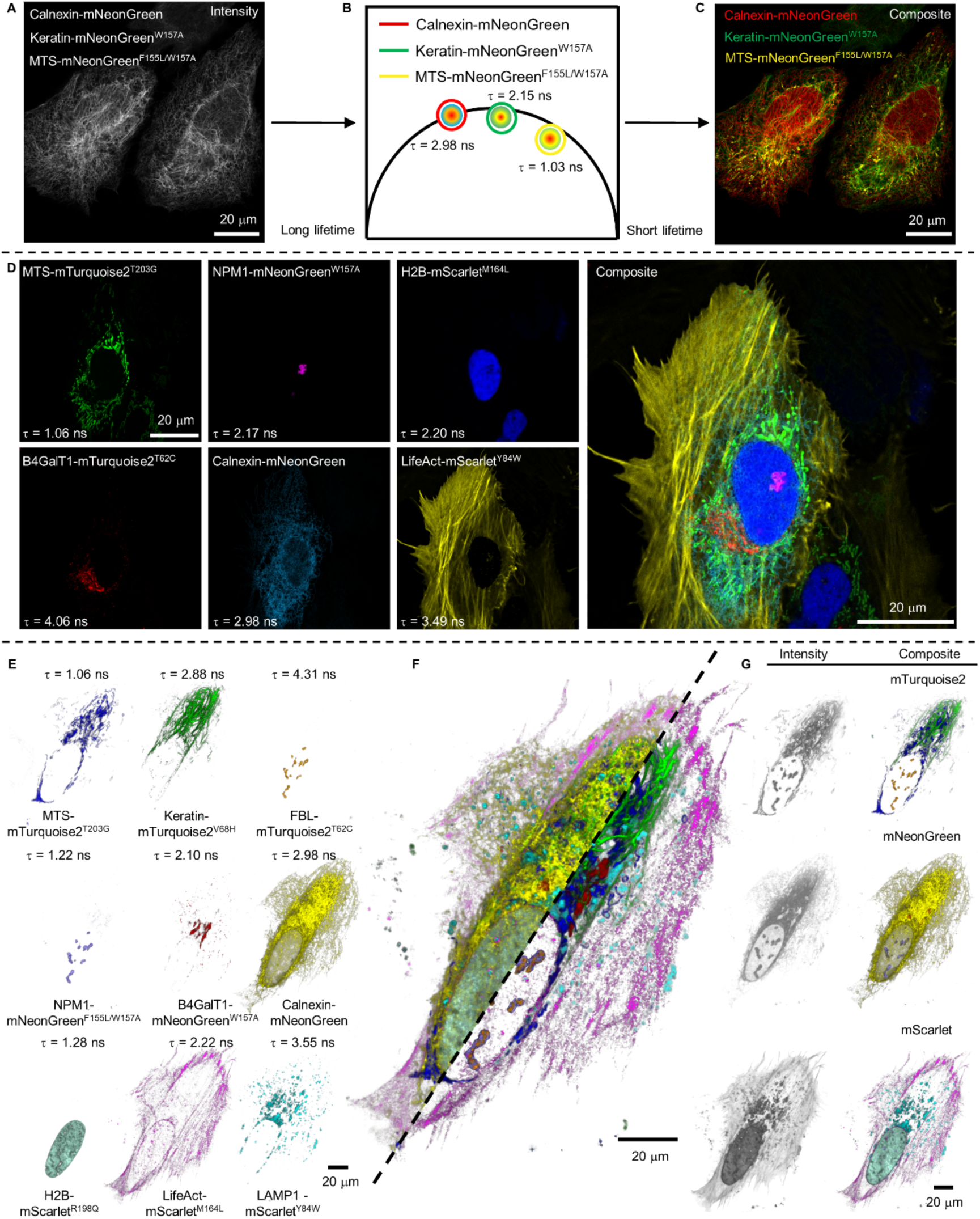
Multiplexing imaging based on fluorescence lifetime and spectra. (A-C) Intensity (A), phasor plot (B) and composite (C) of mNeonGreen, mNeonGreen^W157A^ and mNeonGreen^F155L/W157A^ fusing calnexin targeting endoplasmic reticulum, keratin targeting intermediate filaments and MTS targeting mitochondria in HeLa cells, respectively. (D) Multiplexing imaging of MTS-mTurquoise2^T203G^ targeting mitochondria, B4GalT1-mTurquoise2^T62C^ targeting Golgi apparatus, NPM1-mNeonGreen^W157A^ targeting granular component of nucleolus, Calnexin-mNeonGreen targeting endoplasmic reticulum, H2B-mScarlet^M164L^ targeting nucleoplasm and LifeAct-mScarlet^Y84W^ targeting actin filaments in HeLa cells. (E-G) Multiplexing imaging of MTS-mTurquoise2^T203G^ targeting mitochondria, Keratin-mTurquoise2^V68H^ targeting intermediate filaments, FBL-mTurquoise2^T62C^ targeting fibrillar center of nucleolus, NPM1-mNeonGreen^F155L/W157A^ targeting granular component of nucleolus, B4GalT1-mNeonGreen^W157A^ targeting Golgi apparatus, Calnexin-mNeonGreen targeting endoplasmic reticulum, H2B-mScarlet^R198Q^ targeting nucleoplasm, LifeAct-mScarlet^M164L^ targeting actin filaments and LAMP1-mScarlet^Y84W^ targeting lysosomes in HeLa cells. (E) indicate the subcellar location of each fusion protein. (F) indicate the composite of these nine different target proteins. Right panel show the imaging missing Calnexin-mNeonGreen and H2B-mScarlet^R198Q^. (G) indicate the intensity and composite of cyan (top panel), green (middle panel) and red channel (bottom panel). See also Figures S24-S30.

When the temporal resolution is combined with the spectral resolution, TRFPs could unlock unprecedented opportunities for multiplexing imaging. As a demonstration, we co-expressed six TRFPs genetically fused to different subcellular organelles and cytoskeleton proteins, including MTS-mTurquoise2^T203G^, B4GalT1-mTurquoise2^T62C^, NMP1-mNeonGreen^W157A^, calnexin-mNeonGreen, H2B-mScarlet^M164L^, and LifeAct-mScarlet^Y84W^ (Figure 4D). The FLIM microscopy was carried out in three spectral channels, wherein phasor plot was used to separate lifetime within each channel to separate TRFPs with identical spectral features (three panels to the left of Figure 4D). The resulting composite image provided a clear separation and simultaneous visualization of six TRFPs in the same cell (right panel of Figure 4D), illustrating how these subcellular organelles and actin-filament interact in live cells. Furthermore, we carried out a-stack scanning to provide a comprehensive view of crosstalk between these subcellular compartments in the three-dimensional space (Figure S30C). Finally, we attempted the simultaneous visualization of nine different proteins in live cells. To realize this goal, we co-expressed nine TRFPs in HeLa cells that belonged to three different spectral channels and exhibited different lifetime in each channel, including MTS-mTurquoise2^T203G^, keratin-mTurquoise2^V68H^, FBL-mTurquoise2^T62C^, NMP1-mNeonGreen^F155L/W157A^, B4GalT1-mNeonGreen^W157A^, calnexin-mNeonGreen, H2B-mScarlet^R198Q^, LifeAct-mScarlet^M164L^, and LAMP1-mScarlet^Y84W^. FLIM microscopy not only was able to separate TRFPs with pseudo-color generated for their individual spectra and lifetime features (Figure 4E), but also provide a composite image wherein all nine TRFPs were simultaneously visualized in a life cell (Figure 4F). Given the crowded cellular environment, the chosen subcompartments and cytoskeletons were engaged in an extensive interaction network and arranged to form close contact with each other. After removing calnexin and H2B in the composite image (lower half of Figure 4F), more details of this interaction network were exposed, including the contact of mitochondria with other organelles and the spatial arrangement of NMP1 surrounding FBL in the nucleolus. Such information was only made possible with the combination of temporal and spectral resolution, without which the microscopy would be unable to separate these proteins in a given live cell (compare the regular confocal images with FLIM images in Figure 4G).

### The time-resolved Fucci assay allows quantitative correlation between biological activities and cell cycles

The multiplexing capacity of TRFPs is suited in resolving biological questions that require simultaneous imaging of multiple targets that go beyond the limit of spectral separation. As a demonstration, we attempted to quantitatively study the cyclin-D kinase (CDK) activity during cell cycle. Fluorescent ubiquitination-based cell-cycle indicator (Fucci) has been used to determine the cell cycle through expressing FPs with distinct spectra in different cell cycles (Bajar et al., 2016; Sakaue-Sawano et al., 2008). However, the spectral channels of Fucci FPs would leave limited opportunity to include other FP-based assays in the study of cell cycle. To remove this limitation, we developed time-resolved Fucci (tr-Fucci) assay through incorporating TRFPs into the traditional design. While traditional Fucci biosensors utilized GFP and RFP fusions to the cell cycle-dependent expression and degradation of the hCdt1 and hGem fragments (Sakaue-Sawano et al., 2017), we constructed the tandem fusion of hGem(1/110)-mTurquoise2^T203G^ and hCdt1(1/100)Cy(-)-mTurquoise2^T62C^ (Figure 5A). The tr-Fucci assay would transform the previous detection of cell cycles from emission spectra changes in confocal imaging to fluorescence lifetime changes in FLIM imaging. In particular, the G1 and S phases are marked as lifetime of 4.65 ns and 1.26 ns, respectively; whereas the G2 and M phases should exhibit lifetime in between these two extremes due to the expression of both mTurquoise2^T203G^-hGem(1/110) and mTurquoise2^T62C^-hCdt1(1/100)Cy(-) (Figure 5A). In addition, histone (H1.0) was fused to mTurquoise2^V68H^, whose FLIM images at 3.14 ns could be employed to indicate the change of chromosome to distinguish the G2 and M phases (Figure 5A).

**Figure 5.**
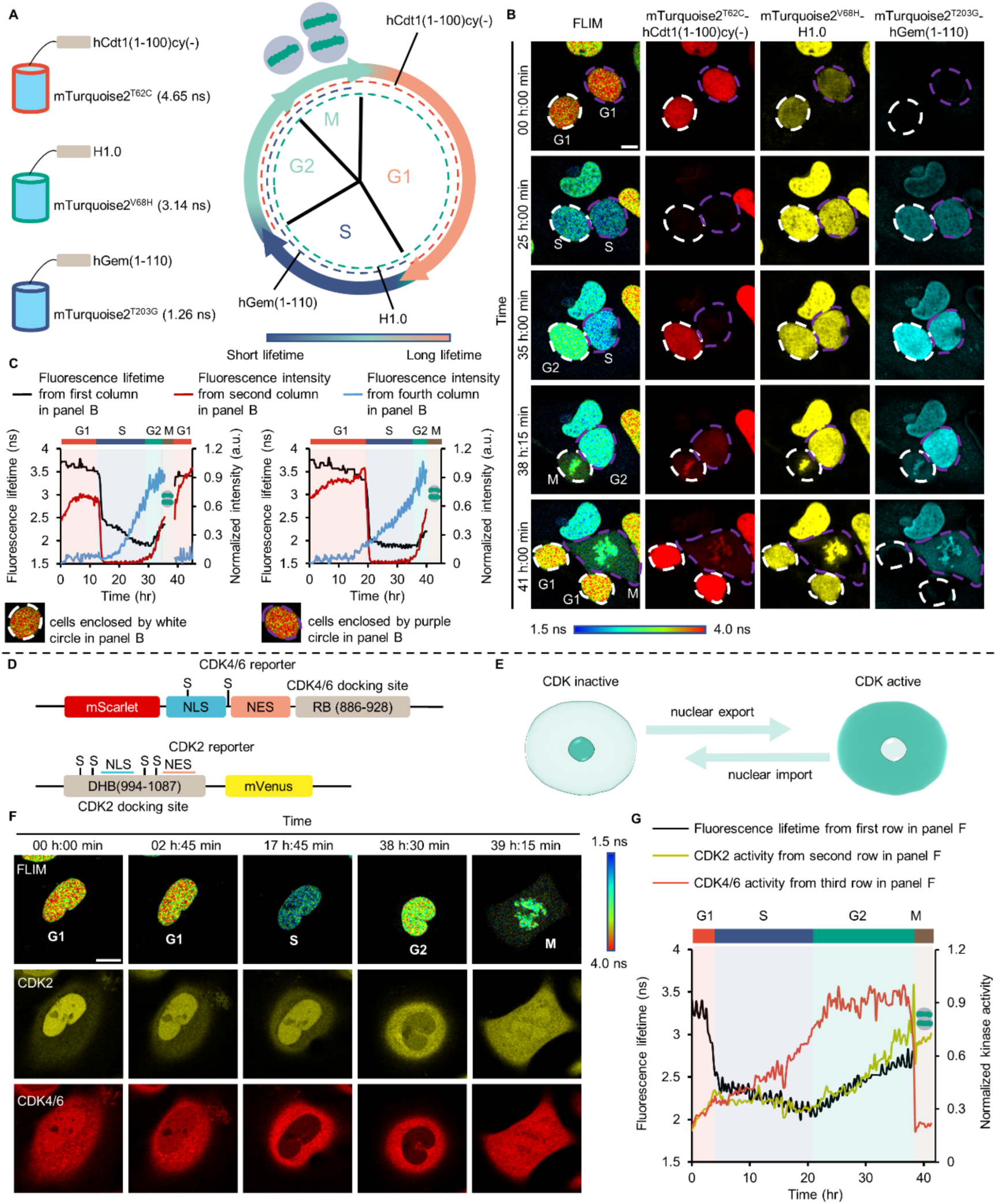
The time-resolved Fucci assay couples CDK activities with cell cycles. (A) Schematic overview of tr-Fucci. During the transition of G1 to S, resulting from the degradation of hCdt1(1-100)cy(-) and the production of hGem(1-110), the fluorescence lifetime decreases. During the transition of S to G2, resulting from the production of hCdt1(1-100)cy(-), the fluorescence lifetime increases. During the transition of G2 to M, H1.0 condenses. During the transition of M to G1, resulting from the degradation of hGem (1-110), the fluorescence lifetime increases. (B) Tracking of cell cycle in HeLa cell via the tr-Fucci. FLIM images were collected every 15 mins. Images of mTurquoise2^T62C^-hCdt1(1-100)cy(-), mTurquoise2^V68H^-H1.0 and mTurquoise2^T203G^-hGem(1-100) were separated through phasor plot. Scale bar, 20 μm (C) The relationship of fluorescence lifetime (black line) with the fluorescence intensity of mTurquoise2^T62C^-hCdt1(1-100)cy(-) (red line) and mTurquoise2^T203G^-hGem(1-110) (blue line). The fluorescence intensity was obtained through phasor plot separated images of column 2 and 4 in (B) and it was calculated by Fiji ImageJ. The icon show the M phase was determined through the condensation of chromatins. Images were collected every 15 mins. Left and right panel represent the cells enclosed by the white and purple circle in (B), respectively. (D) The constructs of CDK4/6 and CDK2 reporter. NLS: nuclear localization signal; NES: nuclear export signal; RB: retinoblastoma protein; DHB: human DNA helicase B. S indicate the phosphorylation site on serine. (E) The activity of CDK is determined by the ratio of fluorescence intensity in the cytoplasm and the nucleus. (F) Tracking of cell cycle and activity of CDK2 and CDK4/6 in live HeLa cell. Images were collected every 15 mins. Scale bar, 20 μm. (G) The activity of CDK2 (yellow line) and CDK4/6 (red line) at different cell cycle in the cell of (F). Black line indicates the fluorescence lifetime which can determine the cell cycle. The icon show the M phase was determined through the condensation of chromatins. Kinase activity was calculated through the ratio of fluorescence intensity in the cytoplasm and nucleus and normalized to the maximum value. Images were collected every 15 mins. See also Figures S31 and S32.

To validate the tr-Fucci assay, we conducted time-lapse FLIM imaging of HeLa cells stably expressing the three TRFP markers, with images of selected time points (Figure 5B) and time traces of two selected single cells (Figure 5C). Two pieces of information was analyzed to monitor the entire cell cycle. First, the apparent fluorescence lifetime within a single nucleus was used to follow the progression of cell cycle (left column of Figure 5B, black curves in Figures 5C and S31). The G1 phase primarily expressed mTurquoise2^T62C^-hCdt1(1/100)Cy(-), resulted in apparent lifetime of 3.6 ns (black curve of Figure 5C and S31). In the transition from G1 to S phase, the degradation of mTurquoise2^T62C^-hCdt1(1/100)Cy(-) and expression of mTurquoise2^T203G^-hGem(1/110) led to a rapid decrease of lifetime to 2.1 ns (black curve of Figure 5C and S31). In the transition from S to G2 phase, expression of mTurquoise2^T62C^-hCdt1(1/100)Cy(-) again resulted in elevation of lifetime (black curve of Figure 5C and S31). After the mitotic phase, the degradation of mTurquoise2^T203G^-hGem(1/110) again increased lifetime increased to 3.6 ns (black curve of Figure 5C). In addition to the apparent fluorescence lifetime, the second information of the tr-Fucci assay is the separation of each individual TRFPs resulted from phasor plots (columns 2-4 of Figure 5B, red and blue curves in Figure 5C and S31). When we followed a single cell, the intensity of phasor separated image of mTurquoise2^T62C^-hCdt1(1/100)Cy(-) and mTurquoise2^T203G^-hGem(1/110) corroborated with the apparent lifetime of individual cells (white circled cell in columns 2 and 4 of Figure 5B, red and blue curves in Figure 5C and S31). Moreover, the morphology of mTurquoise2^V68H^-H1.0 was used to monitor the transition between G2 and M phases, in particular the condensation of H1.0 when cells entered into the M phase (column 3 of Figure 5B). Results from the time-lapse imaging were recorded and presented in movie 1, providing information of both the apparent lifetime and phasor separated TRFPs of tr-Fucci assay. Thus, the tr-Fucci assay could be used to monitor cell cycle based on the lifetime changes. Taking the advantage of only using one spectral channel, tr-Fucci can incorporate other fluorescence-based reporters to couple biomolecular activities and couple them to cell cycles.

To fully demonstrate this advantage, we visualized the activity of CDK2 and CDK4/6 by combining tr-Fucci with their activity reporters that were fused with mVenus and mScarlet FPs, respectively (Figure 5D). For the CDK4/6 reporter ^31^, the 886-928 fragment of retinoblastoma protein (RB) served as CDK-binding domain, which recruited the active CDK4/6 to phosphorylate the serine residues on the NLS and NES domains (top panel of Figure 5D). For the CDK2 reporter ^30^, the 994-1087 fragment of the DNA helicase B (DHB) contained two serine residues as phosphorylation sites adjacent to the NLS sequence and two serine residues next to the NES sequence (bottom panel of Figure 5D). Because phosphorylation enhances the NLS function and dephosphorylation enhances the NES function, the nuclear transport of both reporters could be used as an indicator of the CDK activity (Figure 5E) (Covert et al., 2015). Combined with tr-Fucci, a three-channel FLIM and confocal imaging was conducted to simultaneously couple the CDK2 and CDK4/6 activity with cell cycle, wherein the CDK activity was determined through the ratio of fluorescence intensity in the cytoplasm and nucleus (Figure 5E). Consistent with the previous observations (Coudreuse and Nurse, 2010; Spencer et al., 2013; Yang et al., 2020), CDK4/6 started to elevate its activity during transition from the G1 to S phase (Figure 5F, red curve of Figures 5G and S32). This activity continued to increase during the S phase, reached at the maximal level during the entire G2 phase, and declined when cells started to enter the M phase (Figure 5F, red curve of Figures 5G and S32). Similar to CDK4/6, CDK2 activity also increased during the G1 to S transition (Figure 5F, yellow curve of Figures 5G and S32). Subsequently, CDK2 activity remained steady during the S phase, started to rise in the G2 phase, and exhibited a sharp increase at the entry to the M phase (Figure 5F, yellow curve of Figures 5G and S32). Taken together, we have demonstrated the capacity of TRFP to enable the simultaneous observation of cell cycle, CDK2 and CDK4/6 activity in live cells, showing case the potential of TRFP to provide new insights to biological research.

### TRFPs towards the temporal-spectral resolved super-resolution microscopy

Super-resolution microscopy (SRM) overcomes the light diffraction limit, promoting the revolution of biological research (Betzig et al., 2006; Doksani et al., 2013; Hell, 2003; Moerner and Orrit, 1999). To explore TRFPs that are suitable for SRM, we chose the Stimulated Emission Depletion (STED) microscopy due to its compatibility with FLIM. Because STED achieves super-resolution by reducing effective point spread function (PSF) without post-acquisition calculation, this technology has been combined with FLIM to increase SRM resolution by decreasing background noises and increase the multiplex capacity (Frei et al., 2022; Grotjohann et al., 2011; Pisfil et al., 2022). The requirements for STED compatible FPs are stringent for its brightness, high photostability, and capacity for stimulated depletion. Herein, we adopt a recently reported SIM compatible FP, oxStayGold (oxSG), a variant of a very bright and photostable green fluorescent protein StayGold (Ando et al., 2023; Hirano et al., 2022).

The oxSG FP harbors excitation and emission wavelengths that are similar to GFP, as 488 nm and 504 nm, respectively (Figure 6A). To generate the TRFP variants of oxSG, the residues directly above (K192) and to the side (N137) of the EDG of the chromophore were selected for saturation mutagenesis (Figure 6B). As expected, mutations of K192 reduced the fluorescence lifetime to 1-2 ns (Figures 6C, S33A-C, and S33E-F; Table S9), represented by K192L (2.11 ns, Figure 6D). Compared with K192, mutations of N137 exerted a less pronounced effect of the lifetime for oxSG, resulting in lifetime in the range of 2–3 ns (Figures 6C, S33D, and S33G-I; Table S9) as represented by N137G (3.07 ns, Figure 6D). The Rosetta simulation revealed that the ground state structures of all selected TRFP variants for oxSG were mostly similar to that of the WT-oxSG (Figure S34), suggesting that the regulation of oxSG-based TRFP lifetime was realized at their excited states.

**Figure 6.**
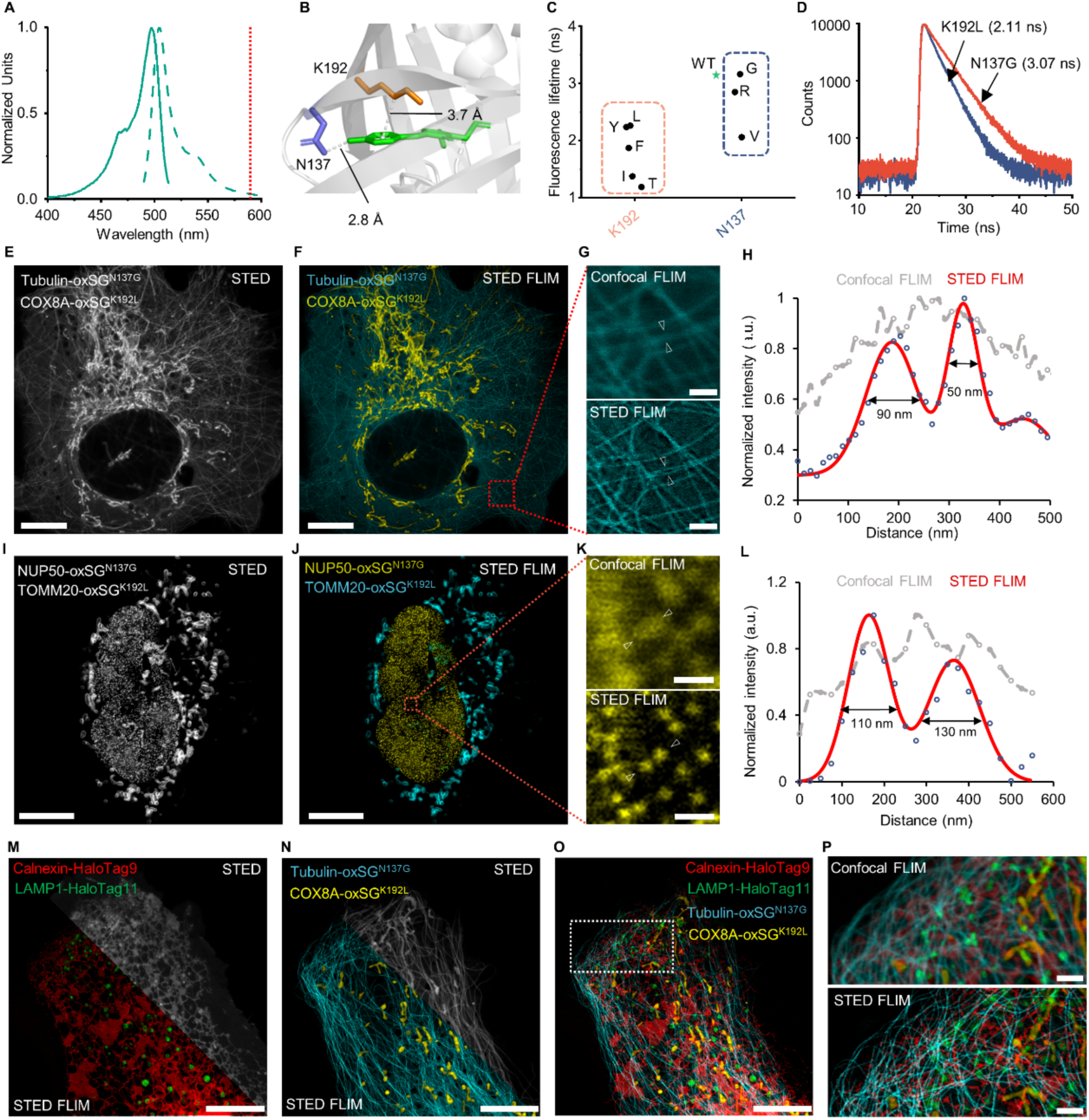
TRFPs enable the temporal and spectra resolved multiplexing imaging with super-resolution. (A) The excitation (solid line) and emission (dash line) spectra of oxSG in the buffer (20 mM Tris, 500 mM NaCl, pH 7.5). Red dash line represents the STED laser beam (592 nm). (B) Residues around the chromophore of oxSG. The orange and the blue indicate the residue which on the top and on the side of chromophore. The structures of StayGold (PDB ID: 8bxt) was used to analysis the fluorescence lifetime control of oxSG. (C) Effect of different residues on fluorescence lifetime for oxSG. The orange and the blue box indicate the residues which on the top and on the side of chromophore. *indicates the wild type. (D) The decay of fluorescence lifetime of oxSG^N137G^ and oxSG^K192L^ in the buffer (20 mM Tris, 500 mM NaCl, pH 7.5). (E-H). STED FLIM of live U-2 OS cells expressing oxSG variants fusing tubulin targeting microtubules and COX8A targeting mitochondria. Intensity (E) and phasor separation (F) of STED imaging of live U-2 OS cells expressing tubulin-oxSG^N137G^ and COX8A-oxSG^K192L^. (G) Enlarged confocal and STED images in (F). (H) line profiles of oxSG^N137G^ fusing tubulin as indicated in (G). (I-L) STED FLIM of live HeLa cells expressing oxSG variants fusing NUP50 targeting nucleus and TOMM20 targeting mitochondria. Intensity (I) and phasor separation (J) of STED imaging of live HeLa cells expressing NUP50-oxSG^N137G^ and TOMM20-oxSG^K192L^. (K) Enlarged confocal and STED images in (J). (L) line profiles of oxSG^N137G^ fusing NUP50 as indicated in (K). (M-P). 2 channels STED FLIM of live U-2 OS cells expressing HaloTag variants fusing calnexin targeting endoplasmic reticulum and LAMP1 targeting lysosomes through labeling Halo-SiR and oxSG variants fusing tubulin targeting microtubules and COX8A targeting mitochondria. (M) Intensity (top panel) and composite (bottom panel) of STED imaging of live U-2 OS cells expressing calnexin-HaloTag9 and LAMP1-HaloTag11. (N) Intensity (top panel) and composite (bottom panel) of STED imaging of live U-2 OS cells expressing Tubulin-oxSG^N137G^ and COX8A-oxSG^K192L^. (O) STED FLIM phasor separation of calnexin-HaloTag9, LAMP1-HaloTag11, tubulin-oxSG^N137G^ and COX8A-oxSG^K192L^. (P) Enlarged confocal and STED images of the white box in (O). oxSG is the abbreviation of oxStayGold. Scale bar, 10 μm (E, F, I, J, M, N and O), 2 μm (P), 1 μm (G and K). See also Figures S33-S35.

To achieve optimal lifetime differences for separation, oxSG^K192L^ (2.11 ns) and oxSG^N137G^ (3.07 ns) were selected for STED-FLIM experiments using the 592 nm STED beam (red line in Figure 6A). FLIM imaging, in the mode of either confocal or STED, was carried out in live U-2 OS cells co-expressing the mitochondria targeting COX8A-oxSG^K192L^ and tubulin-oxSG^N137G^. While regular confocal and STED imaging were unable to separate these proteins due to their identical spectra (Figure 6E), COX8A-oxSG^K192L^ and tubulin-oxSG^N137G^ were separated from each other through phasor plot (Figure 6F). Although the STED beam reduced the overall fluorescence lifetime for both oxSGs from confocal-FLIM (Figures S35B versus S35A; S35D versus S35C), the lifetime differences between oxSG-based TRFPs in STED-FLIM were still sufficient to separate both proteins (Figures S35B and S35D). Importantly, when comparing confocal-FLIM and STED-FLIM, an exciting improvement of resolution for tubulin-oxSG^N137G^ was observed. The adjacent microtubules were unable to be resolved and merged into a mixed set of pixels (top panel of Figure 6G), due to the insufficient resolution of confocal-FLIM (grey curve in Figure 6H). By a sharp contrast, the STED-FLIM resulted in a much-improved spatial resolution to ∼ 50 nm (red curve in Figure 6H), thus clearly resolving two adjacent microtubules that were mixed together in the confocal-FLIM settings (bottom panel in Figure 6G). Similar observations were made when NUP50-oxSG^N137G^ and TOMM20-oxSG^K192L^ were imaged in confocal and STED-FLIM settings (Figures 6I-K). While NUP50-oxSG^N137G^ was unambiguously resolved with ∼110 nm spatial resolution using STED-FLIM (bottom panel in Figure 6K, red curve in Figure 6L), images acquired with confocal-FLIM was unable to discern existence of individual NUP50 due to the low resolution (top panel in Figure 6K, grey curve in Figure 6L).

These results encouraged us to further explore whether we could achieve super-resolution imaging via temporal and spectral-resolved STED-FLIM experiments. To this end, we adopted the recently reported HaloTag variants specifically for fluorescence lifetime multiplexing. In this system, HaloTag was mutated at the area of fluorescent ligand binding, resulting in differential fluorescence lifetime of the Si-Rhodamine (SiR) ligand. Because the spectra of oxSG is orthogonal to that of SiR, we decided to combine oxSG and HaloTag-SiR to enable the temporal-spectral resolved super-resolution microscopy. To this end, we chose the previously reported HaloTag9 and HaloTag11 variants, which could react with to Halo-SiR ligand to produce HaloTag-SiR conjugates with lifetime of 3.43 ns and 2.22 ns, respectively. To fulfill multi-color STED-FLIM, HaloTag9-Calnexin and HaloTag11-LAMP1 fusion proteins labeled with SiR were co-express along with oxSGs in live U-2 OS cells. In each spectral channel, phasor separation was capable of separating HaloTag9-Calnexin from HaloTag11-LAMP1 (bottom half of Figure 6M) and tubulin-oxSG^N137G^ from COX8A-oxSG^K192L^ (bottom half of Figure 6N) although the regular STED imaging was unable to resolve these proteins (top half of Figures 6M-N). In a fully composite image, all four proteins were unambiguously separated via pseudo-color that was dependent on their spectra and lifetime information (Figure 6O). It was noted that STED-FLIM exhibited much improved spatial resolution than the confocal-FLIM technique (Figure 6P), demonstrating the power of TRFPs in the multiplexing temporal-spectral resolved super-resolution imaging.

## DISCUSSION

In summary, we herein present the development and application of time-resolved fluorescent proteins, a new family of FPs that potentiate novel microscopic applications and bring unprecedented biological insights on important questions. We found that the development of TRFP could be achieved via the modulation of residues that are adjacent to the electron-donating group of FP chromophores. Structural and computational studies reveal that these mutations exert minimal changes to the ground state structure of FP chromophore; instead, the excited state structure is significantly modulated within a range from the planar to tilted conformations. These conformational changes in the excited states contribute mostly to the varying fluorescence lifetime, as validated by ultrafast transient absorption spectroscopy. Furthermore, the strategy can be generally applied to most available fluorescent proteins in different emission spectra windows. As a result, we present in this wok a matrix of TRFPs that cover the entire visible spectrum (ex: 383 to 587 nm; em: 451 nm to 627 nm) and a wide range of excited state lifetime (1.1 ns to 4.7 ns). In a foreseeable future, we envision that an expanding number of TRFPs can be developed following this strategy, thus adding a new subgroup to the important family of fluorescent proteins.

The development of TRFPs enables a variety of unprecedented applications. In this work, we demonstrate the temporal-spectral multiplexing imaging technology based on the spectrum domain and time domain through the regulation of FPs fluorescence lifetime. In the past, fluorescence lifetime of FPs was mainly regarded as a screening criterion for increases in quantum yields or was used to calculate the efficiency of FRET (Bindels et al., 2017; Goedhart et al., 2010; Lee et al., 2009). Here, we leverage this feature with the modern FLIM technology, presenting multiplexing imaging with as many as 9 different targeting proteins in live cells. To the best of our knowledge, this is by far the greatest extent of multiplexed imaging in live cells. Using this imaging approach, we also develop the tr-Fucci assay that uses only a single spectra channel, thus leaving sufficient number of channels to monitor activities of biological pathways in cell cycles. Our demonstration has already gone beyond the maximal number of multiplexing reported in literature, including the recently reported temporally multiplexed imaging method utilizing the properties of photo-switching FPs (Qian et al., 2023). Technically, TRFP-based temporal-spectral multiplexing imaging should have uncapped capacity if unlimited number of FP-fusion target proteins can be expressed in live cells.

The temporal-spectral multiplexing imaging of TRFP is primarily achieved via FLIM technology (Ogikubo et al., 2011; Okabe et al., 2012). There is a persistent misperception that FLIM microscopy requires expensive instrumental modules. With the technologically advancement, the reality is that FLIM imaging only needs a pulsed laser source and single-photon detector, both of which are becoming standard of mainstream microscopes. Instrumental wise, a single frequency pulsed-laser source will serve the purpose for time-resolved multiplexing. Because fluorescence lifetime of most TRFPs ranged from 1.0 – 4.0 ns, a repetition rate of 80 MHz (12.5 ns between each pulse) is sufficed for the purpose. Moreover, single-photon detectors have become a common option for conventional confocal microscopes due to the demand in high quality imaging for faster, more sensitive, and wider dynamic range in fluorescence detection, including HyD (Leica microsystems) and SilVIR (Olympus) detectors. To demonstrate this notion, we carried out multiplexing of TRFPs based on mNeonGreen (mNG^F155L/W157A^ and mNG) and mScarlet (mScarlet^R198Q^, mScarlet^M164L^, and mScarlet^Y84W^) on a regular Leica Stellaris 5 confocal microscope equipped with pulsed laser and HyD S detector (Figure S36). The lifetime differences between each variant, around 1.0 ns, was sufficient to differentiate each signal during acquisition using TauSeparation mode – a standard acquisition mode that enables the fluorescence signal separation based on lifetime. With continuously instrumental advancement, capacity of FLIM imaging is expected to become a regular setup in most commercially available microscopes.

In addition to the multiplexing capacity, super-resolution microscopy is another exciting aspect that TRFPs could achieve in this study. In a recent pioneering work, the Boyden group utilizes the switch-off kinetics to differentiate multiple proteins within the same emission channel (Qian et al., 2023). However, this technique has not been shown with the capacity to realize high spatial resolution. For instance, the requirement of a brief movie to record photo-switching kinetics is not compatible with the available techniques for super-resolution microscopy. By contrast, TRFPs provides a brand-new approach for live-cell imaging and offers a whole suite of labeling permutation in emission spectra and fluorescence lifetimes, further, with super-resolution capability. In this work, we developed TRFPs based on oxStayGold (oxSG) FP towards STED-FLIM super-resolution microscopy. In live cell imaging experiments, we acquired a remarkable 50 nm resolution for tubulin-oxSG^N137G^, one of the highest spatial resolutions for FLIM microscopy in living cells. For TRFP variant NUP50-oxSG^N137G^, we also acquired a 110 nm resolution that allowed us to unambitiously separate individual nuclear pore complexes, suggesting that super-resolution could be accomplished by oxSG-based TRFPs. With future development of new STED-compatible FPs, we envision that more varieties of TRFPs would emerge to bring the super-resolution microscopy into the era of temporal-spectral multiplex.

In summary, TRFP represents a novel approach to design and engineer fluorescent proteins, by establishing a general chemical approach that could systematically regulate the excited state lifetime of a wide range of fluorescent proteins. Using this approach, future TRFPs could be developed to potentiate new applications and provide valuable insights. Herein, we demonstrate the temporal-spectral resolved microscopy to realize both multiplexing imaging for up to 9 targets and super-resolution imaging for up to 50 nm, both in live cell samples. However, we believe these cases are only a tip of the iceberg of the application potential for TRFPs. For instance, various FP-based sensors have been developed to report on the existence and activity of biomolecules (e.g., the detection of Ca^2+^, cAMP and cGMP (Lee et al., 2020; Qian et al., 2019; Wang et al., 2022)), mostly replying on fluorescence intensity changes as readout. Because intensity is a concentration-dependent quantity, these FP sensors fall short in providing quantitative measures in live cells or organisms. By contrast, fluorescence lifetime is concentration-independent, thus providing quantitative readout for the biological events under analysis. Thus, we envision TRFP as a transformative technology that have the potential to bring new layers of comprehension and quantification into biological and life sciences.

## ACKNOWLEDGEMENTS

The authors thank Instrumentation and Service Center for Molecular Sciences at Westlake University for the assistance on fluorescence lifetime imaging microscopy. X.Z., P.L.L., Y.X.Y., and Y.H.W. thank support from Research Center for Industries of the Future (RCIF) at Westlake University.

## METHODS

### Plasmids construction

Genes encoding TRFPs and a C-terminal polyhistidine (6xHis) tag were subcloned into pET-28a (+) vector (Novagen) to allow Ni-NTA purification. pCDNA3.1(+) was used as transient expression vector in mammalian cells. H2B (NM_021058.3), LMNB1 (BC012295.1) and α-tubulin (NM_006082) were inserted into the C-terminus of TRFPs. B4GalT1 (NM_001497.3), calnexin (NM_001746.3), FBL (NM_001436.3), keratin 18 (NM_000224.2), LAMP1 (NM_005561.3), NPM1 (NM_002520.5), TOMM20 (NM_014765.2), NUP50 (NM_007172.3), COX8A (NM_004074.2), β-tubulin (NM_178014.2) and vimentin (NM_003380.3) were inserted into the N-terminus of TRFPs. A 20 amino-acid (GGGGS)_4_ linker was used to bridge all the genes and the TRFPs, except MTS, the first 29 amino acid of human cytochrome c oxidase subunit 8A, was insert directly to the C-terminus of TRFPs, and LifeAct has a (GDPPVAT) linker connecting to the C-terminus of TRFPs.

A polycistronic expression vector was engineered based on pcDNA3.1(+) background using two different IRES to separate mTurquoise 2^T203G^, mTurquoise2^V68H^, and mTurquoise2^T62C^. To avoid plasmid amplification errors due to highly repetitive sequences of the three TRFPs. mTurquoise2^V68H^ and mTurquoise2^T62C^ were codon optimized and synthesized by GENEWIZ. The Fucci sensor were then subcloned into the 3’-end of each mTurquoise2 TRFPs resulting: mTurquoise2^T203G^-hGem(1–110)-IRES1-mTurquoise2^T62C^-hCdt1(1–100)Cy(-)-IRES2-H1.0-mTurquoise2^V68H^ (tr-Fucci). tr-Fucci was further subcloned into XLone vector for stable cells construction.

The CDK2 reporter was constructred through fusing partial of human DNA helicase B (aa 994-1087) to the N-termus of mVenus. CDK4/6 reporter was constructed by fusing a CDK4/6-specific docking site (originated from retinoblastoma protein, AA 886-928) to the C-terminus of mScarlet with a linker that contains a 5’-NLS -NES-3’ sequence. Both CDK2 and CDK4/6 reporter were then subclone into a pCDH vector with a P2A separating the two reports for stable cells construction.

The plasmids containing EBFP2, mTurquoise2, mVenus and mKate2 encoding sequences were gifts from Prof. Kiryl Piatkevich (Westlake University). The gene encoding mNeonGreen was a gift from Prof Yongdeng Zhang (Westlake University). The gene encoding oxStayGold, mOrange, CDK2 reporter and CDK4/6 reporter were synthesized from GENEWIZ. The gene encoding NPM1, FBL, TOMM20, β-tubulin, NUP50, B4GalT1, keratin, calnexin and LAMP1 was acquired from the human cDNA library at the Biomedical Research Core Facility at Westlake University.

All plasmid was constructed using homologus recombination. Briefly, PCR products of the vector and the insert gene with homologus sequences to the vectors were digested with DpnI (beyotime, D6257) followed by a clean-up step (Omega E.Z.N.A. Cycle Pure Kit). The cleaned vector and insert gene were then mixed together in 1:2 ratio for recombination using DNA exonase (Vazyme, C112) at 37°C for 30 minutes. The finished product was then used to transform Stbl3 chemically competent cells (Invitrogen, C737303) and allowed to recover for 1 h in SOB at 37°C. Once recovered, the cells were streak on LB plated supplied with corresponding antibiotics. Several single colonies were picked, amplified and sequenced for corrected.

### Site saturation mutagenesis

Site saturation mutagenesis of fluorescent proteins (FPs) was achieved through degenerated primers pairs according to Kille et. al. (Kille et al., 2013). The degenerate primers (a) NDT, (b) VHG, (c) TGG (N=any base, D=A, G, or T, V=A, G, or C, and H=A, C, or T) were used respectively for PCR amplification of fluorescence protein-encoding pET28a (+), generating PCR product of a, b, and c. PCR products were mixed in a ratio of a:b:c=12:9:1. Subsequently, PCR products were digested with DpnI and purified with a PCR clean-up kit. Followed by circularization through homologus recombination using CloneExpress kit (Vazyme, C112). The product was transformed into *E. coli* BL21 DE3* strain chemically competent cells (Invitrogen, C600003). The cells were streak on LB plates supplied with 50 µg/mL of kanamycin sulfate and 0.06 mM isopropyl β-D-1-thiogalactopyranoside (IPTG, Sangon Biotech, A100487).

### Library construction

Well isolated colonies on the plate from saturation mutagenesis that are fluorescently positive were picked and amplified in a deep 96-well plate containing 400 µL LB broth supplemented with 50 µg/mL kanamycin sulfate overnight at 37°C with gentle rotation (220 rpm). By the next day, the cell library is preserved as glycerol stock by inoculating 200 µL of overnight culture into a new deep 96-well plate containing 200 µL glycerol. The remaining culture were then amplified overnight and used for protein expression for screening. On the next day, the overnight culture is centrifuged at 4,000 xg for 10 minutes and stored in -80°C freezer for screening.

### General procedure for screening

The cell libraries were thawed and resuspended with lysis buffer (20 mM Tris, pH 7.5, 500 mM NaCl) in the presence of 1 mg/mL lysozyme (Sangon Biotech, A610308), 0.1 mg/mL DNase (Sigma-Roche, 10104159001) and 1mM phenylmethyl sulfonyl fluoride (Sangon Biotech, A610425). The plates were vortexed to completely resuspend pellets and allowed to shake for 1 h (37 °C, 250 rpm). The crude cell lysates were centrifuged at 4000 g for 30 min at 4 °C. Subsequently, the supernatants (200 µL) were transferred into another 96 well plates and the fluorescence lifetimes were recorded using a spectrofluorometer (FS5, Edinburgh Instruments).

### Protein expression and purification

Plasmids were transformed into E. coli BL21 DE3* strain. A single colony was picked and inoculated into 1 mL of LB media containing 50 µg/mL kanamycin sulfate and cultured at 37 °C with shaking at 220 rpm overnight. And then, the culture was transferred into large culturing flasks with 100 mL LB media containing 50 µg/mL kanamycin sulfate to allow further growth. When the OD600 reached 0.6-0.8, IPTG was introduced to a final concentration of 0.5 mM to induce the expression of FPs. Cultures were allowed to shake at 18°C for another 16-20 h, harvested and stored at -80°C freezer.

The FPs purification was carried out using the Ni^2+^ affinity chromatography. In brief, the *E. coli* cells were thawed in buffer (20 mM Tris, pH 7.5, 500 mM NaCl) and disrupted by sonication. The lysate was cleared out by centrifuging at 16,000 g for 60 min at 4 °C. The obtained supernatant was purified via Ni^2+^ affinity chromatography on chelating Sepharose (Bio-rad) and then washed three times with the same buffer containing 10 mM imidazole. Finally, FPs were eluted with the same buffer containing 300 mM imidazole.

### Extinction coefficient

Purified proteins were diluted such that the absorbance of maximum wavelength between 0.1 and 0.3 in the buffer (20 mM Tris, pH 7.5, 500 mM NaCl). Absorbance spectra were acquired using a spectrophotometer (Cary 3500, Agilent) in the wavelength range of 250-800 nm with a step size of 1 nm for all variants and corrected through subtracting the average absorbance at the range of 700-800 nm. 0.5 M NaOH was used to denaturing the variants of EBFP2, oxStayGold, mNeonGreen, mVenus, mOrange, mScalet and mKate2, but for the variants of mTurquoise2, 2 M NaOH was used to denaturing it. After that, Absorbance spectra of chromophore were acquired and corrected with the same methods. For the variants of EBFP2, based on the extinction coefficient of EBFP2 in the maximum absorbance wavelength (Ai et al., 2007), the extinction coefficient of chromophore of EBFP2 was calculated as 30249 M^−1^ cm^−1^ at 413 nm. For the variants of mTurquoise2, the extinction coefficient of chromophore was assumed as 46000 M^−1^ cm^−1^ at 460 nm (Sarkisyan et al., 2012). For the variants of oxStayGold, mNeonGreen, mVenus, mOrange, mScarlet and mKate2, the extinction coefficient of chromophore was assumed as 44000 M^−1^ cm^−1^ at 457 nm (Bindels et al., 2017). The concentration of the denatured chromophore was calculated through their respective extinction coefficient. Based on the concentration of chromophore, the extinction coefficient for variants was determined at absorbance of maximum peaked. The above procedure was repeated three times for all variants, and the average extinction coefficient were calculated.

### Quantum yield

Purified TRFP proteins were diluted such that the absorbance of maximum wavelength at around 0.02, 0.04, 0.06, 0.08 and 0.1 in the buffer (20 mM Tris, pH 7.5, 500 mM Nacl). Absorbance spectra were acquired using a spectrophotometer (Cary 3500, Agilent) in the wavelength range of 250-800 nm with a step size of 1 nm for all variants. After that, the fluorescence intensity of these samples was recorded using a plate reader (Spark, Tecan) with a step size of 1 nm. The excitation was set as 370 nm, 400 nm, 460 nm, 460nm, 480 nm, 510 nm, 540 nm and 550 nm for the variants of EBFP2, mTurquoise2, oxStayGold, mNeonGreen, mVenus, mOrange, mScarlet and mKate2, respectively. The range of emission was set as 390-600 nm, 420-700 nm, 480-700 nm, 480-700 nm, 505-700 nm, 540-700 nm, 555-800 nm and 580-800 nm for the variants of EBFP2, mTurquoise2, oxStayGold, mNeonGreen, mVenus, mOrange, mScarlet and mKate2, respectively. The bandwidth of excitation and emission was set as 10 nm.

Absorbance spectra were corrected through subtracting the average absorbance at the range of 700-800 nm. The spectral area was obtained through integrating the range of emission for each variant. After that, the absorbance at excitation was plotted versus the spectral area and the slope (α) of the line was determined using linear regression. The following equation was used to calculate the quantum yield (QY):

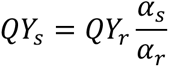

In the equation, s and r indicates the sample and reference, respectively. Result from the solvent for all measurement is water, so, the effect of unequal refractive indices is omitted in this equation. EBFP2 (Ai et al., 2007), mTurquoise2 (Goedhart et al., 2012), oxStayGold (Hirano et al., 2022), mNeonGreen (Shaner et al., 2013), mVenus (Kremers et al., 2006), mOrange (Shaner et al., 2004), mScarlet (Bindels et al., 2017) and mKate2 (Shcherbo et al., 2009) were used as the reference for their respective variants.

### *In vitro* fluorescence lifetime measurement

Fluorescence lifetime measurements were performed on a spectrofluorometer (FS5, Edinburgh Instruments). The amount of purified TRFPs used was adjusted to a level where the absorbance at the maximum wavelength was around 0.1 in the buffer (20 mM Tris, pH 7.5, 500 mM Nacl). For determining the fluorescence lifetime, samples were excited using 10 MHz pulse laser to allow a complete decay and 1×10^4^ photons were collected for each sample. A 405 nm pulsed laser was used for the variants of EBFP2 and mTurquoise2; 488 nm for the variants of oxStayGold, mNeonGreen and mVenus; and 510 nm for the variants of mOrange, mScarlet and mKate. Fluorescence lifetimes were calculated by fitting the decay curve with mono, bi, or triexponential decay models (χ^2^ < 1.2). The reported fluorescence lifetime of the TRFPs represents the average lifetime of three experimental replicates.

### Ultrafast transient absorption (TA) spectroscopy

For the assay of ultrafast transient absorption spectroscopy (Coquelle et al., 2018), the variants (T203G and T62C) of mTurquoise2 were diluted to 150 μM in the buffer (50 mM HEPES, 50 mM NaCl, pH 8.0). A pump beam (spot diameter 250 μm, pulse length150 fs) with 370 nm was used excited the two variants and it was obtained through doubling 800 nm fundamental pulses. The white light with continuum probe beam was generated by focusing the fundamental beam in a 3 mm CaF2 crystal. For measurement, samples were placed in a cuvette with a path length of 1 mm. To avoid multiphoton process, the data of TA spectrum were collected at a low excitation-energy density (0.228 mJ/cm^2^). All experimental data were corrected by the system from Time-Tech Spectra, Inc. Decay associated spectra were obtained by global fitting with two or three exponential functions and a constant convoluted with a Gaussian pulse of 138 fs.

### Structure simulation

All structure simulation was performed with Rosetta (Leaver-Fay et al., 2011). Firstly, parameter files of chromophores were generated allowing Rosetta to identify them as polymers within the structure. Next, MutateResidue Mover was used to introduce single-point mutations. Following this, side chain packing on the mutated residues was performed to avoid clashes. In cases where clashes persisted, FastRelax Mover was employed to relax either the mutated residue or its surrounding residues, thereby obtaining the final structural model. The structure of EBFP2 and mKate2 was simulated through Rosetta based on the template of BFP (PDB ID: 1bfp) and mKate (PDB ID: 3bxa). The structures of mTurquoise2^K206A^ (PDB ID: 3ztf), and StayGold (PDB ID: 8bxt) were used as the template for the modeling of the variants of mTurquoise2 and oxStayGold, respectively. The structures of mNeonGreen (PDB ID: 5ltp), mVenus (PDB ID: 7pnn), mOrange (PDB ID: 2h5o) and mScarlet (PDB ID: 5lk4) were used as the template for their respective variants.

### MD simulation

For each system, the initial structure with the chromophore in its neutral state was immersed in the centre of a truncated octahedral box of water molecules, and all of the protein atoms were no less than 12 Å from the boundary of the water box. The AmberFF14SB, gaff2 and TIP3P force field parameters were used to describe the protein, chromophore (ground state) and water molecules, respectively. To describe the chromophore in excited state, we employed the RESP method to recalculate its charges, and adjusted the related bond lengths, bond angles, and dihedral angle parameters based on frequency analysis. To remove bad contacts before the simulation, 20,000 steps of steepest descent followed by 20,000 steps of conjugate gradient energy minimizations were conducted. The relaxed structure was heated to 300 K in 100 ps, during which all atoms in the protein were restrained with a force constant of 10 kcal/mol·Å2. All bonds with hydrogen atoms were fixed using the SHAKE algorithm. The particle mesh Ewald method with a 10 Å cutoff in real space was used to calculate electrostatic interaction. A Langevin thermostat with a collision frequency of 1.0 ps−1 was used to regulate temperature. Isotropic pressure coupling with a relaxation time of 2 ps was used to maintain the pressure to 1 atm. The integration time step was set to 2 fs. Trajectories were saved every 1 ps, and MD simulations were extended to 300 ns for each system. Simulations for the three systems were conducted using AMBER22 package.

### QM and QM/MM simulation

Using the dihedral angle *tau* as the reaction coordinate, we scanned the potential energy surfaces of the chromophore in both the ground and excited states. The quantum calculations were carried out at the B3LYP/def2-TZVP and TD-cam-B3LYP/def2-TZVP level, respectively. Each time the value of *tau* was changed, it was fixed, and then the other degrees of freedom within the molecule were optimized.

Starting from typical structures in the molecular dynamics simulation trajectories, we also performed QM/MM simulations. The QM region included the chromophore molecule, while the remaining protein was described using the Amber FF14SB force field. The timestep for the ground state simulation was 1 fs, and for the excited state simulation, it was 0.5 fs. To ensure the statistical significance of the simulation results, each simulation was repeated 10 times.

### Cell culture and transfection

HEK293T (ATCC, CRL-3216), HeLa (ATCC, CCL-2), and U-2 OS (ATCC, HTB-96) were purchased from ATCC. The cells were cultured in Dulbecco’s Modified Eagle’s Medium (DMEM, Gibco, 11995-065) supplemented with 10% Fetal Bovine Serum (FBS, Cellmax, SA211.02, LOT#20220322) and 1x penicillin-streptomycin-glutamine (PSQ, Gibco, 10378-016). The cells were incubated at 37°C in a 5% CO_2_ atmosphere using a Heracell Vios 160i CO2 incubator (Thermo Fisher Scientific). Cells were seeded at 30% confluency in 35 mm glass-bottom culture dishes (Cellvis, D35-20-1.5H) 24 hours prior to the transfection. The dishes culturing HEK293T cells were pre-treated with poly-L-lysine (Sigma, P4707) to enhance cell adherence.

For transfection, cells at 40% confluency were transfected with 1 µg of mammalian expression plasmid DNA containing gene of interest using 3 µL X-tremeGENE 9 DNA transfection reagent (Roche, 6365809001) diluted in 100 µL 1x OPTI-MEM I reduced serum medium (Gibco, 31985-062). Subsequently, the cells were then incubated at 37°C in a 5% CO_2_ environment to facilitate protein expression. After 24 hours, the medium was replaced with phenol-free DMEM/F12 (1:1) medium (Gibco, 21041-025) supplemented with 10% FBS and 1x PSQ for imaging. For co-transfection experiments, the ratio of plasmids DNA to X-tremeGENE 9 DNA transfection reagent is adjusted to 1:1.5 (for 9 TRFPs) or 1:2 (for 6 TRFPs), with the subsequent procedures remaining the same as previously described.

### Stable cell lines establishment

Cells expressing the tr-Fucci sensor were engineered using the PiggyBac (PB) transposon system(Randolph et al., 2017). Briefly, the tr-Fucci sensor was subcloned into XLone vector, an improved Tet-On system that contains transposon-specific inverted terminal repeat sequences (ITRs) flanking the gene of interest. XLone-tr-Fucci construct and PB transposase (a gift from Prof. Xiaojun Lian, PSU) were co-transfected into HeLa cells. Selection of positive cells was carried out using blasticidin (10 µg/mL, Sigma, SBR00022). To induce tr-Fucci expression for visualization, cells were treated with 2 µg/mL doxycycline (TCI, D4116) 24 hours prior to the experiments.

CDK2-mVenus and CDK4/6-mScarlet were transduced into tr-Fucci HeLa using a third-generation lentivirus vector(Dull et al., 1998). In summary, pCDH-CDK2-mVenus-P2A-CDK4/6-mScarlet was co-transfected with the necessary packaging plasmids (pMDLg and pRSV-Rev) and the envelope-expressing plasmid (pCMV-VSV-G) into HEK293T cells at 50% confluency. 12 hours post-transfection, the media was replaced with 1 mL of DMEM containing 20% FBS and 1 mM sodium pyruvate (Sangon Biotech A600884). After 24 additional hours, 1 mL lentivirus-containing media were collected and combined with 1 mL DMEM containing 20% FBS and 20 µM polybrene (beyotime, C0351). This mixture was used to infect tr-Fucci HeLa cells. The cells were trypsinized and sorted by flow cytometry 36 hours post-infection to select for mVenus and mScarlet positive cells.

### Fluorescence lifetime imaging microscopy (FLIM)

FLIM experiments were conducted using a Leica STELLARIS 8 FALCON confocal microscope, equipped with an adjustable pulsed white light laser (WLL) and hybrid detectors (HyD), utilizing a 63x/1.4 oil immersion objective lens (Leica, Microsystems). The HyD integrated the features of a photomultiplier tube (PMT) and avalanche photodiodes (APDs), enabling the single-photon counting; this, combined with the pulsed laser, allows for FLIM to be performed in a streamlined manner without the need for additional modules. An 80 MHz WLL is used for most FLIM applications unless specified otherwise. The live-cell FLIM imaging environment was maintained at 37°C, under 5% CO_2_, using a stage top incubator and an incubation system that encloses the microscope (TokiHit, DMi8TB). To excite the TRFPs, the WLL was tuned to 440 nm for mTurquoise2-based, 488 nm for mNeonGreen-based, and 560 nm for mScarlet-based TRFPs. Emissions were detected by HyD X (for mTurquoise2 and mNeonGreen) or HyD R (for mScarlet) detectors within the wavelength ranges of 450-600 nm, 500-650 nm, and 570-700 nm, respectively. All TRFP FLIM multiplex imaging were performed in line sequential mode, with each line scanned 6 times for line accumulation at a 200 Hz scan speed. Images were captured in a 1024 x 1024 format with pixel sizes ranging from 80 to 100 nm. Z-stack TRFP FLIM multiplexing were performed in the same manner as previously described with the additional of 2 µm z-scan at 200 nm step size.

### Phasor plot Analysis and TRFPs separation

The phasor plot is a method used to transform fluorescence lifetime decay data recorded at each pixel of a micrograph. This data is represented as a scatter of dots within a 2D semicircle, where the position of each dot corresponds to the average lifetime of that pixel. The dots on the phasor plot are color-coded in a rainbow spectrum, with blue indicating low and red indicating a higher density (hotspot) of similar lifetimes. When a micrograph captures a single lifetime species, the corresponding hotspot on the phasor plot will be located on the perimeter of the semicircle. However, if multiple lifetime species are present, the hotspot(s) will appear inside the semicircle. For instance, a micrograph with two lifetime species, the phasor plot can yield three possible outcomes: (1) two distinct hotspots, indicating that the two species have distinguishable lifetimes and are spatially segregated with minimal overlap, (2) hotspots that appear smeared, indicating distinguishable lifetimes but significant spatial overlaps, and (3) a single hotspot, suggesting that the lifetimes of the two species are nearly indistinguishable, regardless of their spatial separation. Noted that in the second outcome, the smear is caused by the pixels containing different ratios of the two lifetime species. To separate lifetime species with significant spatial overlap, a circular analytical tool (cursor) is used to identify the pure species at each end of the smear. The proportions of mixed species along the smear can then be quantified and reassigned to their respective locations on the micrograph, delineating the contribution of each lifetime species(Digman et al., 2008; Wang et al., 2021).

Phasor analysis became more complex for micrograph containing three or more lifetime species. The ideal scenario is to observe three or multiple distinct hotspots or at least two or more smeared clusters. As long as the lifetime differences between each species are substantial, separation can be achieved even when the labeled species significantly overlaps spatially. For this reason, the lifetime of each TRFP is measured and critically considered before conducting FLIM multiplexing experiments.

Following this principle, TRPFs were separated in FLIM multiplexing images through phasor plot analysis using the LAS X FLIM/FCS software (Leica Microsystem). This was achieved by positioning cursors on the phasor plot clusters that delineate the pure species-labeled structures in the micrograph. STED-FLIM phasor plot analysis is similar to that of the confocal-FLIM, except the lifetime of each oxStayGold-based TRFPs will be shorter after the STED beam exposure.

### Time-resolved multiplexing using a conventional confocal microscope

U-2 OS cells transfected with subcellular-targeted mNeronGreen-based TRFPs or mScarlet-based TRFPs were imagined using using TauSeparation mode on a Leica Stellaris 5 confocal microscope equipped with a single frequency (80 MHz) pulsed white light laser (WLL) and standard hybrid detectors S (HyD S), utilizing a 63x/1.4 oil immersion objective lens (Leica Microsystems). TauSeparation mode is a standard built-in acquisition mode that allows the separation of signals of the same emission wavelength based on lifetime during live preview. Different lifetime channels were set during preview and signals of different lifetime will be saved as different channels.

### *In cell* fluorescence lifetime measurement

To determine the fluorescence lifetime of mTurquoise2-, mNeonGreen-, and mScarlet-based TRFPs inside the cell. HEK293T cells were transiently transfected with a single TRFP and imaged on a Leica STELLARIS 8 FALCON confocal microscope. The imaging procedures are the same as those described in the FLIM section, except a 10 MHz WLL was used to obtain the complete decay curve, and 1,000 photons per pixel were collected. The fluorescence lifetime was calculated using LAS X FLIM/FCS software (Leica Microsystems) by fitting the decay curve (n-exponential deconvolution method) with mono-, bi-, or tri-exponential decay models (χ2 < 1.2).

The fluorescence lifetime of subcellular-targeting mTurquoise2-, mNeonGreen-, and mScarlet-based TRFPs was measured in HeLa cells. HeLa cells were transiently transfected with a single TRFP and imaged in the same manner as the HEK293T cells, except 500 photons per pixel were collected. The fluorescence lifetime of each subcellular-targeting TRFP was calculated by fitting the decay curve with mono-, bi-, or tri-exponential decay models (χ2 < 1.2). The reported fluorescence lifetime of the TRFPs represents the average lifetime of three experimental replicates.

### Time-lapse FLIM imaging

Time-lapse FLIM imaging was performed using HeLa stable cells expressing tr-Fucci, CDK2-mVenus, and CDK4/6-mScarlet, seeded on 35 mm glass-bottom dishes. The imaging procedure was the same as described in the FLIM multiplexing section, except the images were captured in a 512×512 format with a pixel size at 481 nm and were taken for 48 hours at 15-minute intervals. Since the tr-Fucci sensor, CDK2-mVenus and CDK4/6-mScarlet are expressed at different stages of the cell cycle. To monitor the expression of and to separate each TRFP-labeled protein, cursors were placed on the cluster location representing each individual TRFP during phasor plot analysis.

### STED-FLIM microscopy

STED-FLIM images were acquired on a Leica STELLARIS 8 FALCON confocal microscope (Leica Microsystems), equipped with a pulsed white light laser (WLL), an HC PL APO 100x/1.4 oil STED WHITE lens, and STED laser beams (592 and 775 nm). The images were captured with gating of 1 - 6 ns, pixel size ranging from 15 to 25 nm, and dwell times of 1.5 - 3 µs. Each line was scanned 3 – 5 times (line accumulation) to achieve optimal photon counts (approximately 120 photons per pixel). Noted that the photon counts are lower than in confocal-FLIM due to STED beam depletion, but the collected counts are sufficed for phasor plot analysis in species separation. oxStayGold-based TRFPs were excited at 488 nm and depleted with a 592 nm STED beam.

For multi-color STED FLIM multiplexing, images were recorded in frame sequential mode; HaloTags (9 and 11) labeled SiR was excited at 650 nm and depleted with a pulsed 775 nm STED beam. Images were captured with gating of 1 - 6 ns, pixel size ranging from 15 to 25 nm, and dwell times of 1.5 - 3 µs. Each line was scanned 3 – 5 times (line accumulation) to achieve optimal photon counts. Sequentially the oxStayGold-based TRFPs were imaged in the same manner as described previously.

### Software and image processing

Graphs were generated using Microsoft PowerPoint and Excel 2021 or OriginLab. All images were processed with Fiji ImageJ (Rueden et al., 2017; Schindelin et al., 2012) and LAX S FLIM/FCS software (Leica Microsystems). Movies were assembled using Python with the packages of numpy, pandas, matplotlib, pillow and opencv.

---

**Movie 1, Tracking of cell cycle in HeLa cell via the tr-Fucci.** FLIM images were collected every 15 mins. Fluorescence lifetime was calculated with phasor plot. Cell-cycle phases were assigned based on the principles in figure 5A. Movies were assembled using Python with the packages of numpy, pandas, matplotlib, pillow and opencv.

## REFERENCE

Ai, H.W., Shaner, N.C., Cheng, Z.H., Tsien, R.Y., and Campbell, R.E. (2007). Exploration of new chromophore structures leads to the identification of improved blue fluorescent proteins. Biochemistry 46, 5904–5910.

Ando, R., Shimozono, S., Ago, H., Takagi, M., Sugiyama, M., Kurokawa, H., Hirano, M., Niino, Y., Ueno, G., Ishidate, F., et al. (2023). StayGold variants for molecular fusion and membrane-targeting applications. Nat. Methods 10.1038/s41592-023-02085-6.

Bajar, B.T., Lam, A.J., Badiee, R.K., Oh, Y.H., Chu, J., Zhou, X.X., Kim, N., Kim, B.B., Chung, M.Y., Yablonovitch, A.L., et al. (2016). Fluorescent indicators for simultaneous reporting of all four cell cycle phases. Nat. Methods 13, 993–996.

Betzig, E., Patterson, G.H., Sougrat, R., Lindwasser, O.W., Olenych, S., Bonifacino, J.S., Davidson, M.W., Lippincott-Schwartz, J., and Hess, H.F. (2006). Imaging intracellular fluorescent proteins at nanometer resolution. Science 313, 1642–1645.

Bindels, D.S., Haarbosch, L., van Weeren, L., Postma, M., Wieser, K.E., Mastop, M., Aumonier, S., Gotthard, G., Royant, A., Hink, M.A., et al. (2017). mScarlet: a bright monomeric red fluorescent protein for cellular imaging. Nat. Methods 14, 53–56.

Chudakov, D.M., Belousov, V.V., Zaraisky, A.G., Novoselov, V.V., Staroverov, D.B., Zorov, D.B., Lukyanov, S., and Lukyanov, K.A. (2003). Kindling fluorescent proteins for precise photolabeling. Nat. Biotechnol. 21, 452–452.

Chudakov, D.M., Verkhusha, V.V., Staroverov, D.B., Souslova, E.A., Lukyanov, S., and Lukyanov, K.A. (2004). Photoswitchable cyan fluorescent protein for protein tracking. Nat. Biotechnol. 22, 1435–1439.

Coudreuse, D., and Nurse, P. (2010). Driving the cell cycle with a minimal CDK control network. Nature 468, 1074–1079.

Covert, M.W., Regot, S., Hughey, J., Bajar, B., and Carrasco, S. (2015). High-sensitivity measurements of multiple kinase activities in live single cells. Mol. Biol. Cell 26, 1724–1734.

Datta, R., Heaster, T.M., Sharick, J.T., Gillette, A.A., and Skala, M.C. (2020). Fluorescence lifetime imaging microscopy: fundamentals and advances in instrumentation, analysis, and applications. J. Biomed. Opt. 25, 1–43.

Doksani, Y., Wu, J.Y., de Lange, T., and Zhuang, X.W. (2013). Super-Resolution Fluorescence Imaging of Telomeres Reveals TRF2-Dependent T-loop Formation. Cell 155, 345–356.

Filonov, G.S., Piatkevich, K.D., Ting, L.M., Zhang, J.H., Kim, K., and Verkhusha, V.V. (2011). Bright and stable near-infrared fluorescent protein for imaging. Nat. Biotechnol. 29, 757–761.

Frei, M.S., Tarnawski, M., Roberti, M.J., Koch, B., Hiblot, J., and Johnsson, K. (2022). Engineered HaloTag variants for fluorescence lifetime multiplexing. Nat. Methods 19, 65–70.

Goedhart, J., van Weeren, L., Hink, M.A., Vischer, N.O.E., Jalink, K., and Gadella, T.W.J. (2010). Bright cyan fluorescent protein variants identified by fluorescence lifetime screening. Nat. Methods 7, 137–139.

Goedhart, J., von Stetten, D., Noirclerc-Savoye, M., Lelimousin, M., Joosen, L., Hink, M.A., van Weeren, L., Gadella, T.W.J., and Royant, A. (2012). Structure-guided evolution of cyan fluorescent proteins towards a quantum yield of 93%. Nat. Commun. 3, 751.

Greenwald, E.C., Mehta, S., and Zhang, J. (2018). Genetically Encoded Fluorescent Biosensors Illuminate the Spatiotemporal Regulation of Signaling Networks. Chem. Rev. 118, 11707–11794.

Grotjohann, T., Testa, I., Leutenegger, M., Bock, H., Urban, N.T., Lavoie-Cardinal, F., Willig, K.I., Eggeling, C., Jakobs, S., and Hell, S.W. (2011). Diffraction-unlimited all-optical imaging and writing with a photochromic GFP. Nature 478, 204–208.

Handlin, L.J., and Dai, G. (2023). Direct regulation of the voltage sensor of HCN channels by membrane lipid compartmentalization. Nat. Commun. 14, 6595.

Heim, R., Prasher, D.C., and Tsien, R.Y. (1994). Wavelength Mutations and Posttranslational Autoxidation of Green Fluorescent Protein. Proc. Natl. Acad. Sci. U.S.A. 91, 12501–12504.

Hell, S.W. (2003). Toward fluorescence nanoscopy. Nat. Biotechnol. 21, 1347–1355.

Hirano, M., Ando, R., Shimozono, S., Sugiyama, M., Takeda, N., Kurokawa, H., Deguchi, R., Endo, K., Haga, K., Takai-Todaka, R., et al. (2022). A highly photostable and bright green fluorescent protein. Nat. Biotechnol. 40, 1132–1142.

Kremers, G.J., Goedhart, J., van Munster, E.B., and Gadella, T.W.J. (2006). Cyan and yellow super fluorescent proteins with improved brightness, protein folding, and FRET Forster radius. Biochemistry 45, 6570–6580.

Lee, H.N., Mehta, S., and Zhang, J. (2020). Recent advances in the use of genetically encodable optical tools to elicit and monitor signaling events. Curr. Opin. Cell Biol. 63, 114–124.

Lee, S.J.R., Escobedo-Lozoya, Y., Szatmari, E.M., and Yasuda, R. (2009). Activation of CaMKII in single dendritic spines during long-term potentiation. Nature 458, 299–304.

Linghu, C.Y., Johnson, S.L., Valdes, P.A., Shemesh, O.A., Park, W.M., Park, D., Piatkevich, K.D., Wassie, A.T., Liu, Y.X., An, B., et al. (2020). Spatial Multiplexing of Fluorescent Reporters for Imaging Signaling Network Dynamics. Cell 183, 1682–1698.

Lu, T., Ang, C.E., and Zhuang, X.W. (2023). Spatially resolved epigenomic profiling of single cells in complex tissues. Cell 186, 2275–2279.

Matz, M.V., Fradkov, A.F., Labas, Y.A., Savitsky, A.P., Zaraisky, A.G., Markelov, M.L., and Lukyanov, S.A. (1999). Fluorescent proteins from nonbioluminescent Anthozoa species. Nat. Biotechnol. 17, 969–973.

Moerner, W.E., and Orrit, M. (1999). Illuminating single molecules in condensed matter. Science 283, 1670–1676.

Niehörster, T., Löschberger, A., Gregor, I., Krämer, B., Rahn, H.J., Patting, M., Koberling, F., Enderlein, J., and Sauer, M. (2016). Multi-target spectrally resolved fluorescence lifetime imaging microscopy. Nat. Methods 13, 257–262.

Ogikubo, S., Nakabayashi, T., Adachi, T., Islam, M.S., Yoshizawa, T., Kinjo, M., and Ohta, N. (2011). Intracellular pH Sensing Using Autofluorescence Lifetime Microscopy. J. Phys. Chem. B 115, 10385–10390.

Okabe, K., Inada, N., Gota, C., Harada, Y., Funatsu, T., and Uchiyama, S. (2012). Intracellular temperature mapping with a fluorescent polymeric thermometer and fluorescence lifetime imaging microscopy. Nat. Commun. 3, 705.

Pédelacq, J.D., Cabantous, S., Tran, T., Terwilliger, T.C., and Waldo, G.S. (2006). Engineering and characterization of a superfolder green fluorescent protein (vol 24, pg 79, 2005). Nat. Biotechnol. 24, 1170–1170.

Pisfil, M.G., Nadelson, I., Bergner, B., Rottmeier, S., Thomae, A.W., and Dietzel, S. (2022). Stimulated emission depletion microscopy with a single depletion laser using five fluorochromes and fluorescence lifetime phasor separation. Sci. Rep. 12, 14027.

Qian, Y., Celiker, O.T., Wang, Z., Guner-Ataman, B., and Boyden, E.S. (2023). Temporally multiplexed imaging of dynamic signaling networks in living cells. Cell 186, 5656–5672.

Qian, Y., Piatkevich, K.D., Mc Larney, B., Abdelfattah, A.S., Mehta, S., Murdock, M.H., Gottschalk, S., Molina, R.S., Zhang, W., Chen, Y.C., et al. (2019). A genetically encoded near-infrared fluorescent calcium ion indicator. Nat. Methods 16, 171–174.

Rizzo, M.A., Springer, G.H., Granada, B., and Piston, D.W. (2004). An improved cyan fluorescent protein variant useful for FRET. Nat. Biotechnol. 22, 445–449.

Sakaue-Sawano, A., Kurokawa, H., Morimura, T., Hanyu, A., Hama, H., Osawa, H., Kashiwagi, S., Fukami, K., Miyata, T., Miyoshi, H., et al. (2008). Visualizing spatiotemporal dynamics of multicellular cell-cycle progression. Cell 132, 487–498.

Sakaue-Sawano, A., Yo, M., Komatsu, N., Hiratsuka, T., Kogure, T., Hoshida, T., Goshima, N., Matsuda, M., Miyoshi, H., and Miyawaki, A. (2017). Genetically Encoded Tools for Optical Dissection of the Mammalian Cell Cycle. Mol. Cell. 68, 626–640.

Scipioni, L., Rossetta, A., Tedeschi, G., and Gratton, E. (2021). Phasor S-FLIM: a new paradigm for fast and robust spectral fluorescence lifetime imaging. Nat. Methods 18, 542–550.

Shaner, N.C., Campbell, R.E., Steinbach, P.A., Giepmans, B.N.G., Palmer, A.E., and Tsien, R.Y. (2004). Improved monomeric red, orange and yellow fluorescent proteins derived from Discosoma sp. red fluorescent protein. Nat. Biotechnol. 22, 1567–1572.

Shaner, N.C., Lambert, G.G., Chammas, A., Ni, Y.H., Cranfill, P.J., Baird, M.A., Sell, B.R., Allen, J.R., Day, R.N., Israelsson, M., et al. (2013). A bright monomeric green fluorescent protein derived from Branchiostoma lanceolatum. Nat. Methods 10, 407–409.

Shcherbo, D., Murphy, C.S., Ermakova, G.V., Solovieva, E.A., Chepurnykh, T.V., Shcheglov, A.S., Verkhusha, V.V., Pletnev, V.Z., Hazelwood, K.L., Roche, P.M., et al. (2009). Far-red fluorescent tags for protein imaging in living tissues. Biochem. J. 418, 567–574.

Spencer, S.L., Cappell, S.D., Tsai, F.C., Overton, K.W., Wang, C.L., and Meyer, T. (2013). The Proliferation-Quiescence Decision Is Controlled by a Bifurcation in CDK2 Activity at Mitotic Exit. Cell 155, 369–383.

Starling, T., Carlon-Andres, I., Iliopoulou, M., Kraemer, B., Loidolt-Krueger, M., Williamson, D.J., and Padilla-Parra, S. (2023). Multicolor lifetime imaging and its application to HIV-1 uptake. Nat. Commun. 14, 4994.

Stringari, C., Wang, H., Geyfman, M., Crosignani, V., Kumar, V., Takahashi, J.S., Andersen, B., and Gratton, E. (2015). In Vivo Single-Cell Detection of Metabolic Oscillations in Stem Cells. Cell Rep. 10, 1–7.

Sun, Y.H., Hatami, N., Yee, M., Phipps, J., Elson, D.S., Gorin, F., Schrot, R.J., and Marcu, L. (2010). Fluorescence lifetime imaging microscopy for brain tumor image-guided surgery. J. Biomed. Opt. 15, 056022.

Tian, X.D., Zhang, Y.Y., Li, X.Y., Xiong, Y., Wu, T.C., and Ai, H.W. (2022). A luciferase prosubstrate and a red bioluminescent calcium indicator for imaging neuronal activity in mice. Nat. Commun. 13, 3967.

Valm, A.M., Cohen, S., Legant, W.R., Melunis, J., Hershberg, U., Wait, E., Cohen, A.R., Davidson, M.W., Betzig, E., and Lippincott-Schwartz, J. (2017). Applying systems-level spectral imaging and analysis to reveal the organelle interactome. Nature 546, 162–167.

Wang, L., Wu, C.L., Peng, W.L., Zhou, Z.L., Zeng, J.Z., Li, X.L., Yang, Y.N., Yu, S.G., Zou, Y., Huang, M., et al. (2022). A high-performance genetically encoded fluorescent indicator for in vivo cAMP imaging. Nat. Commun. 13, 5363.

Wolff, J.O., Scheiderer, L., Engelhardt, T., Engelhardt, J., Matthias, J., and Hell, S.W. (2023). MINFLUX dissects the unimpeded walking of kinesin-1. Science 379, 1004–1010.

Wu, T.C., Kumar, M., Zhang, J., Zhao, S.Y., Drobizhev, M., McCollum, M., Anderson, C.T., Wang, Y., Pokorny, A., Tian, X.D., et al. (2023). A genetically encoded far-red fluorescent indicator for imaging synaptically released Zn2+. Sci. Adv. 9, eadd2058.

Yang, H.W., Cappell, S.D., Jaimovich, A., Liu, C., Chung, M.Y., Daigh, L.H., Pack, L.R., Fan, Y., Regot, S., Covert, M., et al. (2020). Stress-mediated exit to quiescence restricted by increasing persistence in CDK4/6 activation. Elife 9, e44571.

Yaseen, M.A., Sutin, J., Wu, W.C., Fu, B.Y., Uhlirova, H., Devor, A., Boas, D.A., and Sakadzic, S. (2017). Fluorescence lifetime microscopy of NADH distinguishes alterations in cerebral metabolism. Biomed. Opt. Express 8, 2368–2385.

Yu, D., Baird, M.A., Allen, J.R., Howe, E.S., Klassen, M.P., Reade, A., Makhijani, K., Song, Y.Q., Liu, S.M., Murthy, Z., et al. (2015). A naturally monomeric infrared fluorescent protein for protein labeling. Nat. Methods 12, 763–765.

Yu, D., Gustafson, W.C., Han, C., Lafaye, C., Noirclerc-Savoye, M., Ge, W.P., Thayer, D.A., Huang, H., Kornberg, T.B., Royant, A., et al. (2014). An improved monomeric infrared fluorescent protein for neuronal and tumour brain imaging. Nat. Commun. 5, 3626.

Zhao, Y.X., Araki, S., Jiahui, W.H., Teramoto, T., Chang, Y.F., Nakano, M., Abdelfattah, A.S., Fujiwara, M., Ishihara, T., Nagai, T., et al. (2011). An Expanded Palette of Genetically Encoded Ca Indicators. Science 333, 1888–1891.

Zhu, W.C., Takeuchi, S., Imai, S., Terada, T., Ueda, T., Nasu, Y., Terai, T., and Campbell, R.E. (2023). Chemigenetic indicators based on synthetic chelators and green fluorescent protein. Nat. Chem. Biol. 19, 38–44.

## METHOD REFERENCES

Coquelle, N., Sliwa, M., Woodhouse, J., Schirò, G., Adam, V., Aquila, A., Barends, T.R.M., Boutet, S., Byrdin, M., Carbajo, S., et al. (2018). Chromophore twisting in the excited state of a photoswitchable fluorescent protein captured by time-resolved serial femtosecond crystallography. Nat. Chem. 10, 31–37.

Digman, M.A., Caiolfa, V.R., Zamai, M., and Gratton, E. (2008). The phasor approach to fluorescence lifetime imaging analysis. Biophys. J. 94, L14–L16.

Dull, T., Zufferey, R., Kelly, M., Mandel, R.J., Nguyen, M., Trono, D., and Naldini, L. (1998). A third-generation lentivirus vector with a conditional packaging system. J. Virol. 72, 8463–8471.

Kille, S., Acevedo-Rocha, C.G., Parra, L.P., Zhang, Z.G., Opperman, D.J., Reetz, M.T., and Acevedo, J.P. (2013). Reducing Codon Redundancy and Screening Effort of Combinatorial Protein Libraries Created by Saturation Mutagenesis. ACS Synth. Biol. 2, 83–92.

Leaver-Fay, A., Tyka, M., Lewis, S.M., Lange, O.F., Thompson, J., Jacak, R., Kaufman, K., Renfrew, P.D., Smith, C.A., Sheffler, W., et al. (2011). Rosetta3: An Object-Oriented Software Suite for the Simulation and Design of Macromolecules. Meth. Enzymol. 487, 545–574.

Randolph, L.N., Bao, X.P., Zhou, C.K., and Lian, X.J. (2017). An all-in-one, Tet-On 3G inducible PiggyBac system for human pluripotent stem cells and derivatives. Sci. Rep. 7, 1549.

Rueden, C.T., Schindelin, J., Hiner, M.C., DeZonia, B.E., Walter, A.E., Arena, E.T., and Eliceiri, K.W. (2017). ImageJ2: ImageJ for the next generation of scientific image data. BMC Bioinform. 18, 529.

Sarkisyan, K.S., Yampolsky, I.V., Solntsev, K.M., Lukyanov, S.A., Lukyanov, K.A., and Mishin, A.S. (2012). Tryptophan-based chromophore in fluorescent proteins can be anionic. Sci. Rep. 2, 608.

Schindelin, J., Arganda-Carreras, I., Frise, E., Kaynig, V., Longair, M., Pietzsch, T., Preibisch, S., Rueden, C., Saalfeld, S., Schmid, B., et al. (2012). Fiji: an open-source platform for biological-image analysis. Nat. Methods 9, 676–682.

Wang, P., Hecht, F., Ossato, G., Tille, S., Fraser, S.E., and Junge, J.A. (2021). Complex wavelet filter improves FLIM phasors for photon starved imaging experiments. Biomed. Opt. Express 12, 3463–3473.

