## Supplemental Information for "Time-resolved fluorescent proteins towards fluorescence microscopy in the temporal and spectral domains"

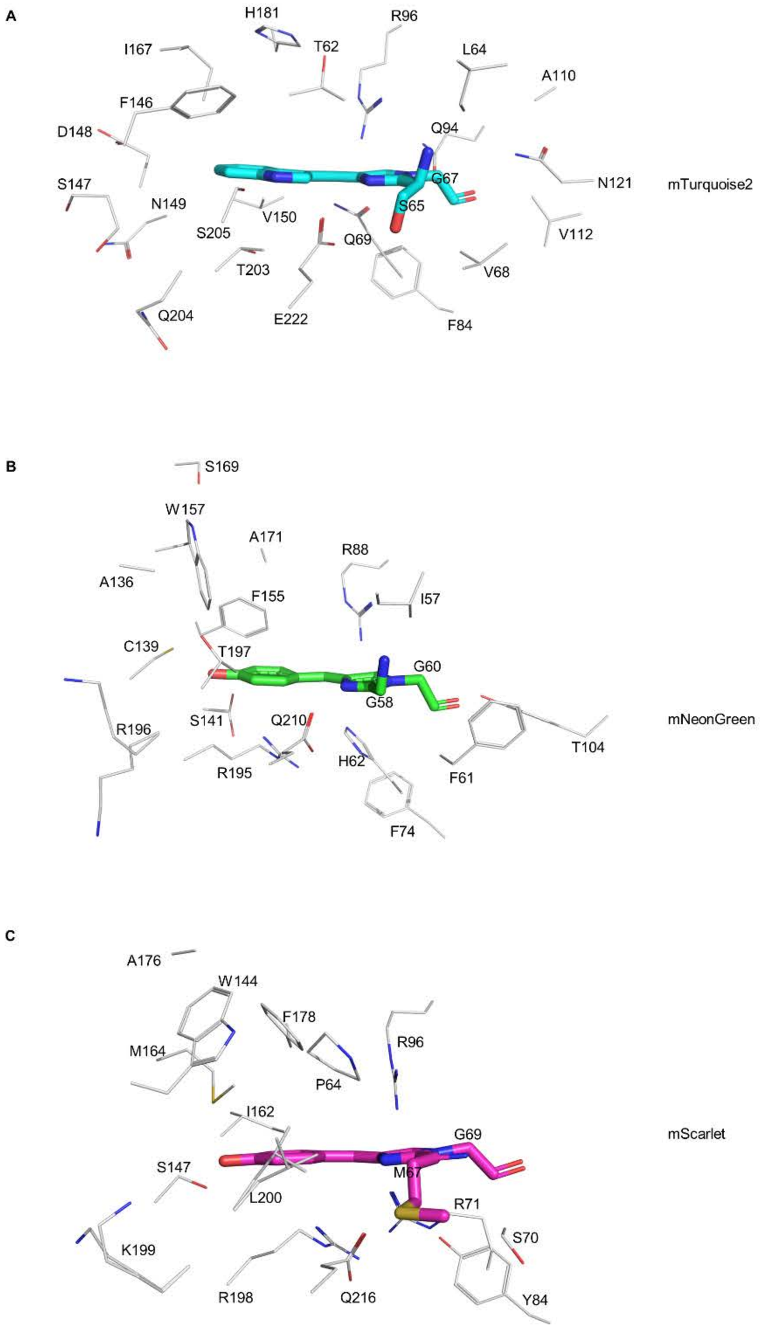


**Figure S1. Selection of residues for saturation mutagenesis.** Residues around from chromophores of mTurquoise2 (A), mNeonGreen (B) and mScarlet (C). Residues with side chains less than 6 Å away from chromophores were selected. For mTurquoise2, the structure of mTurquoise2$^{K206A}$ (PDB ID: 3ztf) was used to select the residues. PDB ID for mNeonGreen and mScarlet are 5ltr and 5lk4, respectively.

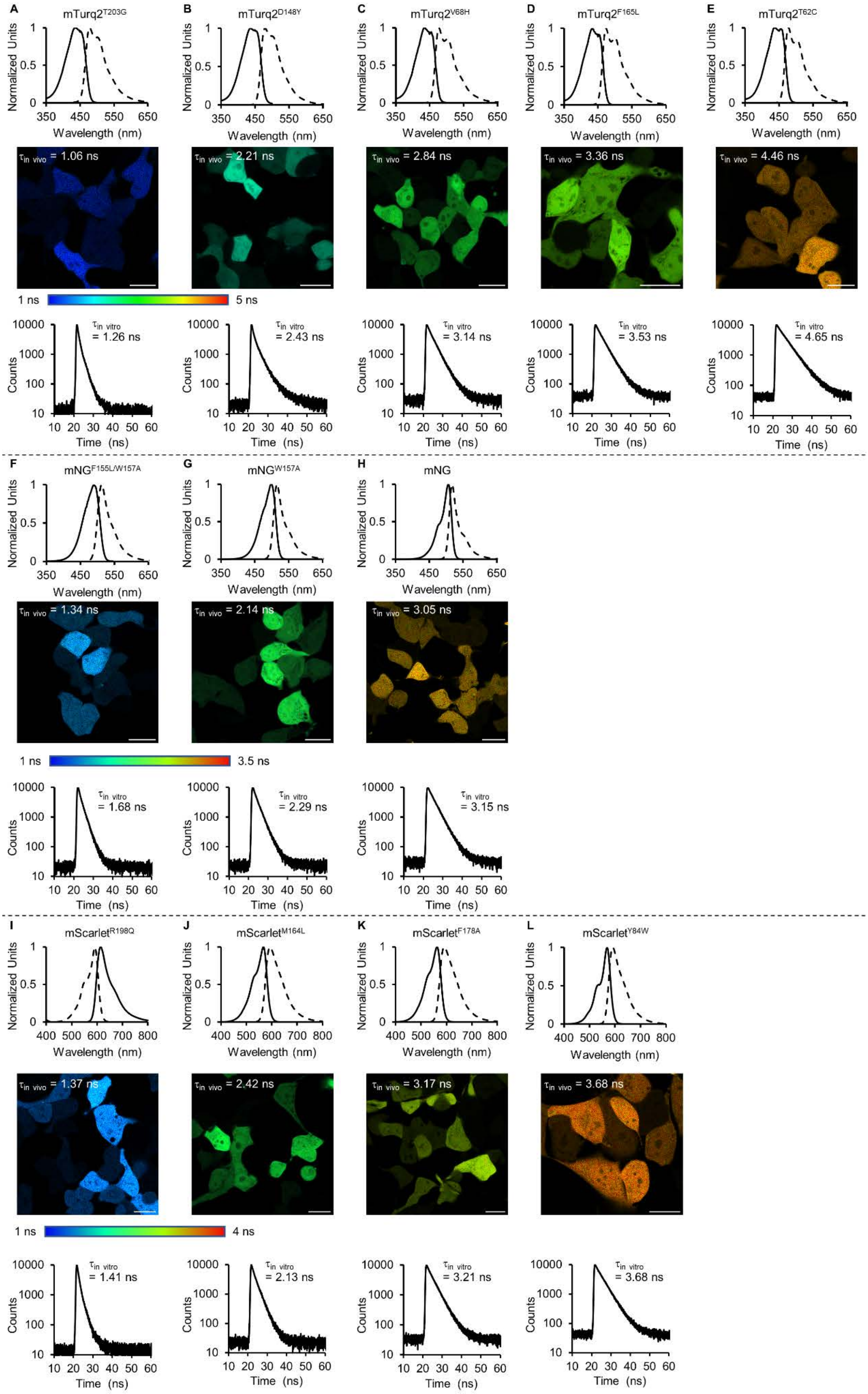


**Figure S2. Fluorescence spectra, TCSPC traces, and FLIM images of TRFPs.**

(A-E) Top panel: excitation (solid line) and emission (dash line) of TRFPs derived from mTurquoise2. Middle panel: FLIM images of TRFPs derived from mTurquoise2 in HEK293T. Bottom panel: decay of fluorescence lifetime of TRFPs derived from mTurquoise2 in the buffer (20 mM Tris, 500 mM NaCl, pH 7.5).
(F-H) Top panel: excitation (solid line) and emission (dash line) of TRFPs derived from mNeonGreen. Middle panel: FLIM images of TRFPs derived from mNeonGreen in HEK293T. Bottom panel: decay of fluorescence lifetime of TRFPs derived from mNeonGreen in the buffer (20 mM Tris, 500 mM NaCl, pH 7.5).
(I-L) Top panel: excitation (solid line) and emission (dash line) of TRFPs derived from mScarlet. Middle panel: FLIM images of TRFPs derived from mScarlet in HEK293T. Bottom panel: decay of fluorescence lifetime of TRFPs derived from mScarlet in the buffer (20 mM Tris, 500 mM NaCl, pH 7.5).
mTurq2 and mNG are the abbreviation of mTurquoise2 and mNeonGreen, respectively. Scale bar, 20 μm.

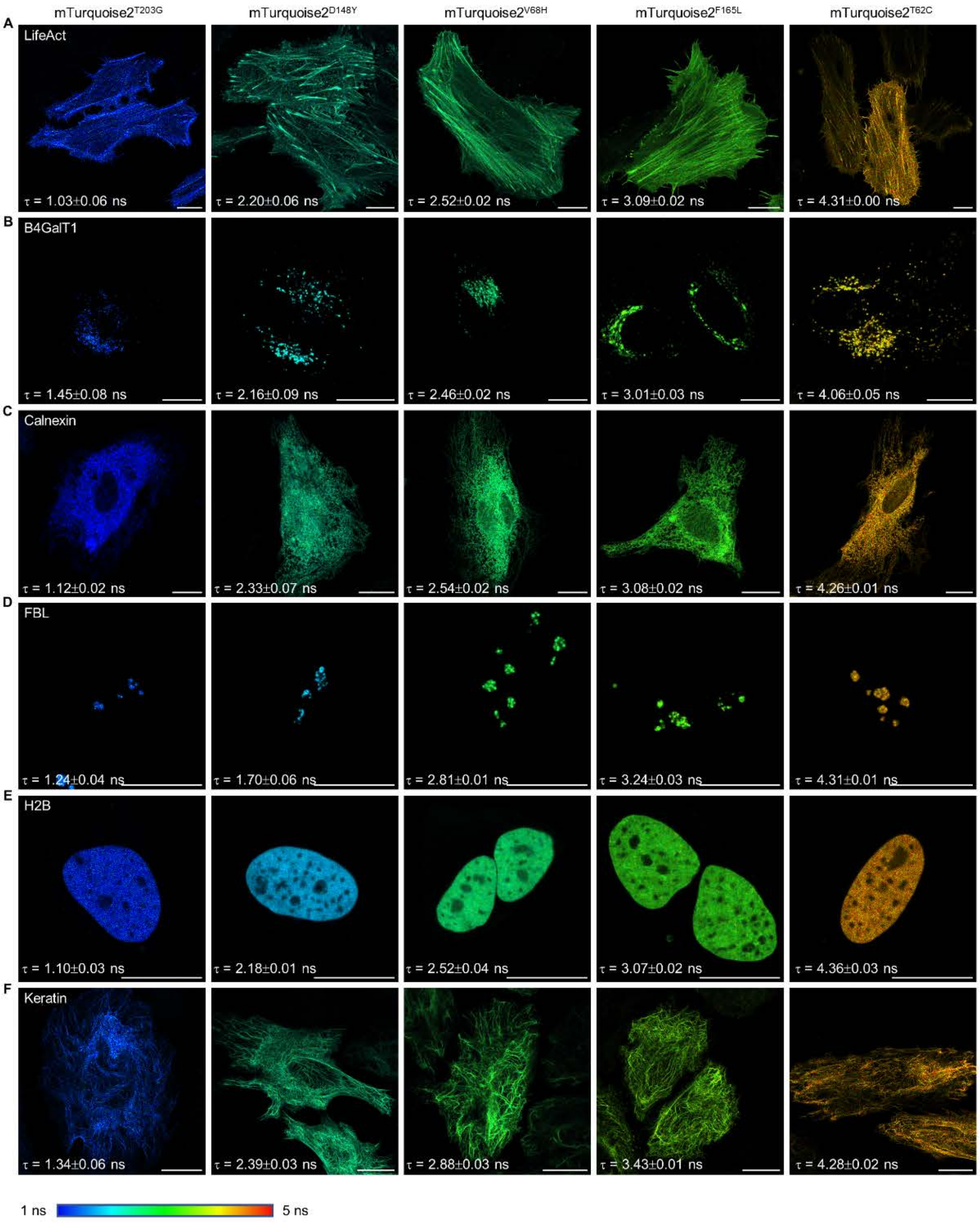


**Figure S3. FLIM images of target proteins fused with mTurquoise2$^{T203G}$, mTurquoise2$^{D148Y}$, mTurquoise2$^{V68H}$, mTurquoise2$^{F165L}$ and mTurquoise2$^{T62C}$.**
(A) Actin filaments, fusing the actin-binding peptide LifeAct.
(B) Golgi apparatus, fusing beta-1,4-galactosidase (B4GalT1).
(C) Endoplasmic reticulum, fusing calnexin.
(D) Nucleoli, fusing Fibrillarin (FBL).
(E) Nucleus, fusing histone 2B (H2B).
(F) Intermediate filaments, fusing keratin 18.
Cells were transiently transfected with the fusion proteins, collecting 500 photons per pixel. Fluorescence lifetime show the average of three independent experiments. Error bar indicate the standard deviation of three independent experiments. Scale bars, 20 μm.

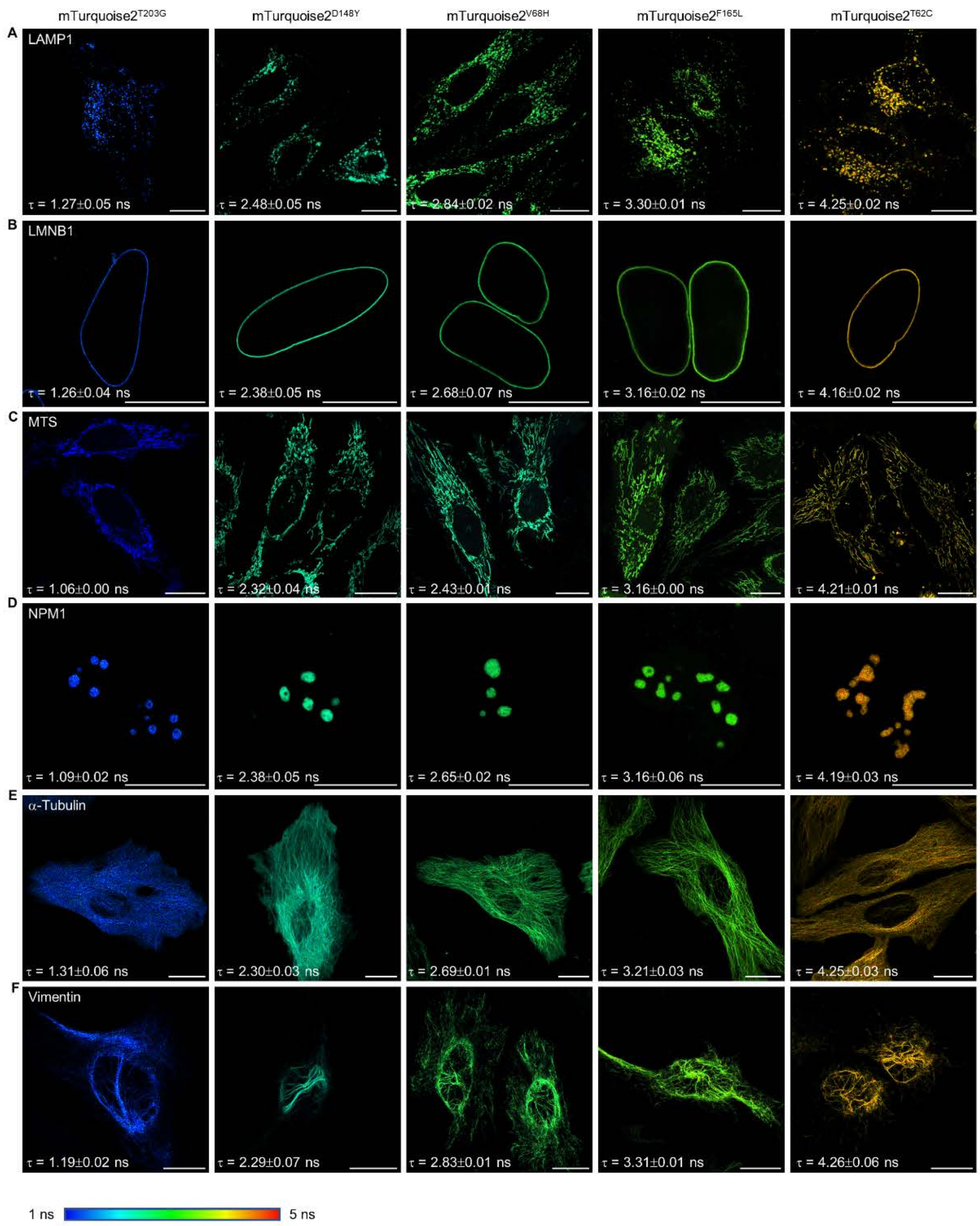


**Figure S4. FLIM images of target proteins fused with mTurquoise2$^{T203G}$, mTurquoise2$^{D148Y}$, mTurquoise2$^{V68H}$, mTurquoise2$^{F165L}$ and mTurquoise2$^{T62C}$ fusion proteins.**

(A) Lysosomes, fusing lysosome associated membrane glycoprotein 1 (LAMP1).

(B) Nuclear membrane, fusing Lamin B1 (LMNB1).

(C) Mitochondria, fusing mitochondrial targeting sequence (MTS), which is the first 29 aa of human cytochrome c oxidase subunit 8A.

(D) Nucleoli, fusing Nucleophosmin 1 (NPM1).

(E) Microtubules, fusing α-tubulin.

(F) Intermediate filaments, fusing vimentin.

Cells were transiently transfected with the fusion proteins, collecting 500 photons per pixel. Fluorescence lifetime show the average of three independent experiments. Error bar indicate the standard deviation of three independent experiments. Scale bars, 20 μm.

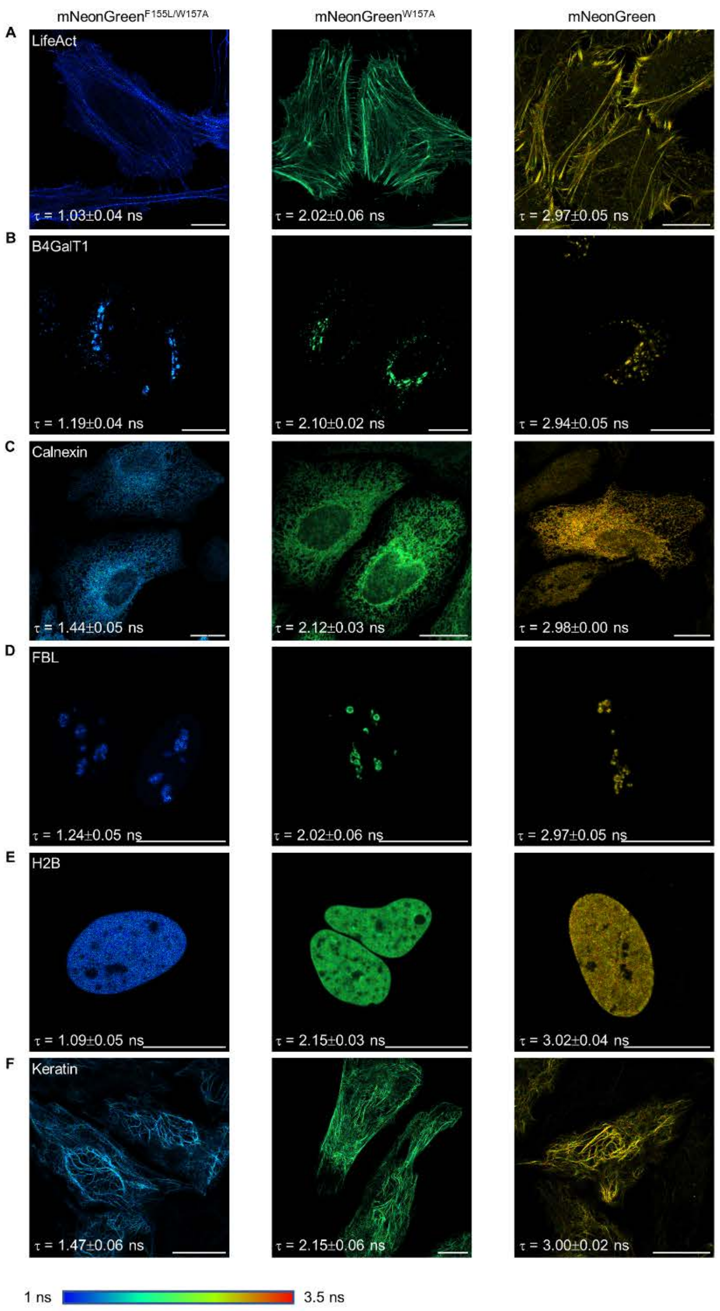


**Figure S5. FLIM images of target proteins fused with mNeonGreen$^{F155L/W157A}$, mNeonGreen$^{W157A}$ and mNeonGreen fusion proteins.**

(A) Actin filaments, fusing the actin-binding peptide LifeAct.

(B) Golgi apparatus, fusing beta-1,4-galactosidase (B4GalT1).

(C) Endoplasmic reticulum, fusing calnexin.

(D) Nucleoli, fusing Fibrillarin (FBL).

(E) Nucleus, fusing histone 2B (H2B).

(F) Intermediate filaments, fusing keratin 18.

Cells were transiently transfected with the fusion proteins, collecting 500 photons per pixel. Fluorescence lifetime show the average of three independent experiments. Error bar indicate the standard deviation of three independent experiments. Scale bars, 20 μm.

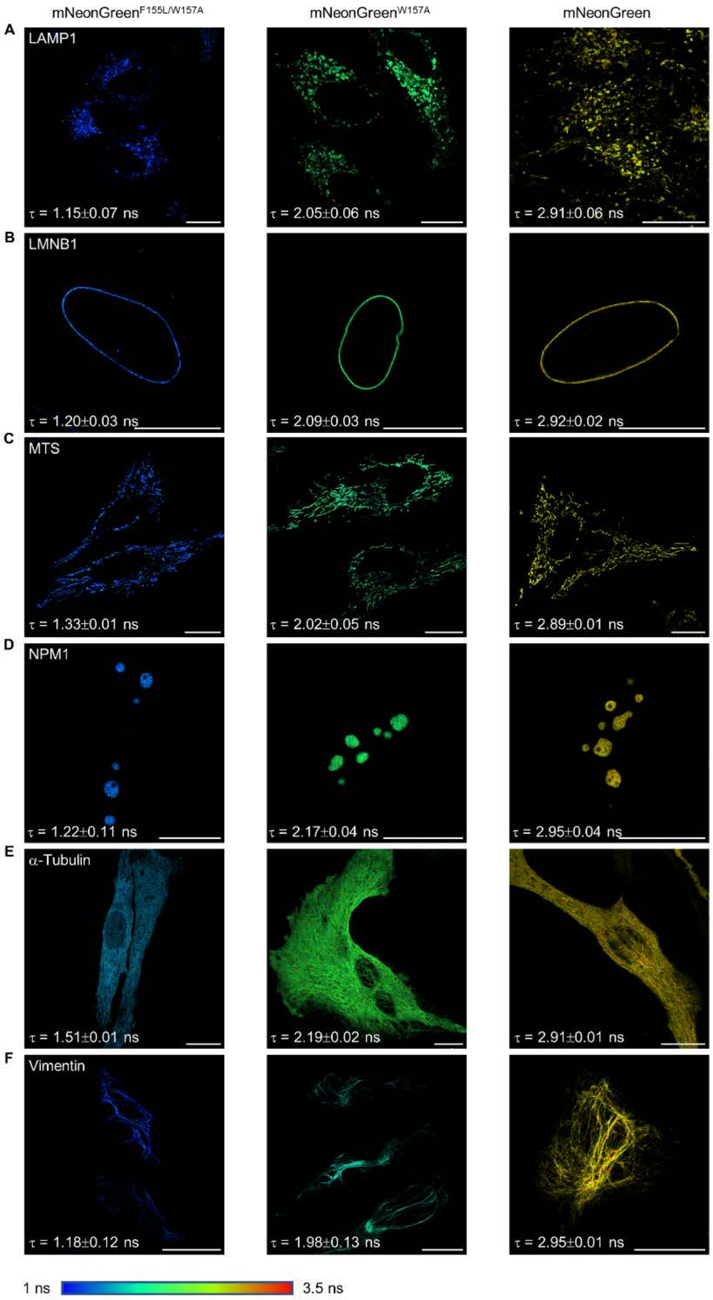


**Figure S6. FLIM images of target proteins fused with mNeonGreen$^{F155L/W157A}$, mNeonGreen$^{W157A}$ and mNeonGreen fusion proteins.**

(A) Lysosomes, fusing lysosome associated membrane glycoprotein 1 (LAMP1).

(B) Nuclear membrane, fusing Lamin B1 (LMNB1).

(C) Mitochondria, fusing mitochondrial targeting sequence (MTS), which is the first 29 aa of human cytochrome c oxidase subunit 8A.

(D) Nucleoli, fusing Nucleophosmin 1 (NPM1).

(E) Microtubules, fusing α-tubulin.

(F) Intermediate filaments, fusing vimentin.

Cells were transiently transfected with the fusion proteins, collecting 500 photons per pixel. Fluorescence lifetime show the average of three independent experiments. Error bar indicate the standard deviation of three independent experiments. Scale bars, 20 μm.

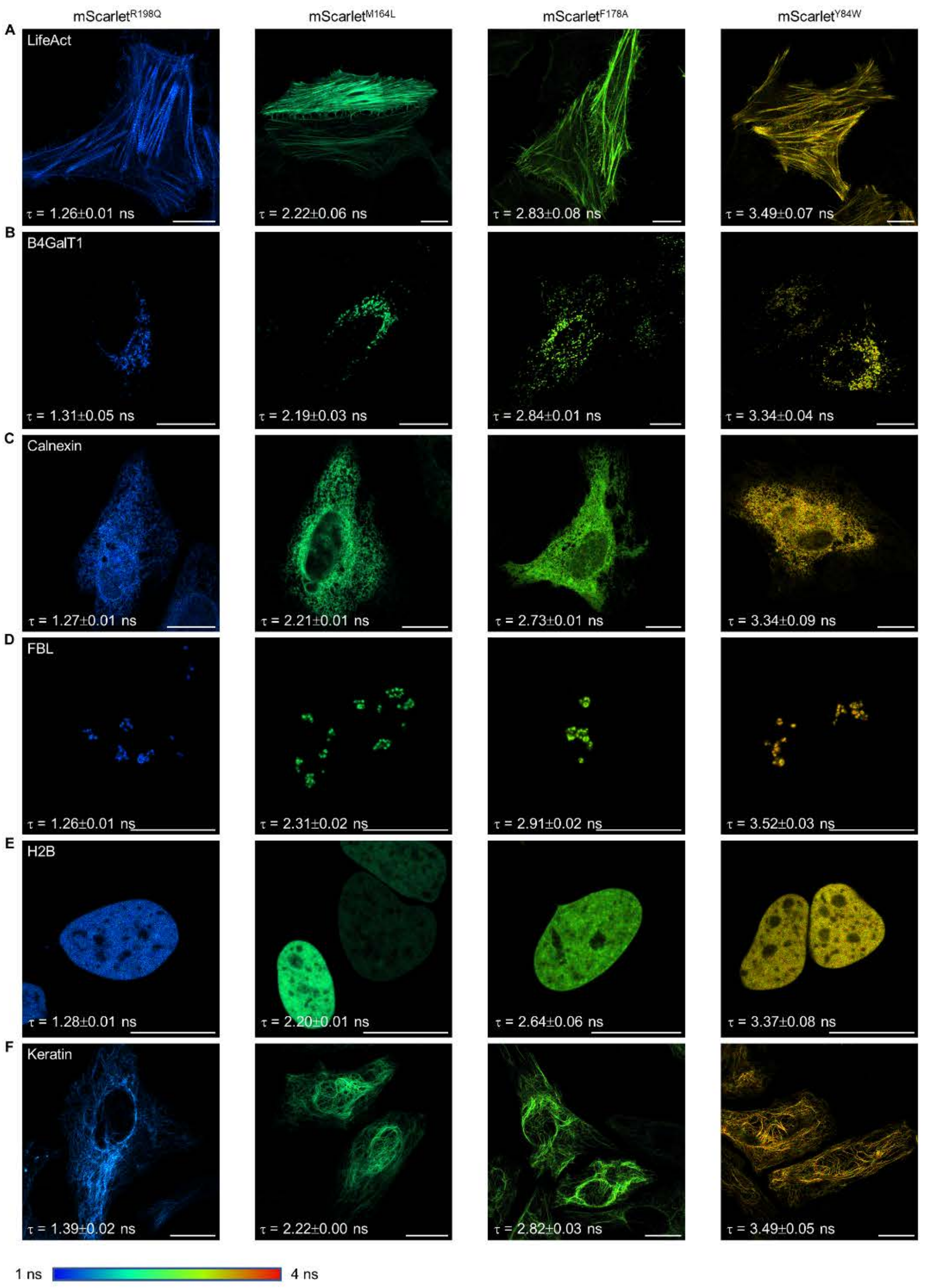


**Figure S7. FLIM images of target proteins fused with mScarlet$^{R198Q}$, mScarlet$^{M164L}$, mScarlet$^{F178A}$ and mScarlet$^{Y84W}$ fusion proteins.**

(A) Actin filaments, fusing the actin-binding peptide LifeAct.

(B) Golgi apparatus, fusing beta-1,4-galactosidase (B4GalT1).

(C) Endoplasmic reticulum, fusing calnexin.

(D) Nucleoli, fusing Fibrillarin (FBL).

(E) Nucleus, fusing histone 2B (H2B).

(F) Intermediate filaments, fusing keratin 18.

Cells were transiently transfected with the fusion proteins, collecting 500 photons per pixel. Fluorescence lifetime show the average of three independent experiments. Error bar indicate the standard deviation of three independent experiments. Scale bars, 20 μm.

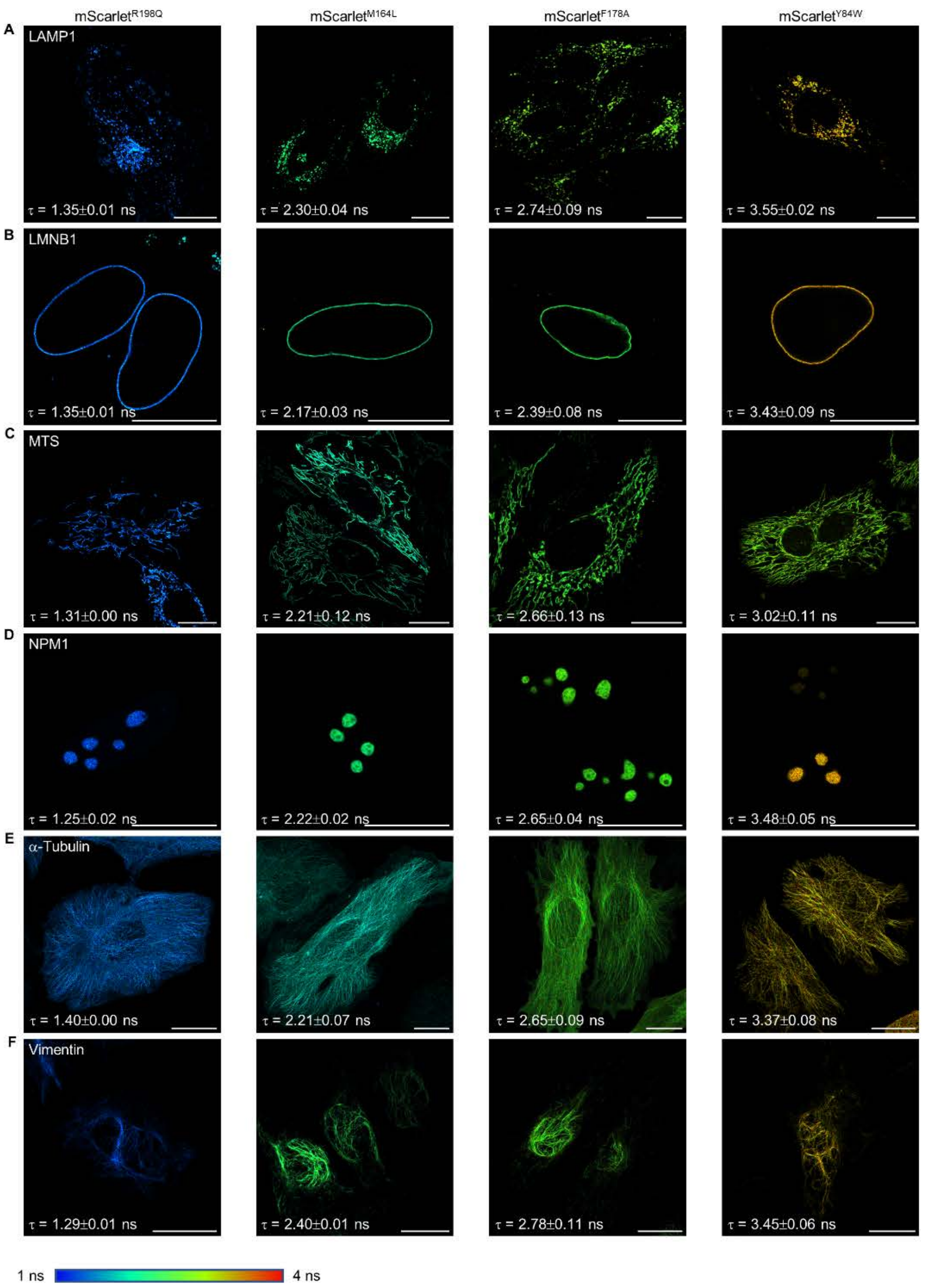


**Figure S8. FLIM images of target proteins fused with mScarlet$^{R198Q}$, mScarlet$^{M164L}$, mScarlet$^{F178A}$ and mScarlet$^{Y84W}$ fusion proteins.**

(A) Lysosomes, fusing lysosome associated membrane glycoprotein 1 (LAMP1).

(B) Nuclear membrane, fusing Lamin B1 (LMNB1).

(C) Mitochondria, fusing mitochondrial targeting sequence (MTS), which is the first 29 aa of human cytochrome c oxidase subunit 8A.

(D) Nucleoli, fusing Nucleophosmin 1 (NPM1).

(E) Microtubules, fusing α-tubulin.

(F) Intermediate filaments, fusing vimentin.

Cells were transiently transfected with the fusion proteins, collecting 500 photons per pixel. Fluorescence lifetime show the average of three independent experiments. Error bar indicate the standard deviation of three independent experiments. Scale bars, 20 μm.

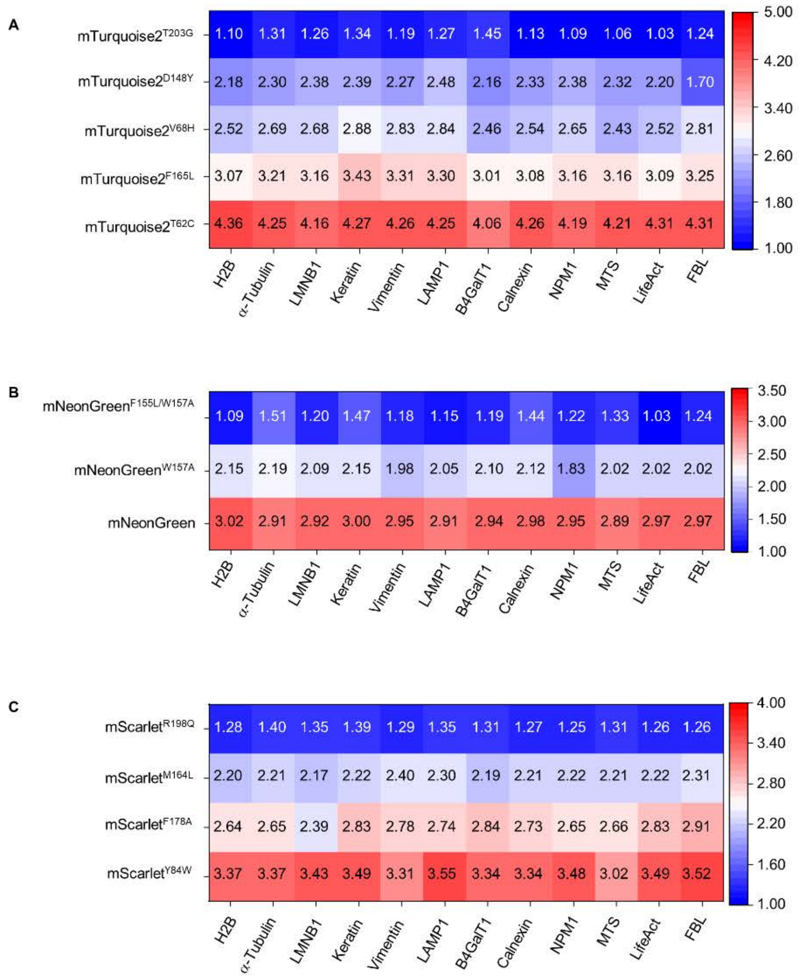


**Figure S9. Summary of fluorescence lifetimes of TRFPs based on mTurquoise2 (A), mNeonGreen (B) and mScarlet (C) fused with different subcellular targets**. Fluorescence lifetime show the average of three independent experiments. Error bar indicate the standard deviation of three independent experiments. Overall fluorescence lifetime was relatively stable over different subcellular targets for different TRFPs.

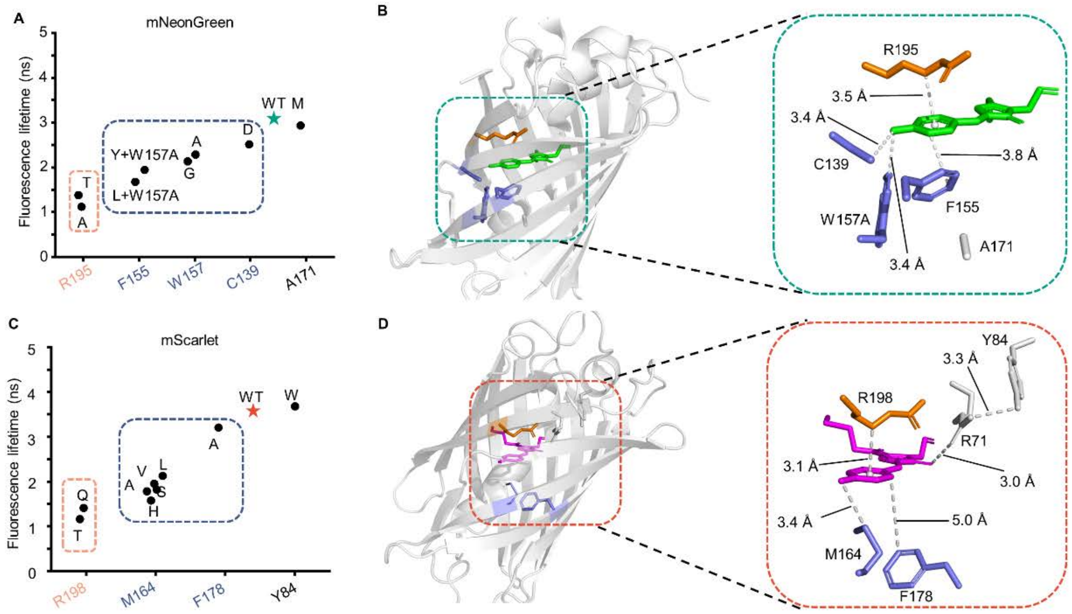


**Figure S10. Structural basis of lifetime regulation for TRFPs based on mNeonGreen and mScarlet.**

(A) Effect of different residues on fluorescence lifetime for mNeonGreen. Orange and blue box indicate the residues which on the top and on the side of phenol, respectively. * indicate the wild type.

(B) Overall structure of mNeonGreen (left panel) and the distance between residues which have effect on the fluorescence lifetime with the chromophore of mNeonGreen (right panel). Orange and blue indicate the residues which on the top and on the side of phenol. PDB ID for mNeonGreen is 5ltp.

(C) Effect of different residues on fluorescence lifetime for mScarlet. Orange and blue box indicate the residues which on the top and on the side of phenol, respectively. * indicate the wild type.

(D) Overall structure of mScarlet (left panel) and the distance between residues which have effect on the fluorescence lifetime with the chromophore of mScarlet (right panel). Orange and blue indicate the residues which on the top and on the side of phenol. PDB ID for mScarlet is 5lk4.

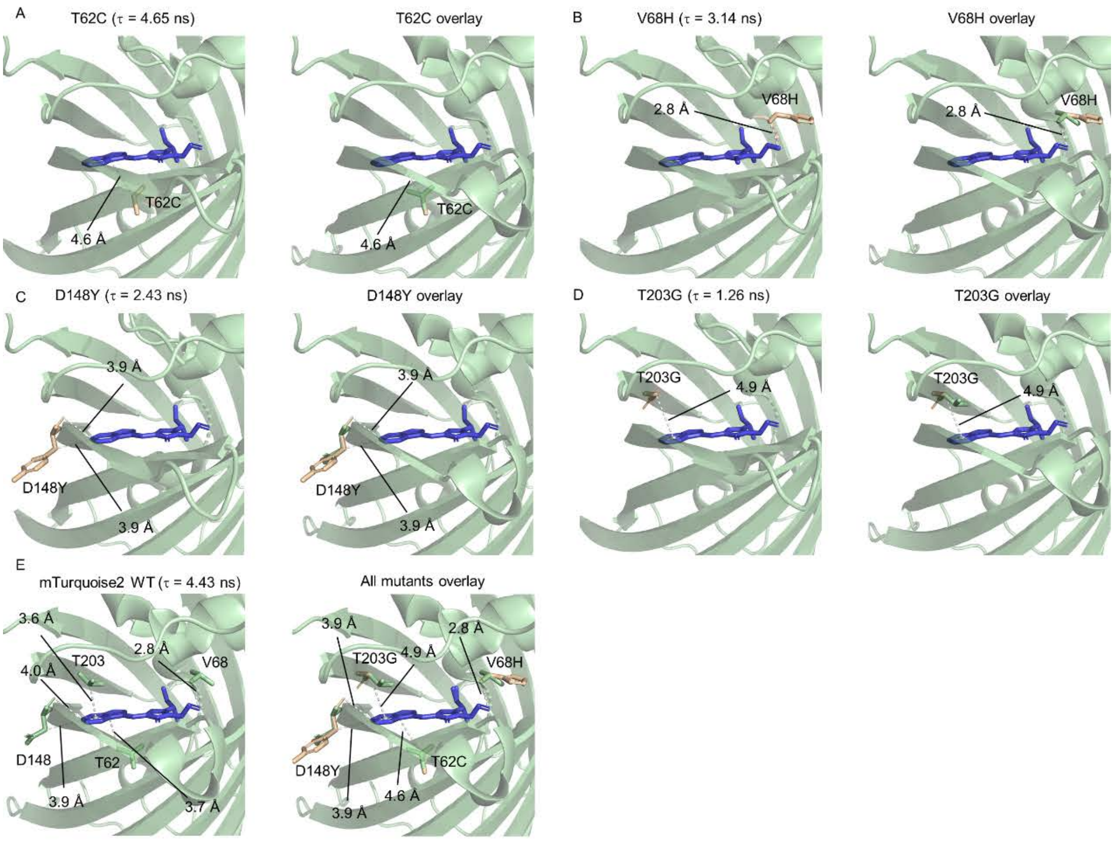


**Figure S11. Structural alignment of mTurquoise2 with its TRFP variants.**

(A-D) The structure of different variants (left panel) and overlay with mTurquoise2 (right panel).

(E) The structure of mTurquoise2 (left panel) and overlay with all variants (right panel). Green and Orange indicated the structure of mTurquoise2 and variants, respectively. The structures of mTurquoise2$^{K206A}$ (PDB ID: 3ztf) was used as the template for the modeling of the variants of mTurquoise2 through Rosetta.

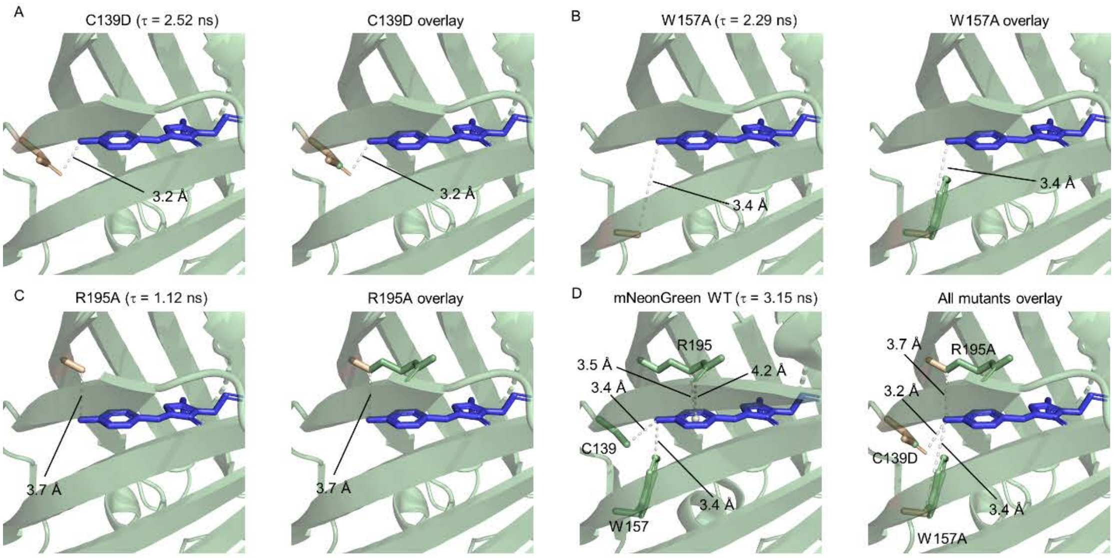


**Figure S12. Structural alignment of mNeonGreen with its TRFP variants.**

(A-C) The structure of different variants (left panel) and overlay with mNeonGreen (right panel).

(D) The structure of mNeonGreen (left panel) and overlay with all variants (right panel). Green and Orange indicated the structure of mNeonGreen and variants, respectively. The structures of mNeonGreen (PDB ID: 5ltp) was used as the template for the modeling of the variants through Rosetta.

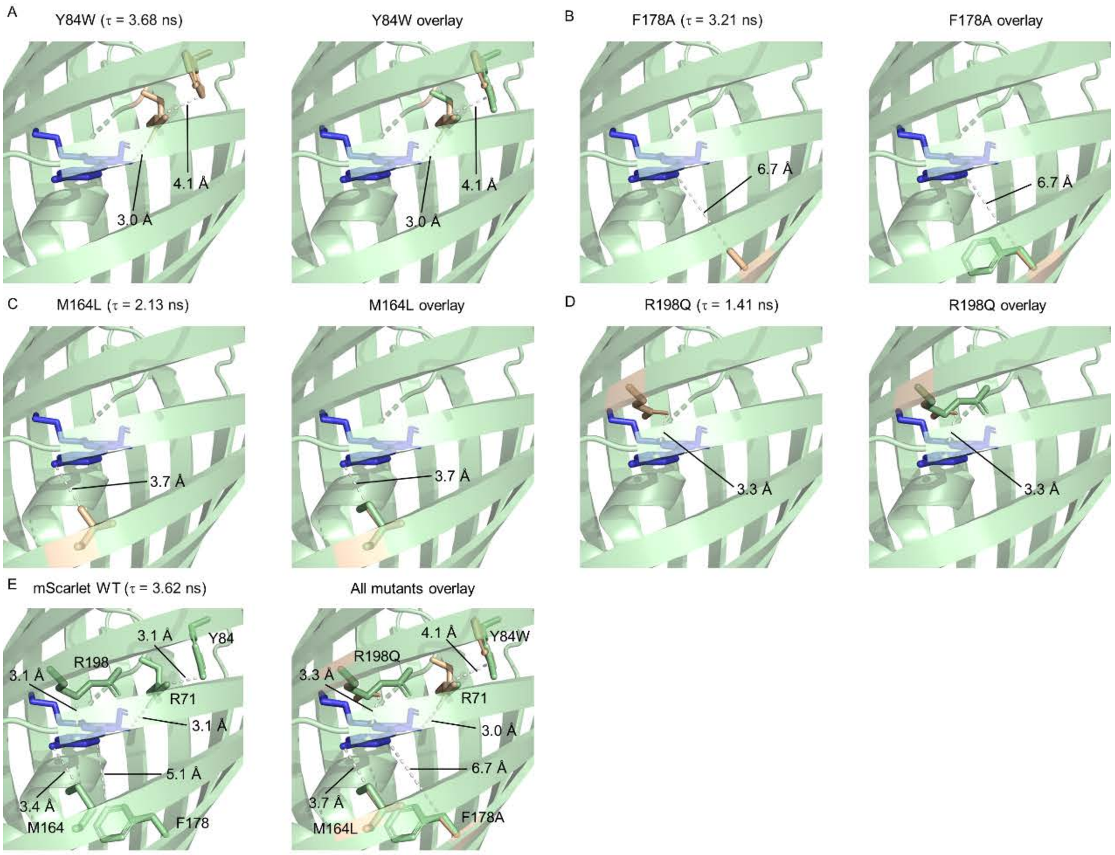


**Figure S13. Structural alignment of mScarlet with its TRFP variants.**

(A-D) The structure of different variants (left panel) and overlay with mScarlet (right panel).

(E) The structure of mScarlet (left panel) and overlay with all variants (right panel).

Green and Orange indicated the structure of mScarlet and variants, respectively. The structures of mScarlet (PDB ID: 5lk4) was used as the template for the modeling of the variants through Rosetta.

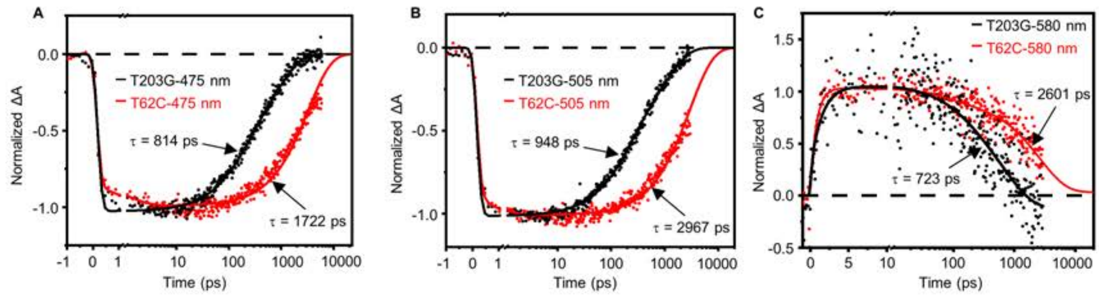


**Figure S14. Kinetic traces of transient absorption of different mTurquoise2 TRFP variants at 475 nm (A), 505 nm (B) and 580 nm (C).** Femtosecond transient absorption was recorded after a pump-laser at 370 nm, which has low energy density (0.23 mJ $cm^{-2}$) to avoid multiphoton process. The fitting curves were based on a global analysis with two (475 nm and 505 nm) or three (580 nm) exponential functions.

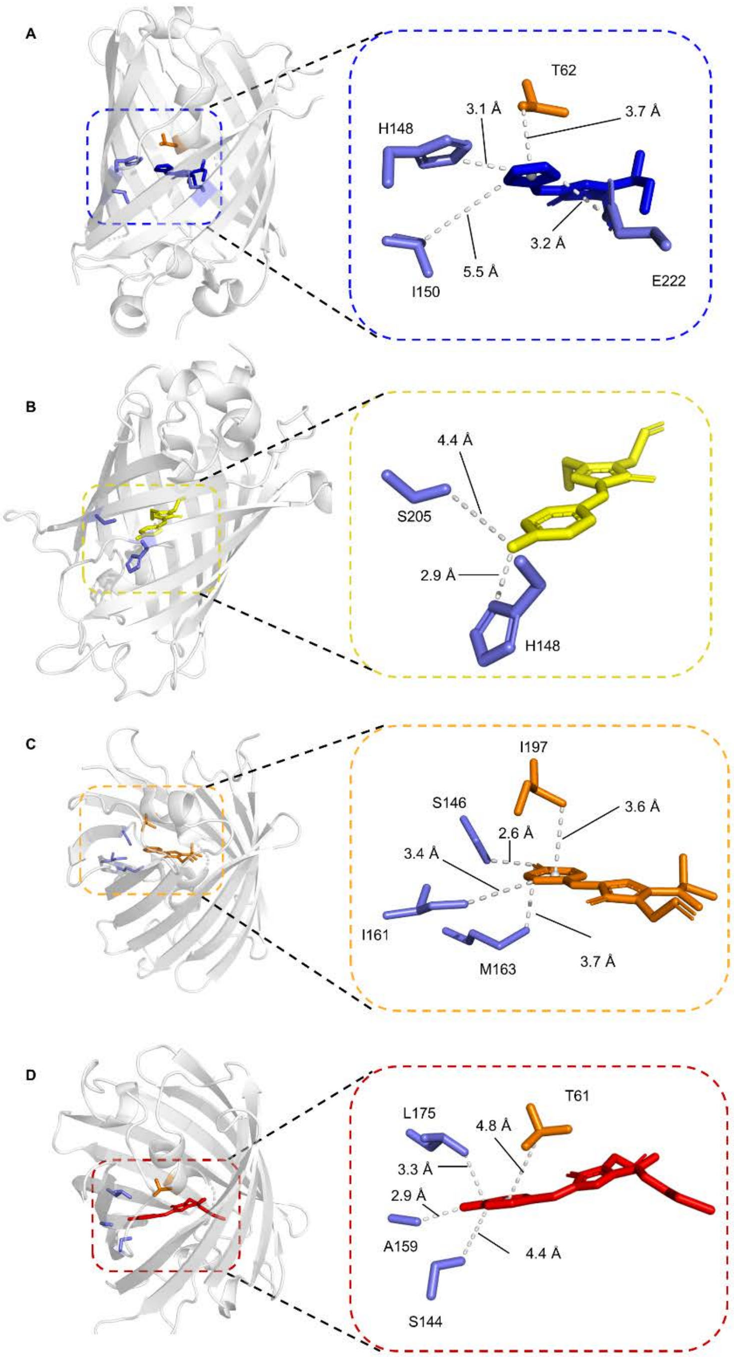


**Figure S15. Structural basis of lifetime control of TRFPs based on EBFP2 (A), mVenus (B), mOrange (C) and mKate2 (D).** Left panel show the overall structure of each protein. Right panel show the distance between these residues which has effect on fluorescence lifetime with chromophore. Orange and blue indicate the residues which on the top and on the side of chromophore, respectively. The structure of EBFP2 and mKate2 was predicted using Rosetta based on the template of BFP (PDB ID: 1bfp) and mKate (PDB ID: 3bxa), respectively. The PDB ID for mVenus and mOrange are 7pnn and 2h5o, respectively.

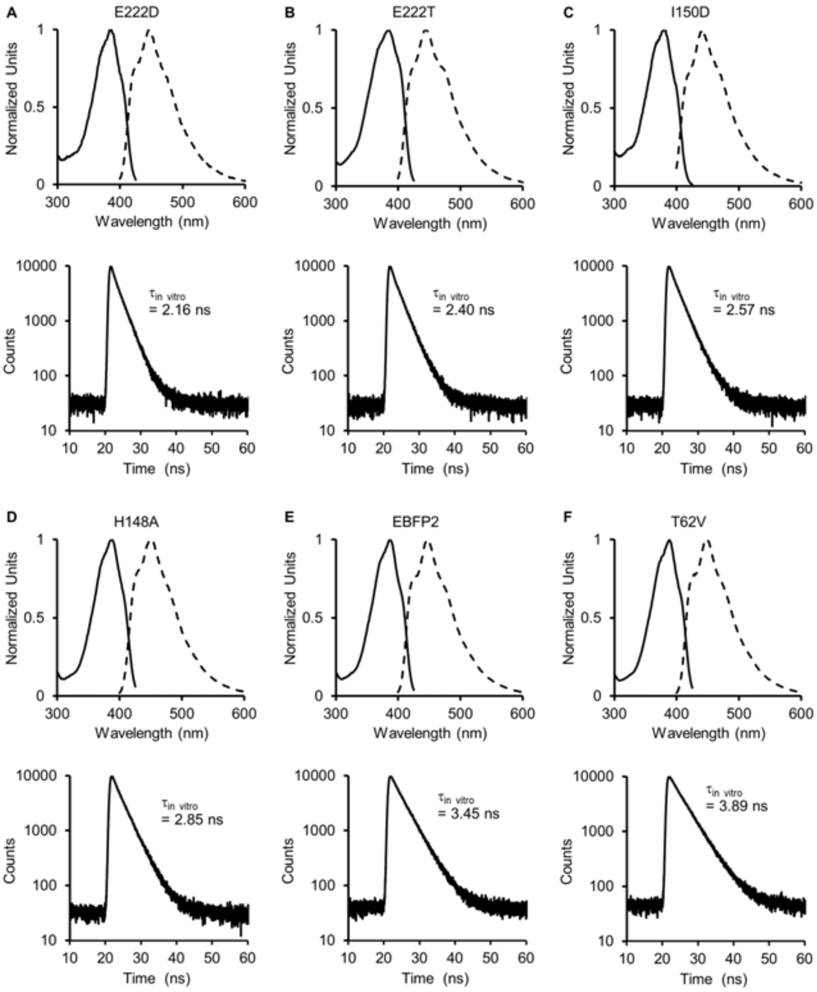


**Figure S16. Fluorescence spectra and TCSPC traces of TRFP based on EBFP2.**

Top panel: excitation (solid line) and emission (dash line) of EBFP2$^{E222D}$ (A), EBFP2$^{E222T}$ (B), EBFP2$^{I150D}$ (C), EBFP2$^{H148A}$ (D), EBFP2 (E) and EBFP2$^{T62V}$ (F) in the buffer (20 mM Tris, 500 mM NaCl, pH 7.5).

Bottom panel: decay of fluorescence lifetime of EBFP2$^{E222D}$ (A), EBFP2$^{E222T}$ (B), EBFP2$^{I150D}$ (C), EBFP2$^{H148A}$ (D), EBFP2 (E) and EBFP2$^{T62V}$ (F) in the buffer (20 mM Tris, 500 mM NaCl, pH 7.5).

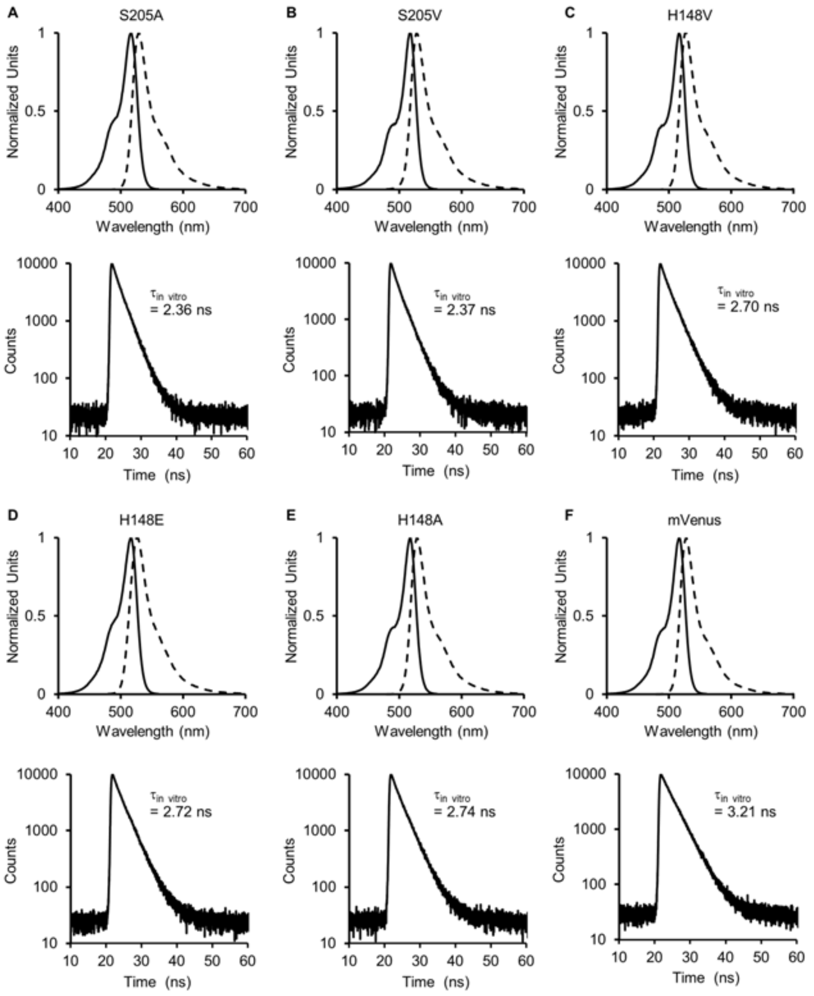


**Figure S17. Fluorescence spectra and TCSPC traces of TRFP based on mVenus.**

Top panel: excitation (solid line) and emission (dash line) of mVenus$^{S205A}$ (A), mVenus$^{S205V}$ (B), mVenus$^{H148V}$ (C), mVenus$^{H148E}$ (D), mVenus$^{H148A}$ (E) and mVenus (F) in the buffer (20 mM Tris, 500 mM NaCl, pH 7.5).

Bottom panel: decay of fluorescence lifetime of mVenus$^{S205A}$ (A), mVenus$^{S205V}$ (B), mVenus$^{H148V}$ (C), mVenus$^{H148E}$ (D), mVenus$^{H148A}$ (E) and mVenus (F) in the buffer (20 mM Tris, 500 mM NaCl, pH 7.5).

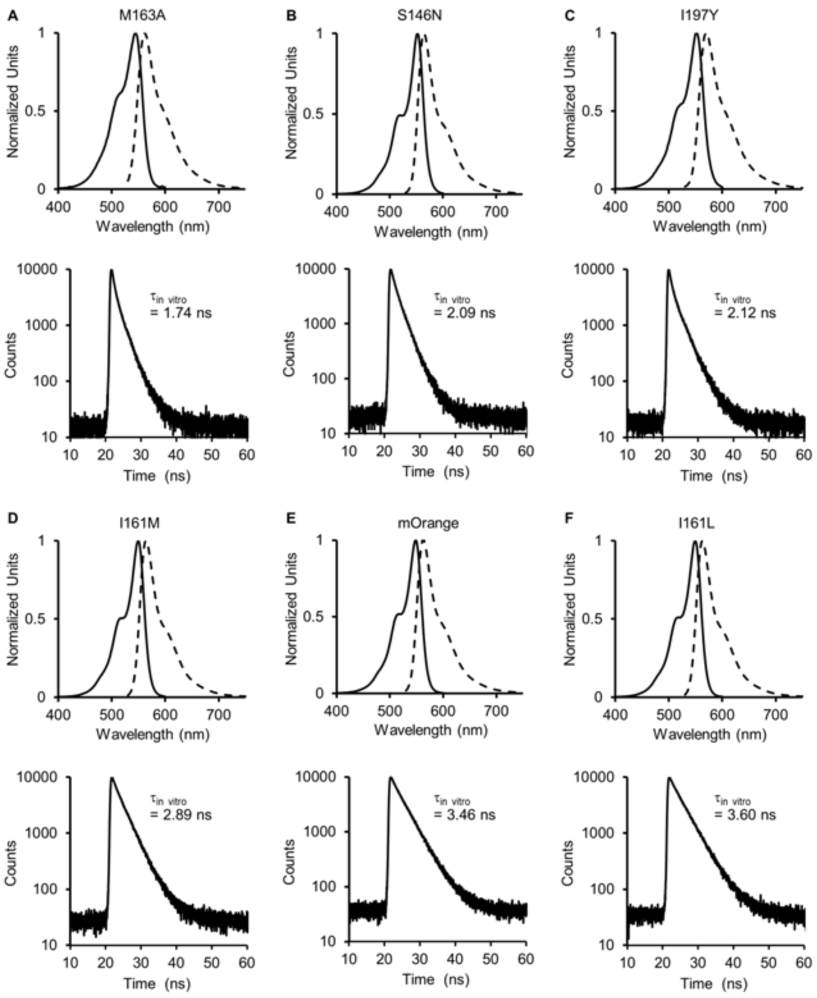


**Figure S18. Fluorescence spectra and TCSPC traces of TRFP based on mOrange.**
Top panel: excitation (solid line) and emission (dash line) of mOrange$^{M163A}$ (A), mOrange$^{S146N}$ (B), mOrange$^{I197Y}$ (C), mOrange$^{I161M}$ (D), mOrange (E) and mOrange$^{I161L}$ (F) in the buffer (20 mM Tris, 500 mM NaCl, pH 7.5).
Bottom panel: decay of fluorescence lifetime of mOrange$^{M163A}$ (A), mOrange$^{S146N}$ (B), mOrange$^{I197Y}$ (C), mOrange$^{I161M}$ (D), mOrange (E) and mOrange$^{I161L}$ (F) in the buffer (20 mM Tris, 500 mM NaCl, pH 7.5).

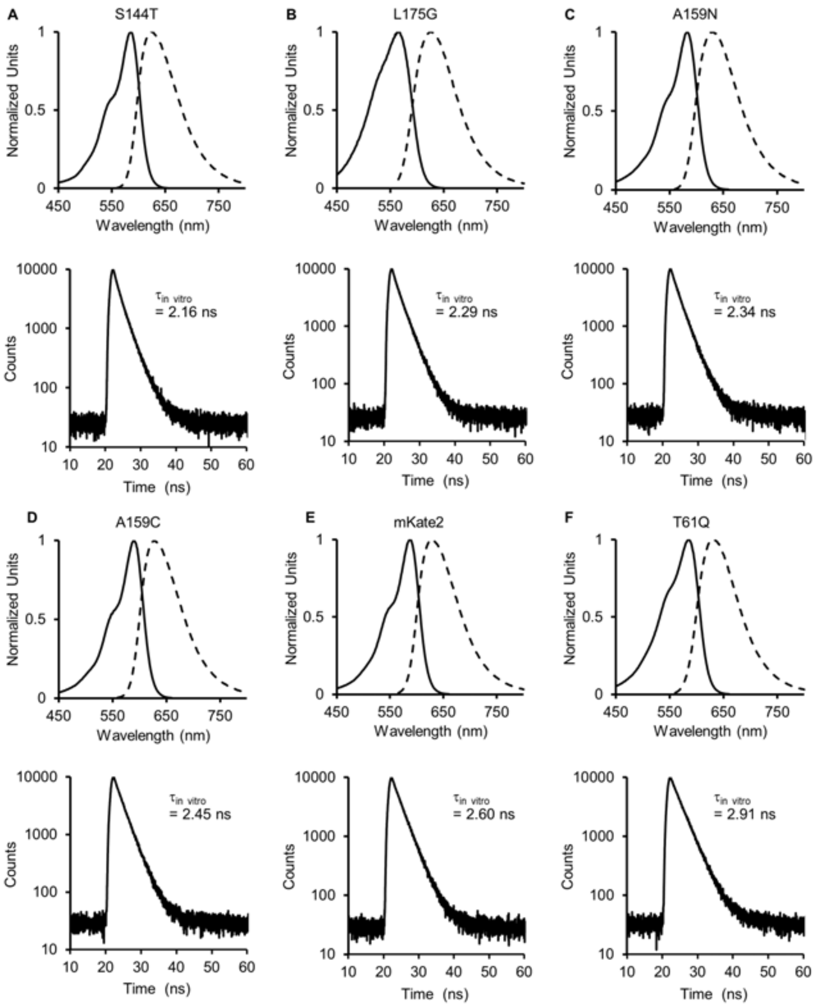


**Figure S19. Fluorescence spectra and TCSPC traces of TRFP based on mKate2.**
Top panel: excitation (solid line) and emission (dash line) of mKate2$^{S144T}$ (A), mKate2$^{L175G}$ (B), mKate2$^{A159N}$ (C), mKate2$^{A159C}$ (D), mKate2 (E) and mKate2$^{T61Q}$ (F) in the buffer (20 mM Tris, 500 mM NaCl, pH 7.5).
Bottom panel: decay of fluorescence lifetime of mKate2$^{S144T}$ (A), mKate2$^{L175G}$ (B), mKate2$^{A159N}$ (C), mKate2$^{A159C}$ (D), mKate2 (E) and mKate2$^{T61Q}$ (F) in the buffer (20 mM Tris, 500 mM NaCl, pH 7.5).

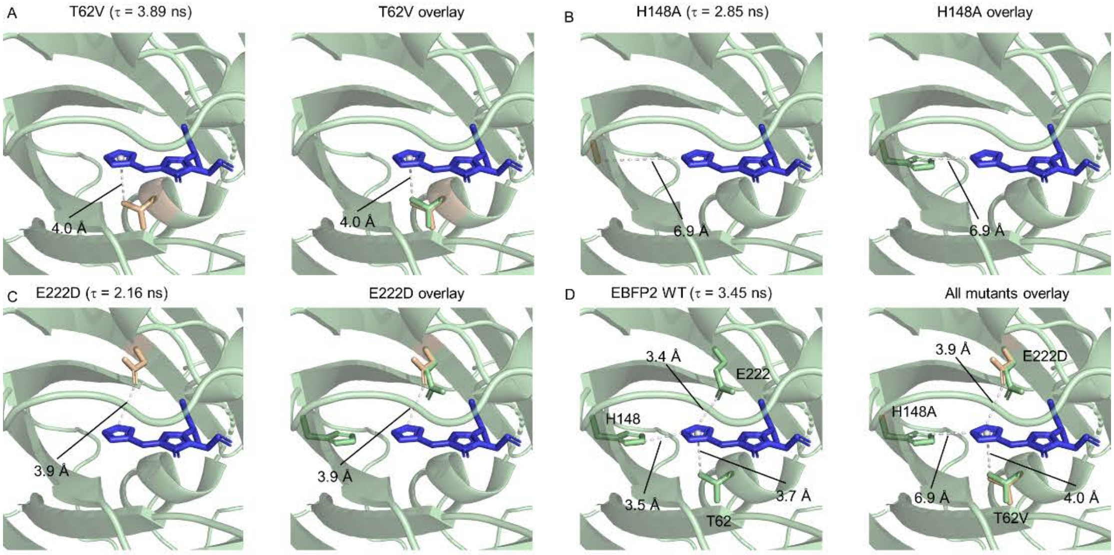


**Figure S20. Structural alignment of EBFP2 and its TRFP variants.**

(A-C) The structure of different variants (left panel) and overlay with EBFP2 (right panel).

(D) The structure of EBFP2 (left panel) and overlay with all variants (right panel).

Green and Orange indicated the structure of EBFP2 and variants, respectively. The structures of BFP (PDB ID: 1bfp) was used as the template for the modeling of the variants through Rosetta.

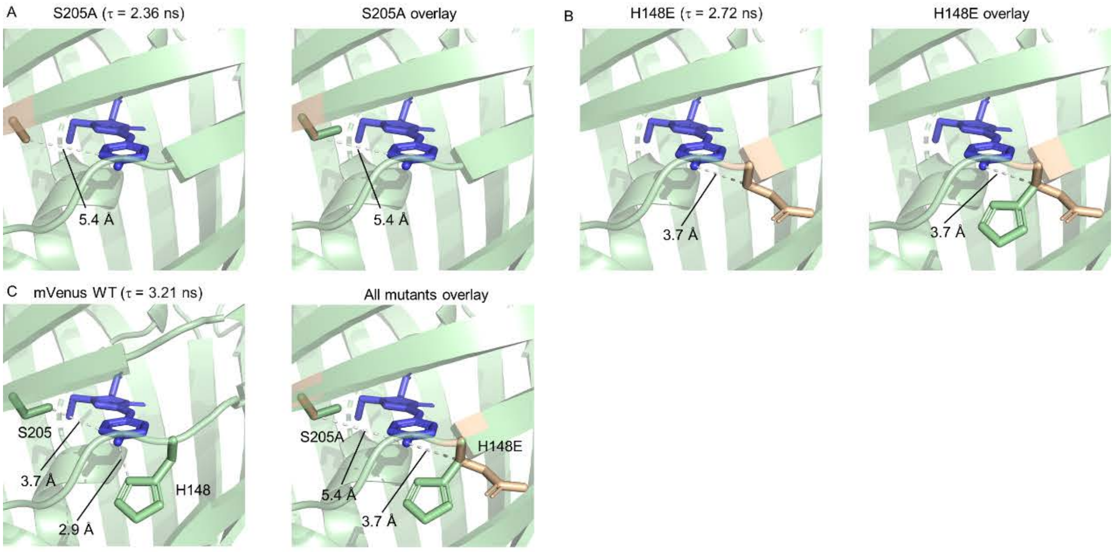


**Figure S21. Structural alignment of mVenus and its TRFP variants.**

(A and B) The structure of different variants (left panel) and overlay with mVenus (right panel).

(C) The structure of mVenus (left panel) and overlay with all variants (right panel).

Green and Orange indicated the structure of mVenus and variants, respectively. The structures of mVenus (PDB ID: 7pnn) was used as the template for the modeling of the variants through Rosetta.

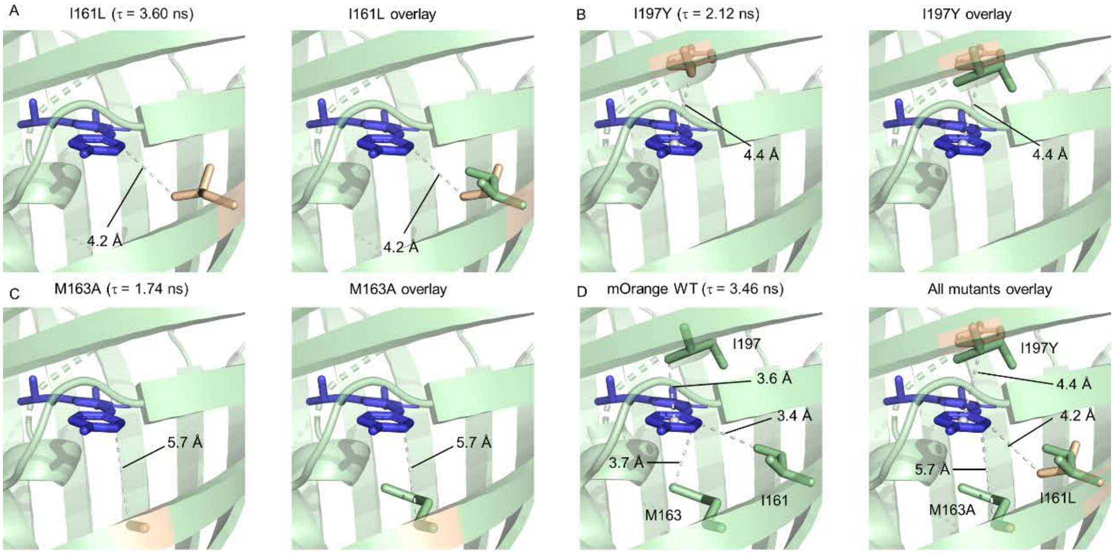


**Figure S22. Structural alignment of mOrange and its TRFP variants.**

(A-C) The structure of different variants (left panel) and overlay with mOrange (right panel).

(D) The structure of mOrange (left panel) and overlay with all variants (right panel). Green and Orange indicated the structure of mOrange and variants, respectively. The structures of mOrange (PDB ID: 2h5o) was used as the template for the modeling of the variants through Rosetta.

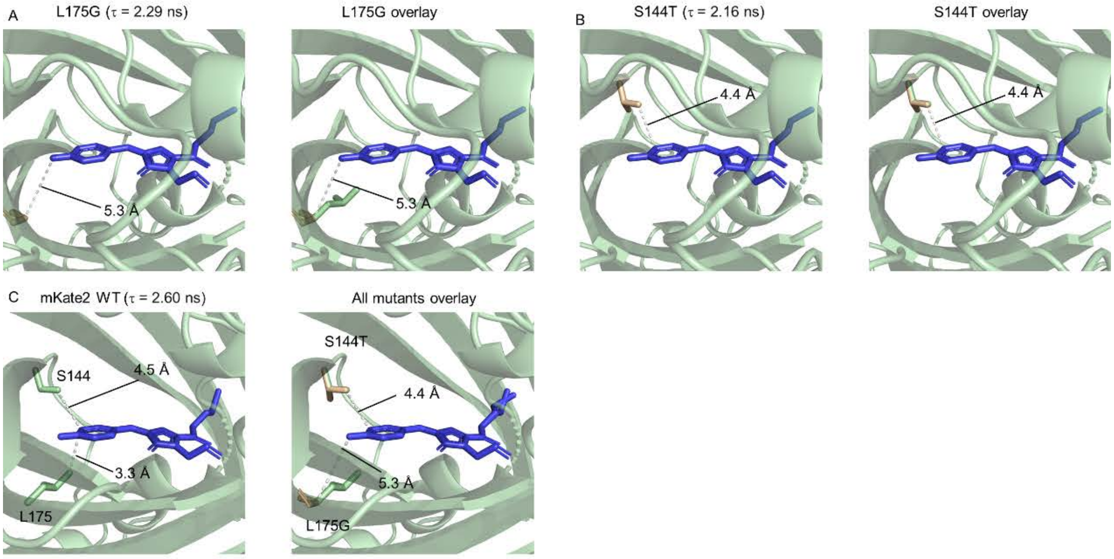


**Figure S23. Structural alignment of mKate2 and its TRFP variants.**

(A-C) The structure of different variants (left panel) and overlay with mKate2 (right panel).

(D) The structure of mKate2 (left panel) and overlay with all variants (right panel). Green and Orange indicated the structure of mKate2 and variants, respectively. The structures of mKate (PDB ID: 3bxa) was used as the template for the modeling of the variants through Rosetta.

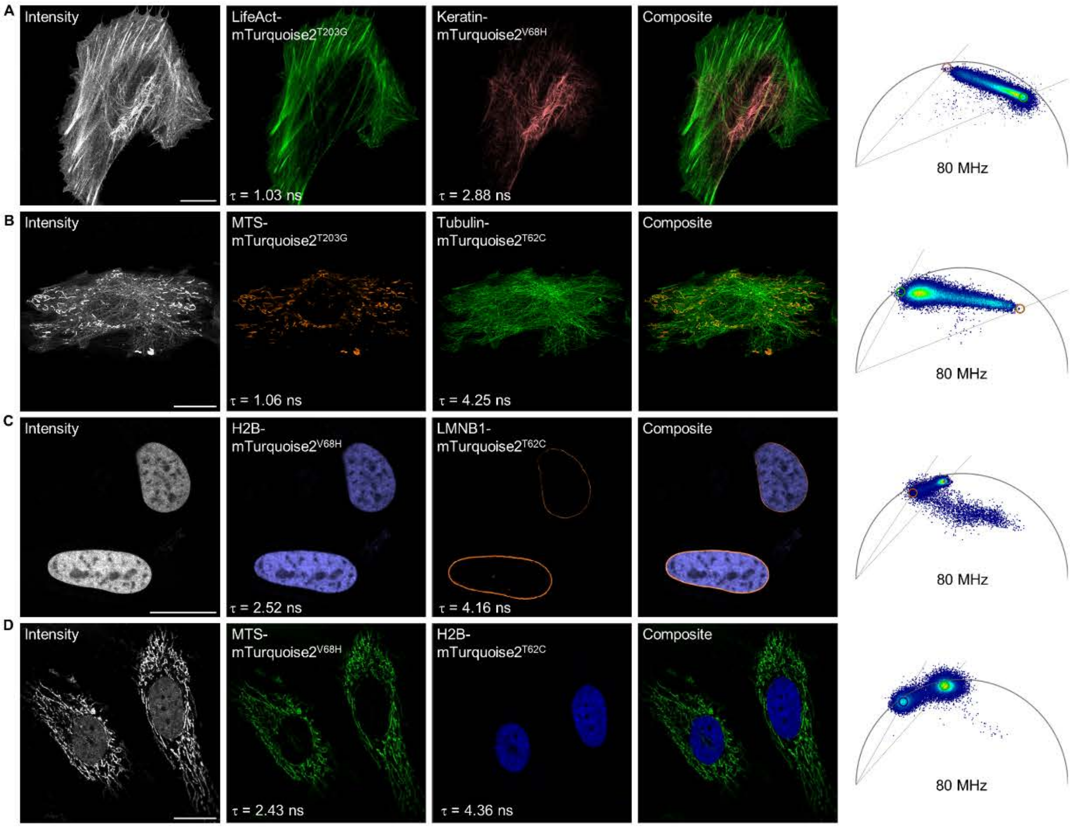


**Figure S24. Multiplexing imaging of two mTurquoise2 TRFPs.**

(A) LifeAct-mTurquoise2$^{T203G}$ targeting mitochondria and Keratin-mTurquoise2$^{V68H}$ targeting intermediate filament.

(B) MTS-mTurquoise2$^{T203G}$ targeting mitochondria and Tubulin-mTurquoise2$^{T62C}$ targeting microtubules.

(C) H2B-mTurquoise2$^{V68H}$ targeting nucleus and LMNB1-mTurquoise2$^{T62C}$ targeting nuclear membrane.

(D) MTS-mTurquoise2$^{V68H}$ targeting mitochondria and H2B-mTurquoise2$^{T62C}$ targeting nucleus.

The fluorescence intensity, the composite, the two individual separated species, the fluorescence lifetime of each fusion proteins as well as the corresponding phasor plot used for separation are given. Scale bars, 20 μm.

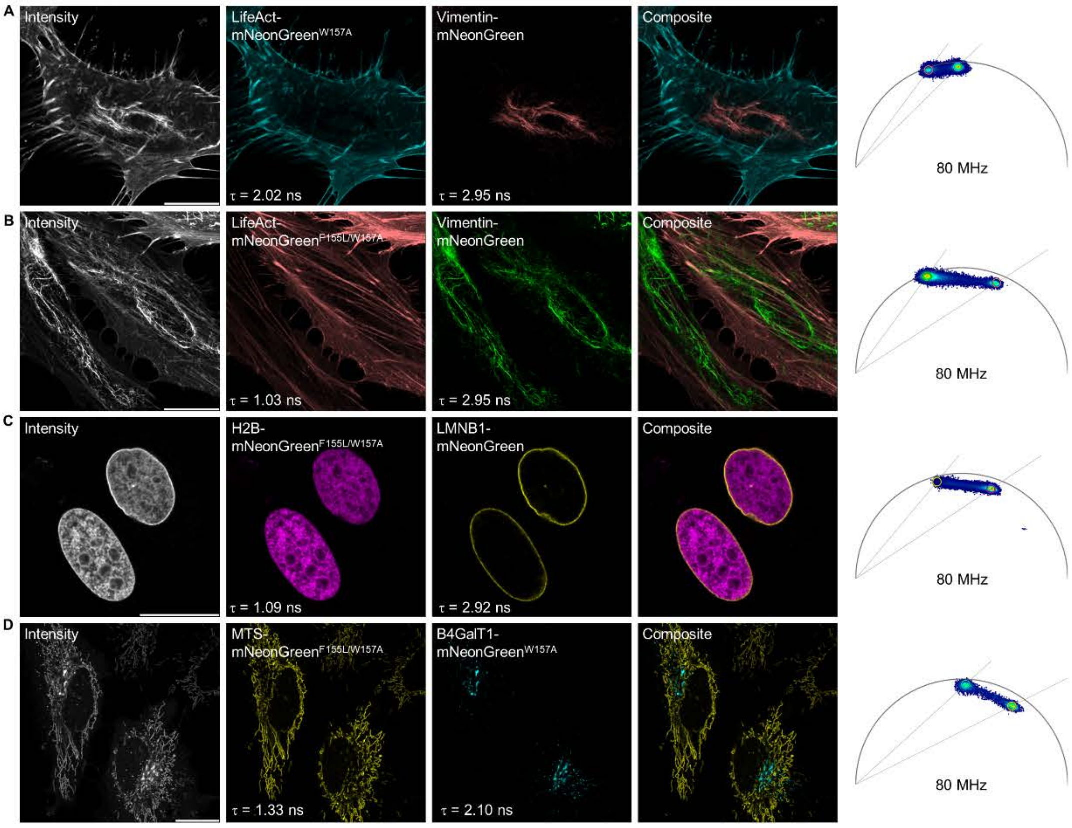


**Figure S25. Multiplexing imaging of two mNeonGreen TRFPs.**

(A) LifeAct-mNeonGreen$^{W157A}$ targeting actin filaments and Vimentin-mNeonGreen targeting intermediate filaments.

(B) LifeAct-mNeonGreen$^{F155L/W157A}$ targeting actin filaments and Vimentin-mNeonGreen targeting intermediate filaments.

(C) H2B-mNeonGreen$^{F155L/W157A}$ targeting nucleus and LMNB1-mNeonGreen targeting nuclear membrane.

(D) MTS-mNeonGreen$^{F155L/W157A}$ targeting mitochondria and B4GalT1-mNeonGreen$^{W157A}$ targeting Golgi apparatus.

The fluorescence intensity, the composite, the two individual separated species, the fluorescence lifetime of each fusion proteins as well as the corresponding phasor plot used for separation are given. Scale bars, 20 μm.

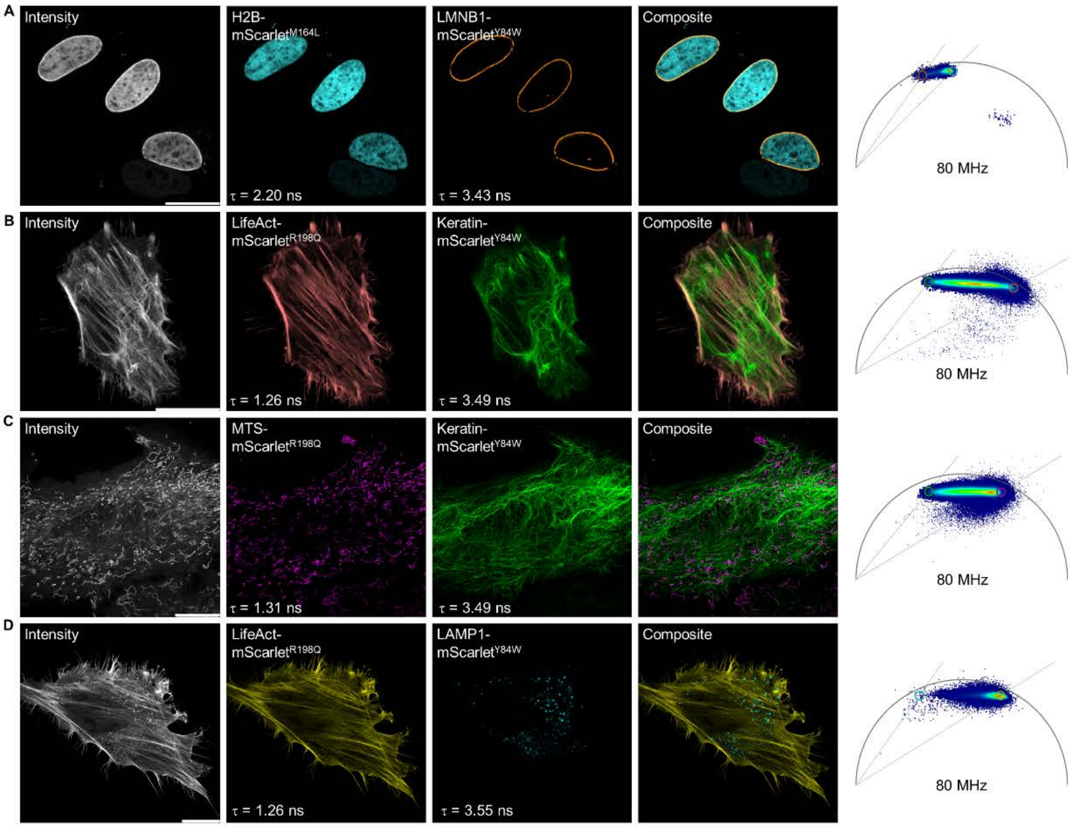


**Figure S26. Multiplexing imaging of two mScarlet TRFPs.**
(A) H2B-mScarlet$^{M164L}$ targeting nucleus and LMNB1-mScarlet$^{Y84W}$ targeting nuclear membrane.
(B) LifeAct-mScarlet$^{R198Q}$ targeting actin filaments and Keratin-mScarlet$^{Y84W}$ targeting intermediate filaments.
(C) MTS-mScarlet$^{R198Q}$ targeting mitochondria and Keratin-mScarlet$^{Y84W}$ targeting intermediate filaments.
(D) LifeAct-mScarlet$^{R198Q}$ targeting actin filaments and LAMP1-mScarlet$^{Y84W}$ targeting lysosomes.
The fluorescence intensity, the composite, the two individual separated species, the fluorescence lifetime of each fusion proteins as well as the corresponding phasor plot used for separation are given. Scale bars, 20 μm.

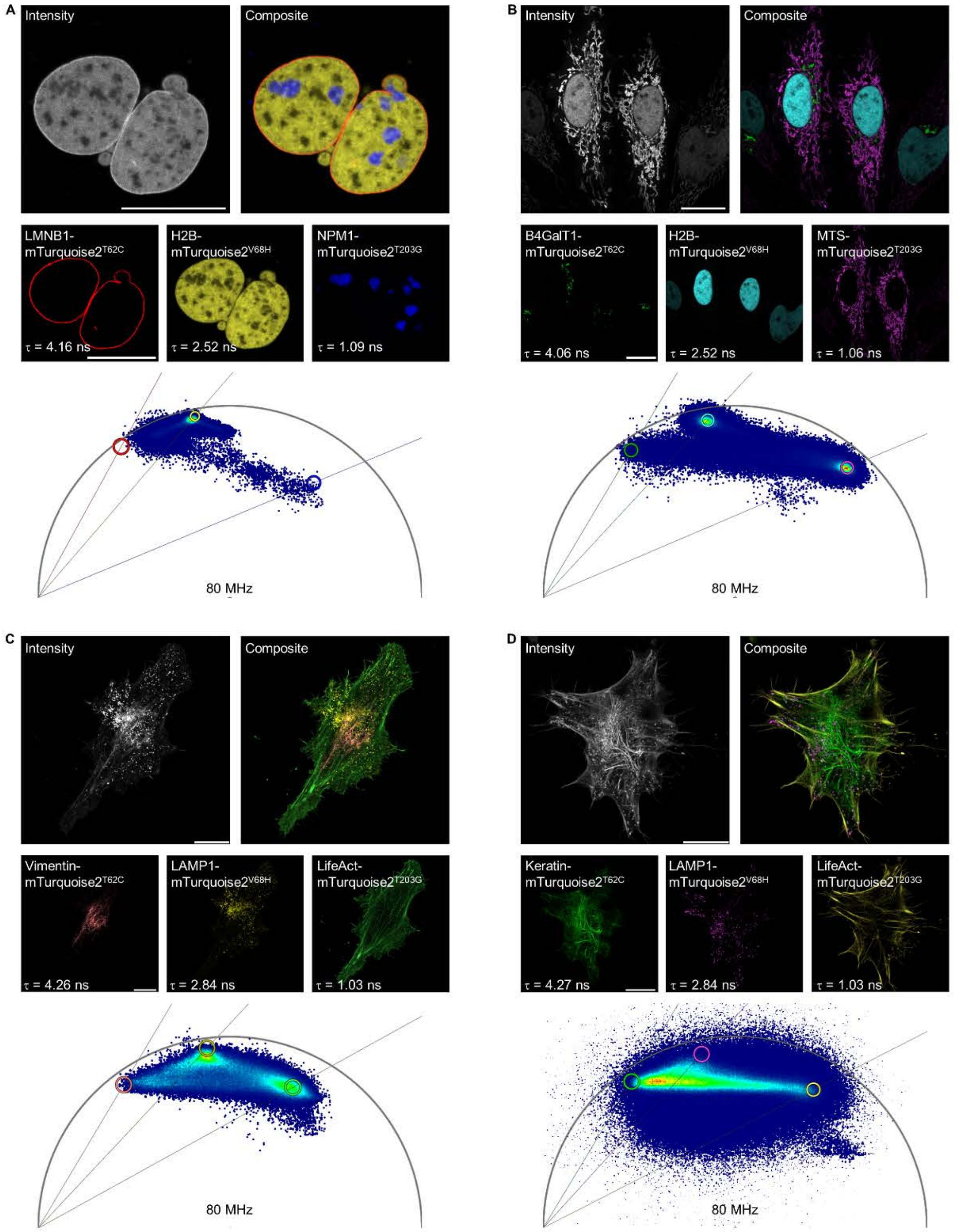


**Figure S27. Multiplexing imaging of three mTurquoise2 TRFPs.**

(A) NPM1-mTurquoise2$^{T203G}$ targeting nucleoli, H2B-mTurquoise2$^{V68H}$ targeting nucleus and LMNB1-mTurquoise2$^{T62C}$ targeting nuclear membrane.

(B) MTS-mTurquoise2$^{T203G}$ targeting mitochondria, H2B-mTurquoise2$^{V68H}$ targeting nucleus and B4GalT1-mTurquoise2$^{T62C}$ targeting Golgi apparatus.

(C) LifeAct-mTurquoise2$^{T203G}$ targeting actin filaments, LAMP1-mTurquoise2$^{V68H}$ targeting lysosomes and Vimentin-mTurquoise2$^{T62C}$ targeting intermediate filaments.

(D) LifeAct-mTurquoise2$^{T203G}$ targeting actin filaments, LAMP1-mTurquoise2$^{V68H}$ targeting lysosomes and Keratin-mTurquoise2$^{T62C}$ targeting intermediate filaments.

The fluorescence intensity, the composite, the three individual separated species, the fluorescence lifetime of each fusion proteins as well as the corresponding phasor plot used for separation are given. Scale bars, 20 µm.

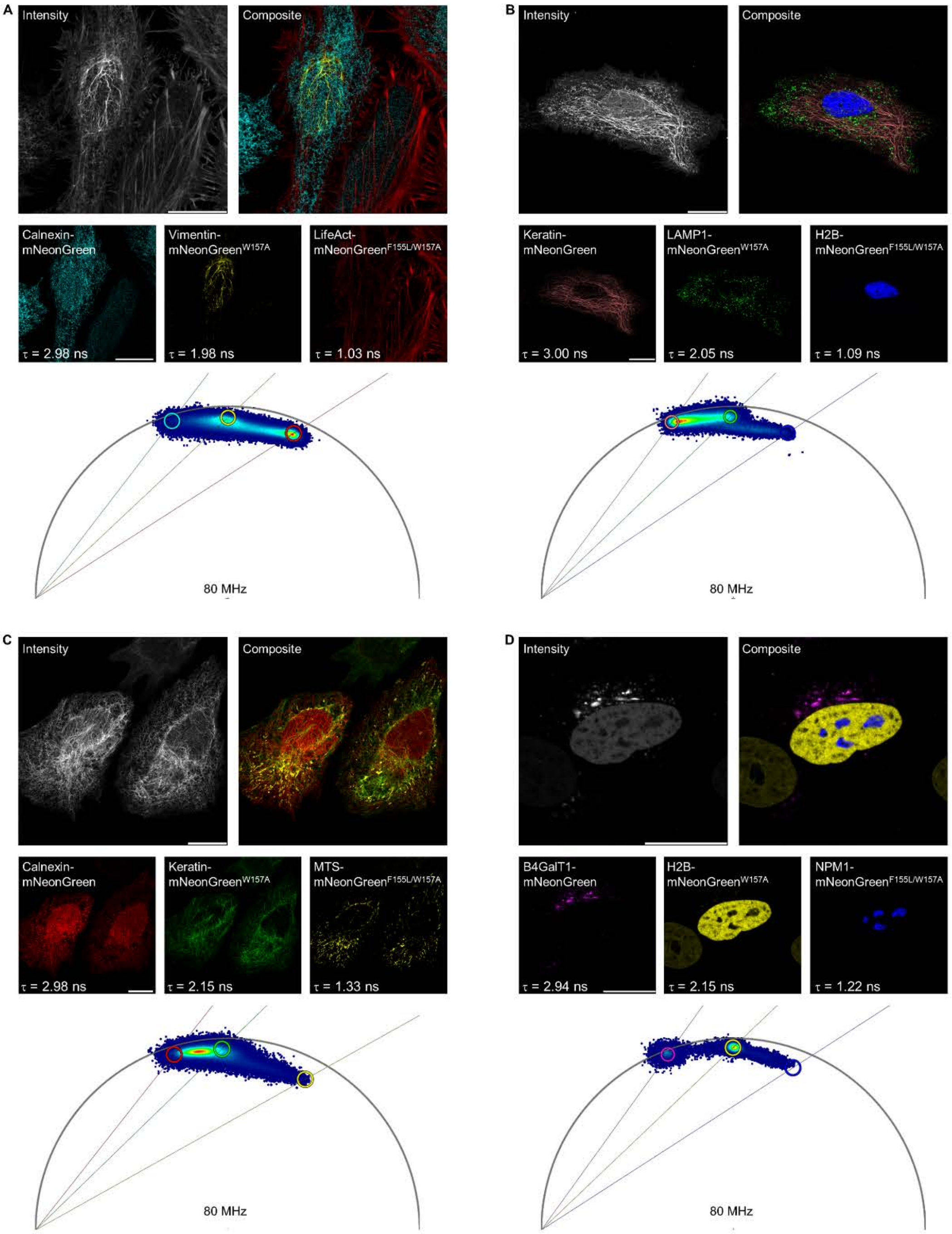


**Figure S28. Multiplexing imaging of three mNeonGreen TRFPs.**

(A) LifeAct-mNeonGreen$^{F155L/W157A}$ targeting actin filaments, Vimentin-mNeonGreen$^{W157A}$ targeting intermediate filaments and Calnexin-mNeonGreen targeting endoplasmic reticulum.

(B) H2B-mNeonGreen$^{F155L/W157A}$ targeting nucleus, LAMP1-mNeonGreen$^{W157A}$ targeting lysosomes and Keratin-mNeonGreen targeting intermediate filaments.

(C) MTS-mNeonGreen$^{F155L/W157A}$ targeting mitochondria, Keratin-mNeonGreen$^{W157A}$ targeting intermediate filaments and Calnexin-mNeonGreen targeting endoplasmic reticulum.

(D) NPM1-mNeonGreen$^{F155L/W157A}$ targeting nucleoli, H2B-mNeonGreen$^{W157A}$ targeting nucleus and B4GalT1-mNeonGreen targeting Golgi apparatus.

The fluorescence intensity, the composite, the three individual separated species, the fluorescence lifetime of each fusion proteins as well as the corresponding phasor plot used for separation are given. Scale bars, 20 µm.

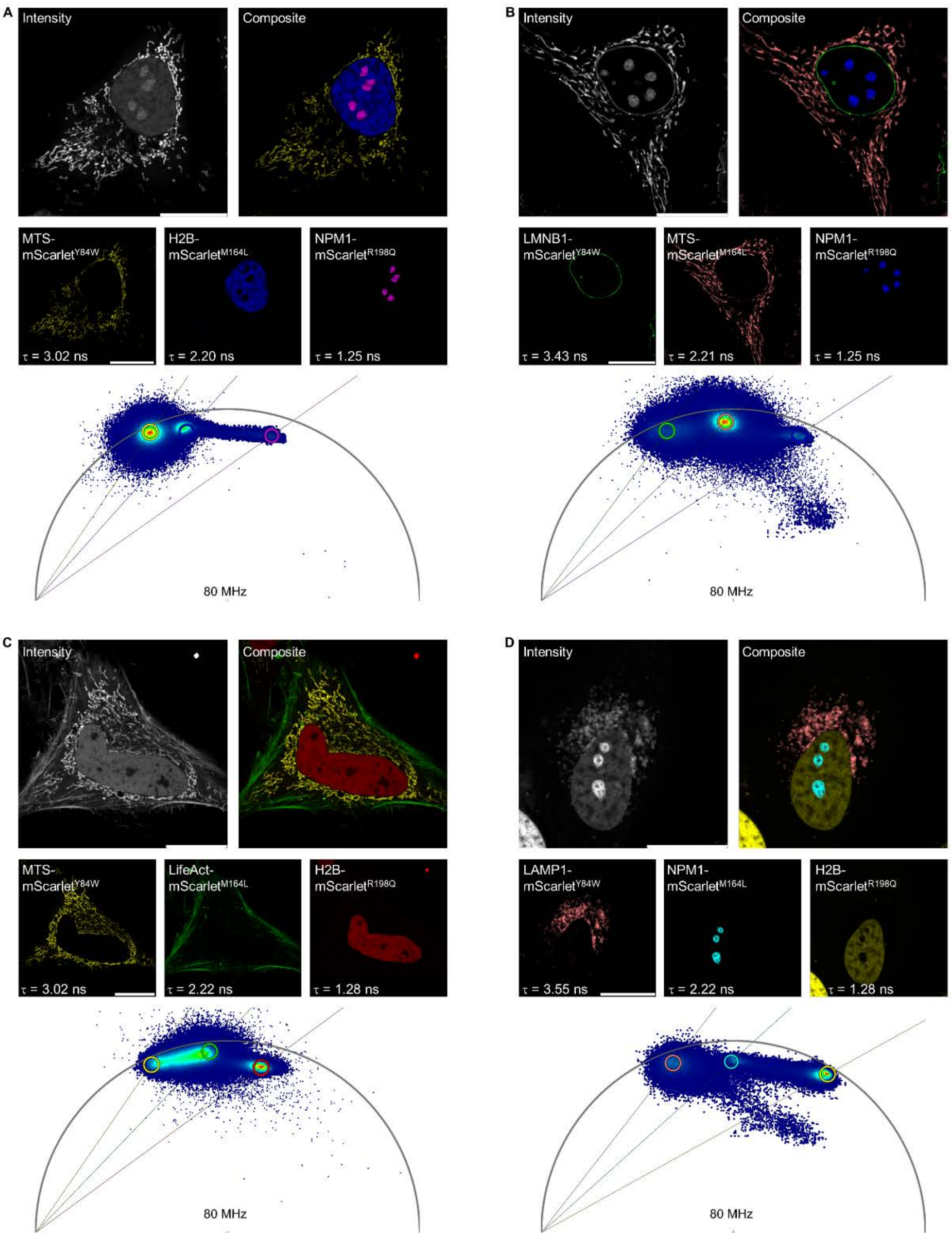


**Figure S29. Multiplexing imaging of three mScarlet TRFPs.**

(A) NPM1-mScarlet$^{R198Q}$ targeting nucleoli, H2B-mScarlet$^{M164L}$ targeting nucleus and MTS-mScarlet$^{Y84W}$ targeting mitochondria.

(B) NPM1-mScarlet$^{R198Q}$ targeting nucleoli, MTS-mScarlet$^{M164L}$ targeting mitochondria and LMNB1-mScarlet$^{Y84W}$ targeting nuclear membrane.

(C) H2B-mScarlet$^{R198Q}$ targeting nucleus, LifeAct-mScarlet$^{M164L}$ targeting actin filaments and MTS-mScarlet$^{Y84W}$ targeting mitochondria.

(D) H2B-mScarlet$^{R198Q}$ targeting nucleus, NPM1-mScarlet$^{M164L}$ targeting nucleoli and LAMP1-mScarlet$^{Y84W}$ targeting lysosomes.

The fluorescence intensity, the composite, the three individual separated species, the fluorescence lifetime of each fusion proteins as well as the corresponding phasor plot used for separation are given. Scale bars, 20 µm.

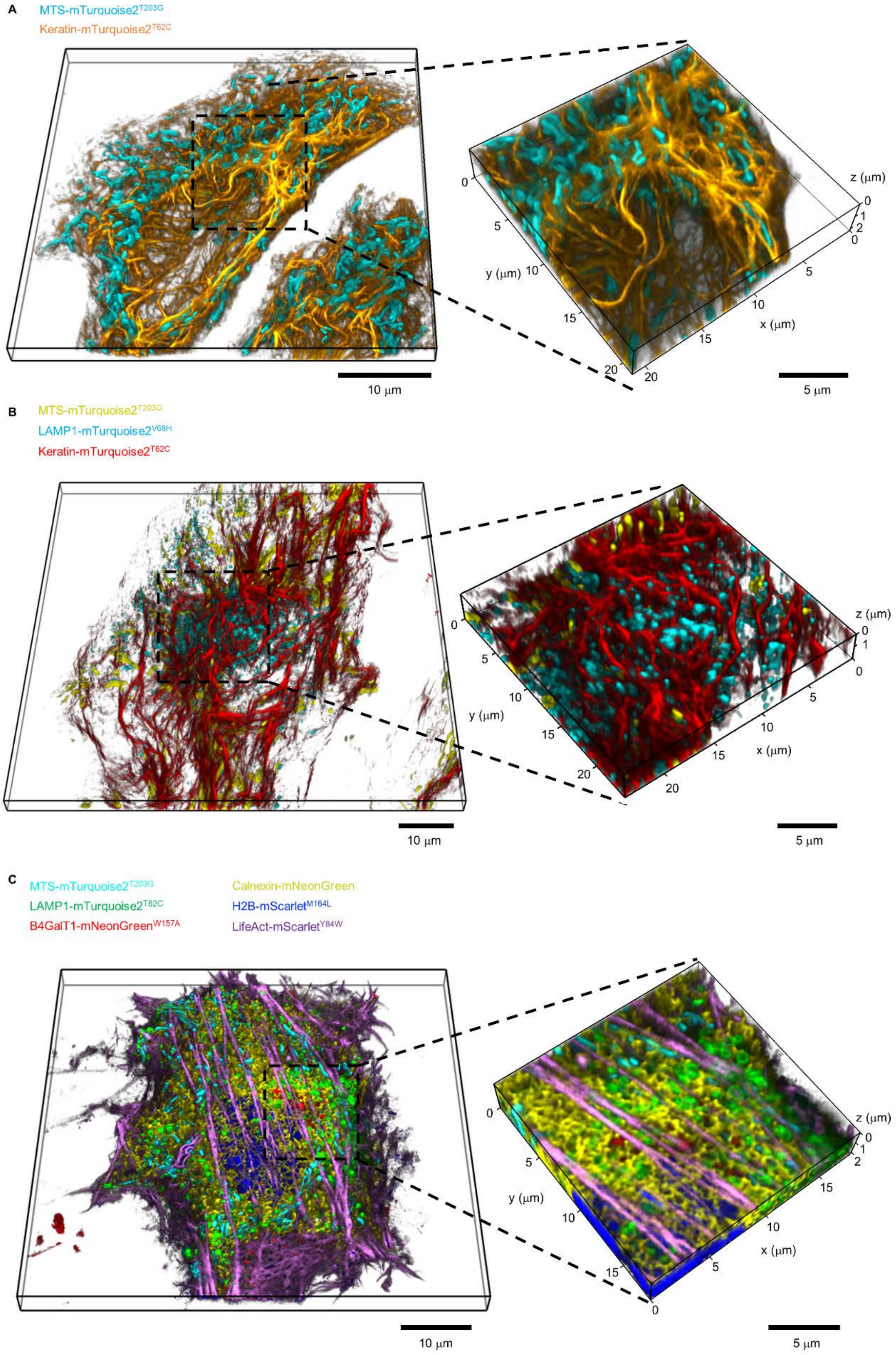


**Figure S30. Three dimensional images generated by TRFP multiplexing imaging.** (A) Z-stack images of MTS-mTurquoise2$^{T203G}$ targeting mitochondria and Keratin-mTurquoise2$^{T62C}$ targeting intermediate filaments.

(B) Z-stack images of MTS-mTurquoise2$^{T203G}$ targeting mitochondria, LAMP1-mTurquoise2$^{V68H}$ targeting lysosomes and Keratin-mTurquoise2$^{T62C}$ targeting intermediate filaments.
(C) Z-stack images of MTS-mTurquoise2$^{T203G}$ targeting mitochondria, B4GalT1-mTurquoise2$^{T62C}$ targeting Golgi apparatus, NPM1-mNeonGreen$^{F155L/W157A}$ targeting nucleoli, Calnexin-mNeonGreen targeting endoplasmic reticulum, H2B-mScarlet$^{M164L}$ targeting nucleus and LifeAct-mScarlet$^{Y84W}$ targeting actin filaments.
Species separation was performed via phasor analysis which has describe at methods. The images on the right are enlarged from the left images.

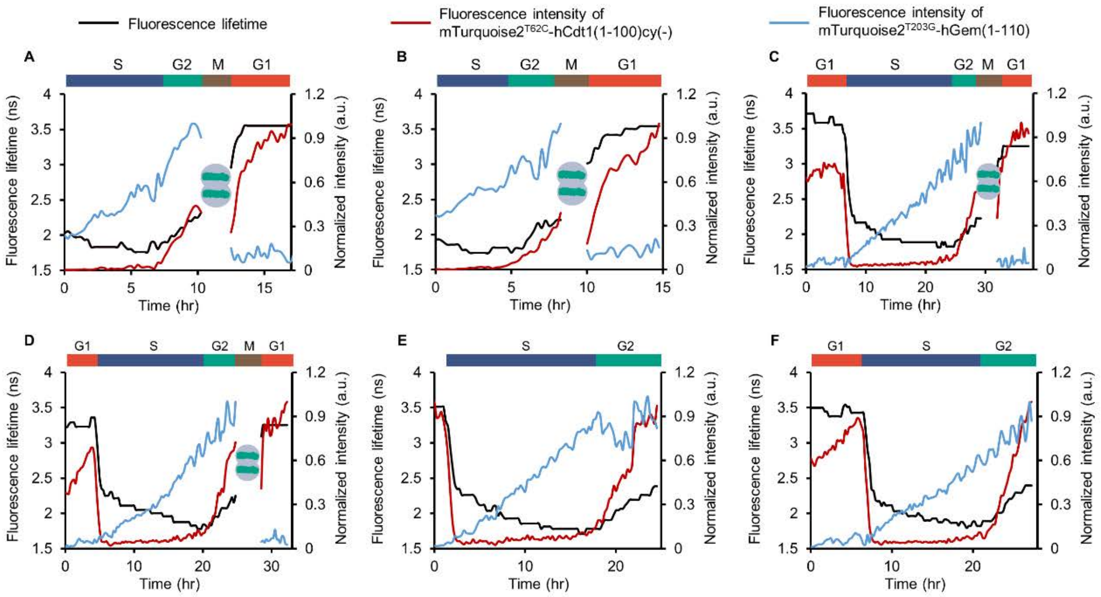


**Figure S31. Additional fluorescence lifetime and fluorescence intensity traces of mTurquoise2$^{T62C}$-hCdt1(1-100)cy(-) and mTurquoise2$^{T203G}$-hGem(1-110).** The coordinates on the left show the overall fluorescence lifetime (black line). The coordinates on the right show the intensity of mTurquoise2$^{T62C}$-hCdt1(1-100)cy(-) (red line) and mTurquoise2$^{T203G}$-hGem(1-110) (cyan line) after species separation. Fluorescence intensity was normalized to maximum value. Cell-cycle phases were assigned based on the principles in figure 5A.

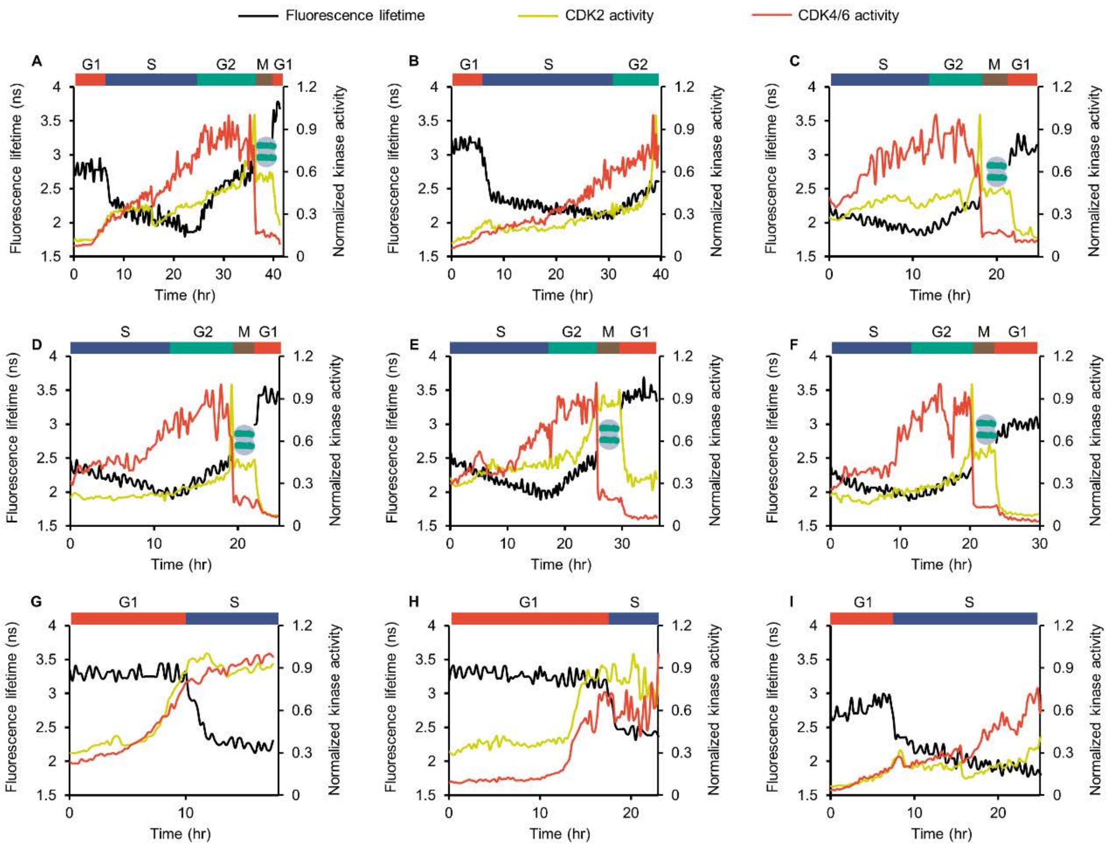


**Figure S32. Additional traces of cell cycle and the activity of CDK.** The coordinates on the left show the fluorescence lifetime (black line), which indicated the cell cycle. The coordinates on the right show the activity of CDK2 (yellow line) and CDK4/6 (red line). Kinase activity was calculated through the ratio of fluorescence intensity in the cytoplasm and nucleus and normalized to the maximum value. Cell-cycle phases were assigned based on the principles in figure 5A.

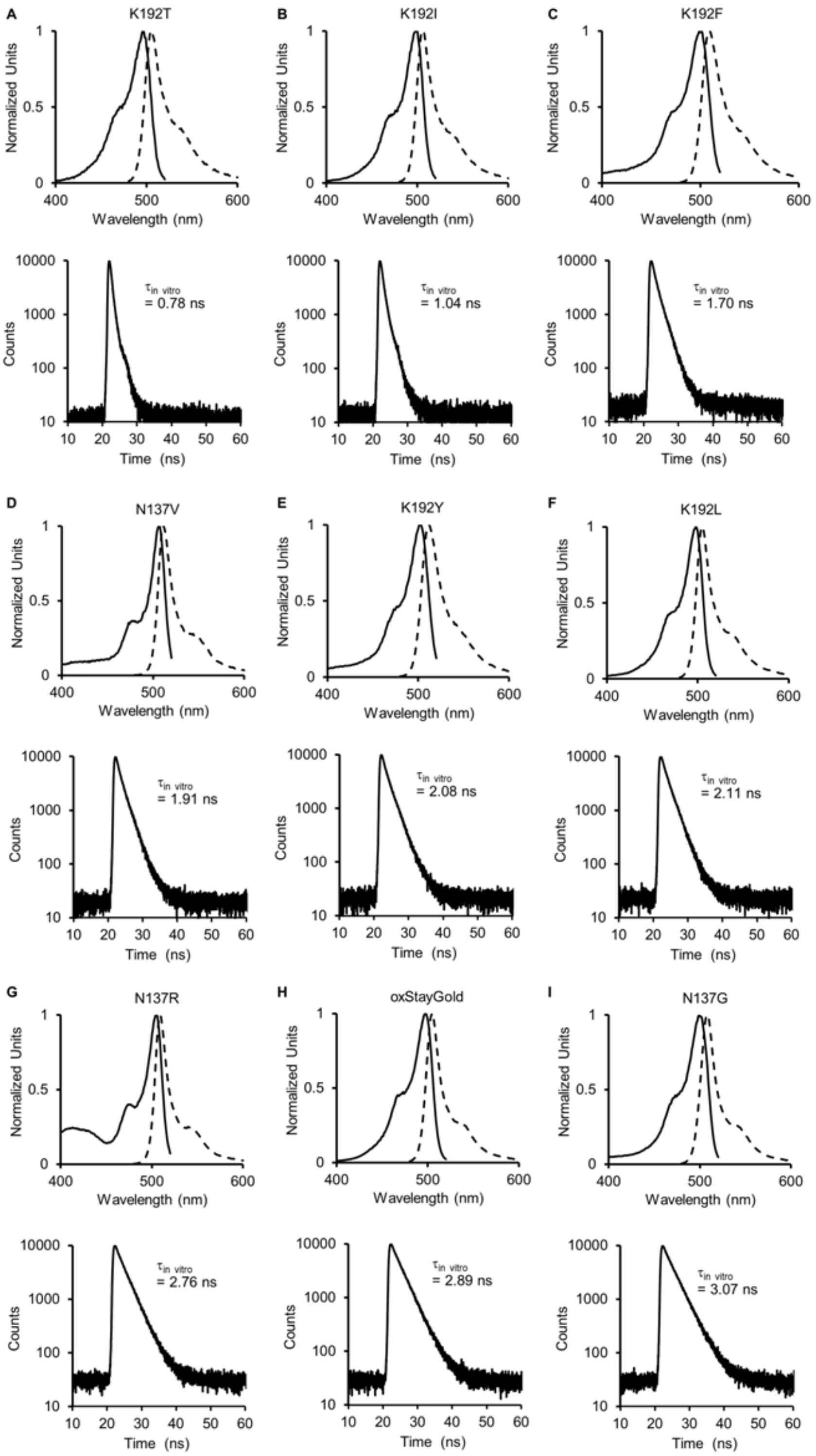


**Figure S33. Fluorescence spectra and TCSPC traces of oxStayGold TRFPs.**

Top panel: excitation (solid line) and emission (dash line) of oxSG$^{K192T}$ (A), oxSG$^{K192I}$ (B), oxSG$^{K192F}$ (C), oxSG$^{N137V}$ (D), oxSG$^{K192Y}$ (E), oxSG$^{K192L}$ (F), oxSG$^{N137R}$ (G), oxSG (H) and oxSG$^{N137G}$ (I) in the buffer (20 mM Tris, 500 mM NaCl, pH 7.5).

Bottom panel: decay of fluorescence lifetime of oxSG$^{K192T}$ (A), oxSG$^{K192I}$ (B), oxSG$^{K192F}$ (C), oxSG$^{N137V}$ (D), oxSG$^{K192Y}$ (E), oxSG$^{K192L}$ (F), oxSG$^{N137R}$ (G), oxSG (H) and oxSG$^{N137G}$ (I) in the buffer (20 mM Tris, 500 mM NaCl, pH 7.5). oxSG is the abbreviation of oxStayGold.

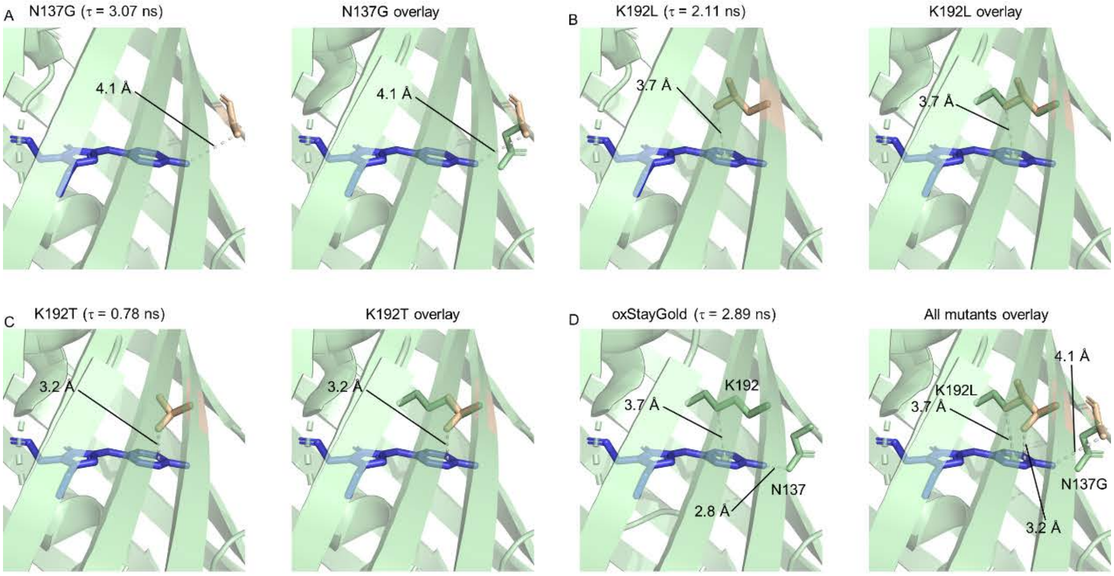


**Figure S34. Structural alignment of oxStayGold with its TRFP variants.**

(A-C) The structure of different variants (left panel) and overlay with oxStayGold (right panel).

(D) The structure of oxStayGold (left) and overlay with all variants (right). Green and Orange indicated the structure of oxStayGold and variants, respectively.

The structures of StayGold (PDB ID: 8bxt) was used as the template for the modeling of the variants through Rosetta.

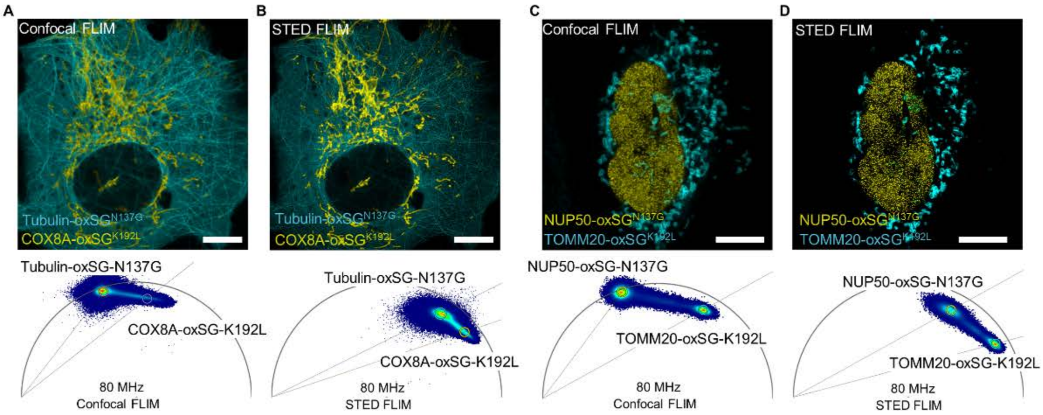


**Figure S35. STED-FLIM multiplexing imaging in living cells.**

(A and B) Phasor separation of live U-2 OS cells expressing tubulin-oxSG$^{N137G}$ and COX8A-oxSG$^{K192L}$ in confocal (A) and STED (B) mode.

(C and D) Phasor separation of live HeLa cells expressing NUP50-oxSG$^{N137G}$ and TOMM20-oxSG$^{K192L}$ in confocal (C) and STED (D) mode.

oxSG is the abbreviation of oxStayGold. Top panel and bottom panel indicate the composite and the phasor plot for separation. Scale bar, 10 μm.

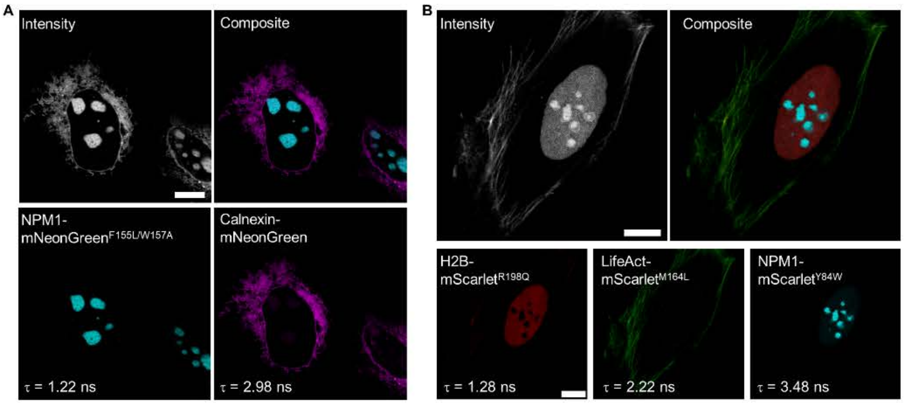


**Figure S36. FLIM multiplexing images acquired on a Leica STELLARIS 5 confocal microscope.**

(A) NPM1-mNeonGreen$^{F155L/W157A}$ targeting nucleoli and Calnexin-mNeonGreen targeting endoplasmic reticulum;

(B) H2B-mScarlet$^{R198Q}$ targeting nucleus, LifeAct-mScarlet$^{M164L}$ targeting actin filaments, and NPM1-mScarlet$^{Y84W}$ targeting nucleoli.

STELLARIS 5 equipped with a single frequency 80 MHz white light laser (WLL) and HyD S detectors. All the multiplexing images were acquired under TauSeparation mode, one of the standard counting acquisition modes. In TauSeparation mode, fluorescence signals of the same emission range could be collected based on specific fluorescence lifetime of choice. TRFPs were co-expressed in live U-2 OS cells. The fluorescence intensity, the composite, the individual separated species are showed. Scale bar, 10 μm.

| CFPs | $\lambda_{ex}$ (nm) | $\lambda_{em}$ (nm) | ε ($M^{-1}$ • $cm^{-1}$) | $\Phi_F$ | Brightness | $\tau_{in\ vitro}$ (ns) | $\tau_{in\ vivo}$ (ns) | $\tau_{in\ vitro}$-$\tau_{in\ vivo}$ (ns) |
|---|---|---|---|---|---|---|---|---|
| T203G | 436 | 479 | 31189 | 0.17 | 0.19 | 1.26±0.02 | 1.06±0.05 | 0.20 |
| T203L | 446 | 484 | 30616 | 0.26 | 0.28 | 1.99±0.01 | 1.64±0.01 | 0.35 |
| S205A | 436 | 481 | 27901 | 0.27 | 0.27 | 2.23±0.01 | 2.18±0.05 | 0.05 |
| D148Y | 436 | 479 | 28265 | 0.20 | 0.20 | 2.43±0.01 | 2.21±0.02 | 0.22 |
| S205P | 440 | 484 | 26456 | 0.21 | 0.20 | 2.16±0.01 | 2.21±0.07 | -0.05 |
| D148V | 440 | 478 | 31166 | 0.43 | 0.47 | 2.93±0.01 | 2.78±0.08 | 0.15 |
| V68H | 435 | 477 | 35439 | 0.57 | 0.71 | 3.14±0.00 | 2.84±0.03 | 0.30 |
| F146T | 437 | 478 | 30892 | 0.55 | 0.60 | 3.08±0.01 | 2.90±0.07 | 0.18 |
| F146Q | 439 | 479 | 29292 | 0.62 | 0.64 | 3.22±0.01 | 3.05±0.11 | 0.17 |
| F165L | 434 | 474 | 27823 | 0.66 | 0.65 | 3.53±0.02 | 3.36±0.02 | 0.17 |
| V68C | 435 | 476 | 30741 | 0.61 | 0.66 | 3.83±0.01 | 3.77±0.10 | 0.06 |
| mTurquoise2 | 435 | 474 | 30500 | 0.93 | 1.00 | 4.43±0.01 | 4.28±0.10 | 0.15 |
| T62P | 433 | 472 | 29111 | 0.98 | 1.01 | 4.66±0.01 | 4.38±0.01 | 0.28 |
| T62C | 437 | 477 | 35185 | 1.00 | 1.24 | 4.65±0.01 | 4.46±0.05 | 0.19 |

**Table S1. Photophysical parameters of mTurquoise2-based TRFPs.** Measurements were carried out in buffer containing 20 mM Tris, 500 mM NaCl, pH 7.5. The fluorescence lifetime in vivo was recorded using HEK293T cells expressing TRFPs. Fluorescence lifetime is the average value of three independent experiments. Standard deviations were derived from three independent experiments. The brightness of mTurquoise2 was set to 1.00.

| GFPs | $\lambda_{ex}$ (nm) | $\lambda_{em}$ (nm) | ε ($mol^{-1} \bullet cm^{-1}$) | $\Phi_F$ | Brightness | $\tau_{in\ vitro}$ (ns) | $\tau_{in\ vivo}$ (ns) | $\tau_{in\ vitro}$-$\tau_{in\ vivo}$ (ns) |
|---|---|---|---|---|---|---|---|---|
| R195A | 502 | 513 | 58996 | 0.05 | 0.03 | 1.12±0.01 | 0.83±0.06 | 0.29 |
| R195T | 498 | 512 | 52445 | 0.05 | 0.02 | 1.38±0.01 | 1.12±0.02 | 0.26 |
| F155L/W157A | 489 | 513 | 64626 | 0.21 | 0.13 | 1.68±0.03 | 1.34±0.08 | 0.34 |
| W157G | 500 | 515 | 73891 | 0.29 | 0.20 | 2.14±0.01 | 1.91±0.05 | 0.23 |
| W157A | 498 | 512 | 87814 | 0.41 | 0.34 | 2.29±0.01 | 2.14±0.07 | 0.15 |
| F155Y/W157A | 497 | 515 | 59645 | 0.30 | 0.17 | 1.95±0.00 | 2.19±0.03 | -0.24 |
| C139D | 500 | 514 | 58354 | 0.36 | 0.20 | 2.52±0.01 | 2.52±0.03 | 0.00 |
| A171M | 506 | 517 | 129859 | 0.62 | 0.75 | 2.94±0.01 | 2.80±0.07 | 0.14 |
| mNeonGreen | 506 | 517 | 133830 | 0.80 | 1.00 | 3.15±0.01 | 3.05±0.02 | 0.10 |

**Table S2. Photophysical parameters of mNeonGreen-based TRFPs.** Measurements were carried out in buffer containing 20 mM Tris, 500 mM NaCl, pH 7.5. The fluorescence lifetime in vivo was recorded using HEK293T cells expressing TRFPs. Fluorescence lifetime is the average value of three independent experiments. Standard deviations were derived from three independent experiments. The brightness of mNeonGreen was set to 1.00.

| RFPs | $\lambda_{ex}$ (nm) | $\lambda_{em}$ (nm) | ε ($M^{-1} \bullet cm^{-1}$) | $\Phi_F$ | Brightness | $\tau_{in\ vitro}$ (ns) | $\tau_{in\ vivo}$ (ns) | $\tau_{in\ vitro}$-$\tau_{in\ vivo}$ (ns) |
|---|---|---|---|---|---|---|---|---|
| R198T | 594 | 615 | 102307 | 0.16 | 0.21 | 1.16±0.01 | 1.26±0.01 | -0.10 |
| R198Q | 593 | 609 | 105115 | 0.23 | 0.30 | 1.41±0.01 | 1.37±0.00 | 0.04 |
| M164H | 551 | 591 | 81893 | 0.24 | 0.25 | 1.58±0.01 | 1.38±0.06 | 0.20 |
| M164S | 561 | 598 | 56450 | 0.28 | 0.20 | 1.82±0.00 | 1.77±0.03 | 0.05 |
| M164A | 555 | 595 | 75433 | 0.28 | 0.27 | 1.78±0.01 | 1.82±0.07 | -0.04 |
| M164V | 570 | 594 | 77252 | 0.3 | 0.29 | 1.95±0.01 | 1.82±0.02 | 0.13 |
| M164L | 569 | 593 | 81669 | 0.32 | 0.33 | 2.13±0.00 | 2.42±0.01 | -0.29 |
| F178A | 564 | 592 | 89325 | 0.51 | 0.57 | 3.21±0.01 | 3.17±0.04 | 0.04 |
| mScarlet | 569 | 593 | 113766 | 0.70 | 1.00 | 3.62±0.00 | 3.62±0.00 | 0.00 |
| Y84W | 569 | 593 | 116263 | 0.73 | 1.07 | 3.68±0.01 | 3.68±0.02 | 0.00 |

**Table S3. Photophysical parameters of mScarlet-based TRFPs.** Measurements were carried out in buffer containing 20 mM Tris, 500 mM NaCl, pH 7.5. The fluorescence lifetime in vivo was recorded using HEK293T cells expressing TRFPs. Fluorescence lifetime is the average value of three independent experiments. Standard deviations were derived from three independent experiments. The brightness of mScarlet was set to 1.00.

| BFPs | $\lambda_{ex}$ (nm) | $\lambda_{em}$ (nm) | ε ($M^{-1}$ • $cm^{-1}$) | $\Phi_F$ | Brightness | $\tau_{in\ vitro}$ (ns) |
|---|---|---|---|---|---|---|
| E222D | 383 | 451 | 33576 | 0.21 | 0.39 | 2.16±0.01 |
| E222T | 379 | 449 | 32077 | 0.29 | 0.52 | 2.40±0.01 |
| I150D | 377 | 447 | 29140 | 0.45 | 0.73 | 2.57±0.01 |
| H148A | 383 | 454 | 29283 | 0.44 | 0.72 | 2.85±0.00 |
| EBFP2 | 383 | 451 | 32000 | 0.56 | 1.00 | 3.45±0.00 |
| T62V | 385 | 451 | 29062 | 0.52 | 0.84 | 3.89±0.00 |

**Table S4. Photophysical parameters of EBFP2-based TRFPs.** Measurements were carried out in buffer containing 20 mM Tris, 500 mM NaCl, pH 7.5. The fluorescence lifetime in vivo was recorded using HEK293T cells expressing TRFPs. Fluorescence lifetime is the average value of three independent experiments. Standard deviations were derived from three independent experiments. The brightness of EBFP2 was set to 1.00.

| YFPs | $\lambda_{ex}$ (nm) | $\lambda_{em}$ (nm) | $\varepsilon$ ($M^{-1} \bullet cm^{-1}$) | $\Phi_F$ | Brightness | $\tau_{in\ vitro}$ (ns) |
|---|---|---|---|---|---|---|
| S205A | 516 | 529 | 95430 | 0.42 | 0.60 | 2.36±0.01 |
| S205V | 517 | 528 | 44269 | 0.37 | 0.25 | 2.37±0.00 |
| H148V | 516 | 526 | 103331 | 0.48 | 0.75 | 2.70±0.02 |
| H148E | 516 | 526 | 56710 | 0.49 | 0.42 | 2.72±0.00 |
| H148A | 517 | 528 | 49773 | 0.54 | 0.40 | 2.74±0.01 |
| mVenus | 516 | 528 | 104000 | 0.64 | 1.00 | 3.21±0.00 |

**Table S5. Photophysical parameters of mVenus-TRFPs.** Measurements were carried out in buffer containing 20 mM Tris, 500 mM NaCl, pH 7.5. The fluorescence lifetime in vivo was recorded using HEK293T cells expressing TRFPs. Fluorescence lifetime is the average value of three independent experiments. Standard deviations were derived from three independent experiments. The brightness of mVenus was set to 1.00.

| OFPs | $\lambda_{ex}$ (nm) | $\lambda_{em}$ (nm) | $\varepsilon$ ($M^{-1} \bullet cm^{-1}$) | $\Phi_F$ | Brightness | $\tau_{in\ vitro}$ (ns) |
|---|---|---|---|---|---|---|
| M163A | 544 | 560 | 78616 | 0.15 | 0.24 | 1.74±0.01 |
| S146N | 551 | 564 | 73997 | 0.23 | 0.35 | 2.09±0.00 |
| I197Y | 551 | 569 | 45676 | 0.19 | 0.18 | 2.12±0.01 |
| I161M | 549 | 564 | 66540 | 0.47 | 0.64 | 2.89±0.00 |
| mOrange | 548 | 562 | 71000 | 0.69 | 1.00 | 3.46±0.00 |
| I161L | 549 | 563 | 97911 | 0.89 | 1.78 | 3.60±0.00 |

**Table S6. Photophysical parameters of mOrange-based TRFPs.** Measurements were carried out in buffer containing 20 mM Tris, 500 mM NaCl, pH 7.5. The fluorescence lifetime in vivo was recorded using HEK293T cells expressing TRFPs. Fluorescence lifetime is the average value of three independent experiments. Standard deviations were derived from three independent experiments. The brightness of mOrange was set to 1.00.

| Far-red FPs | $\lambda_{ex}$ (nm) | $\lambda_{em}$ (nm) | ε ($M^{-1}$ • $cm^{-1}$) | $\Phi_F$ | Brightness | $\tau_{in\ vitro}$ (ns) |
|---|---|---|---|---|---|---|
| S144T | 586 | 623 | 53447 | 0.29 | 0.62 | 2.16±0.00 |
| L175G | 564 | 624 | 32941 | 0.17 | 0.22 | 2.29±0.01 |
| A159N | 583 | 629 | 40545 | 0.34 | 0.5 | 2.34±0.02 |
| A159C | 589 | 632 | 39007 | 0.37 | 0.58 | 2.45±0.01 |
| mKate2 | 587 | 627 | 62500 | 0.40 | 1.00 | 2.60±0.00 |
| T61Q | 585 | 631 | 35827 | 0.45 | 0.64 | 2.91±0.01 |

**Table S7. Photophysical parameters of mKate2-based TRFPs.** Measurements were carried out in buffer containing 20 mM Tris, 500 mM NaCl, pH 7.5. The fluorescence lifetime in vivo was recorded using HEK293T cells expressing TRFPs. Fluorescence lifetime is the average value of three independent experiments. Standard deviations were derived from three independent experiments. The brightness of mKate2 was set to 1.00.

| TRFPs | $\lambda_{ex}$ (nm) | $\lambda_{em}$ (nm) | $\varepsilon$ ($M^{-1} \bullet cm^{-1}$) | $\Phi_F$ | Brightness | $\tau_{in\ vitro}$ (ns) |
|---|---|---|---|---|---|---|
| EBFP2$^{E222D}$ | 383 | 451 | 33576 | 0.21 | 0.06 | 2.16±0.01 |
| EBFP2$^{H148A}$ | 383 | 454 | 29283 | 0.44 | 0.12 | 2.85±0.00 |
| EBFP2 | 383 | 451 | 32000 | 0.56 | 0.17 | 3.45±0.00 |
| EBFP2$^{T62V}$ | 385 | 451 | 29062 | 0.52 | 0.14 | 3.89±0.00 |
| mTurq2$^{T203G}$ | 436 | 479 | 31189 | 0.17 | 0.05 | 1.26±0.02 |
| mTurq2$^{T203L}$ | 446 | 484 | 30616 | 0.26 | 0.07 | 1.99±0.01 |
| mTurq2$^{D148Y}$ | 436 | 479 | 28265 | 0.20 | 0.05 | 2.43±0.01 |
| mTurq2$^{V68H}$ | 435 | 477 | 35439 | 0.57 | 0.19 | 3.14±0.00 |
| mTurq2$^{F165L}$ | 434 | 474 | 27823 | 0.66 | 0.17 | 3.53±0.02 |
| mTurq2$^{V68C}$ | 435 | 476 | 30741 | 0.61 | 0.66 | 3.83±0.01 |
| mTurq2$^{T62C}$ | 437 | 477 | 35185 | 1.00 | 0.33 | 4.65±0.01 |
| mNG$^{R195A}$ | 502 | 513 | 58996 | 0.05 | 0.03 | 1.12±0.01 |
| mNG$^{F155L/W157A}$ | 489 | 513 | 64626 | 0.21 | 0.13 | 1.68±0.03 |
| mNG$^{W157A}$ | 498 | 512 | 87814 | 0.41 | 0.34 | 2.29±0.01 |
| mNG | 506 | 517 | 133830 | 0.80 | 1.00 | 3.15±0.01 |
| mVenus$^{S205A}$ | 516 | 529 | 95430 | 0.42 | 0.37 | 2.36±0.01 |
| mVenus$^{H148A}$ | 517 | 528 | 49773 | 0.54 | 0.25 | 2.74±0.01 |
| mVenus | 516 | 528 | 104000 | 0.64 | 0.62 | 3.21±0.00 |
| mOrange$^{M163A}$ | 544 | 560 | 78616 | 0.15 | 0.11 | 1.74±0.01 |
| mOrange$^{I197Y}$ | 551 | 569 | 45676 | 0.19 | 0.08 | 2.12±0.01 |
| mOrange$^{I161M}$ | 549 | 564 | 66540 | 0.47 | 0.29 | 2.89±0.00 |
| mOrange$^{I161L}$ | 549 | 563 | 97911 | 0.89 | 0.81 | 3.60±0.00 |
| mScarlet$^{R198Q}$ | 593 | 609 | 105115 | 0.23 | 0.23 | 1.41±0.01 |
| mScarlet$^{M164L}$ | 569 | 593 | 81669 | 0.32 | 0.24 | 2.13±0.00 |
| mScarlet$^{F178A}$ | 564 | 592 | 89325 | 0.51 | 0.43 | 3.21±0.01 |
| mScarlet$^{Y84W}$ | 569 | 593 | 116263 | 0.73 | 0.79 | 3.68±0.01 |
| mKate2$^{S144T}$ | 586 | 623 | 53447 | 0.29 | 0.62 | 2.16±0.00 |
| mKate2 | 587 | 627 | 62500 | 0.40 | 0.23 | 2.60±0.00 |
| mKate2$^{T61Q}$ | 585 | 631 | 35827 | 0.45 | 0.15 | 2.91±0.01 |

**Table S8. Photophysical parameters of TRFPs recommended for the temporal and spectral resolved multiplexing imaging.** Measurements were carried out in buffer containing 20 mM Tris, 500 mM NaCl, pH 7.5. The fluorescence lifetime in vivo was recorded using HEK293T cells expressing TRFPs. Fluorescence lifetime is the average value of three independent experiments. Standard deviations were derived from three independent experiments. The brightness of mNeonGreen was set to 1.00. mTurq2 and mNG are the abbreviation of mTurquoise2 and mNeonGreen, respectively.

| GFPs | $\lambda_{ex}$ (nm) | $\lambda_{em}$ (nm) | $\varepsilon$ ($M^{-1} \bullet cm^{-1}$) | $\Phi_F$ | Brightness | $\tau_{in\ vitro}$ (ns) |
|---|---|---|---|---|---|---|
| K192T | 496 | 505 | 173217 | 0.13 | 0.12 | 0.78±0.00 |
| K192I | 498 | 507 | 191601 | 0.20 | 0.21 | 1.04±0.00 |
| K192F | 499 | 509 | 142916 | 0.36 | 0.28 | 1.70±0.00 |
| N137V | 506 | 511 | 143423 | 0.43 | 0.34 | 1.91±0.01 |
| K192Y | 502 | 512 | 118213 | 0.51 | 0.33 | 2.08±0.00 |
| K192L | 498 | 505 | 188955 | 0.40 | 0.41 | 2.11±0.00 |
| N137R | 505 | 509 | 120843 | 0.75 | 0.50 | 2.76±0.02 |
| oxStayGold | 497 | 505 | 196397 | 0.93 | 1.00 | 2.89±0.01 |
| N137G | 499 | 508 | 143381 | 0.85 | 0.67 | 3.07±0.00 |

**Table S9. Comparisons of biochemical and biophysical characteristics of oxStayGold and its variants.** Measurements were carried out in buffer containing 20 mM Tris, 500 mM NaCl, pH 7.5. The fluorescence lifetime in vivo was recorded using HEK293T cells expressing TRFPs. Fluorescence lifetime is the average value of three independent experiments. Standard deviations were derived from three independent experiments. The brightness of oxStayGold was set to 1.00.
